# FoxA1 and FoxA2 regulate growth and cellular identity in NKX2-1-positive lung adenocarcinoma

**DOI:** 10.1101/2021.06.29.450247

**Authors:** Grace Orstad, Gabriela Fort, Timothy J Parnell, Alex Jones, Chris Stubben, Brian Lohman, Katherine L. Gillis, Walter Orellana, Rushmeen Tariq, Olaf Klingbeil, Klaus Kaestner, Christopher R. Vakoc, Benjamin T. Spike, Eric L. Snyder

**Affiliations:** Huntsman Cancer Institute, University of Utah, Salt Lake City, Utah, USA; Department of Oncological Sciences, University of Utah, Salt Lake City, Utah, USA; Bioinformatics Shared Resource, Huntsman Cancer Institute, University of Utah, Salt Lake City, Utah, USA; Department of Pathology, University of Utah, Salt Lake City, Utah, USA; HCI Clinical Trials Operations, Huntsman Cancer Institute, University of Utah, Salt Lake City, Utah, USA; Cold Spring Harbor Laboratory, Cold Spring Harbor, New York, USA; Department of Genetics, Perelman School of Medicine, University of Pennsylvania, Philadelphia, Pennsylvania, USA

**Keywords:** FoxA1, FoxA2, NKX2-1, lung adenocarcinoma, cellular identity

## Abstract

Changes in cellular identity (also known as histologic transformation or lineage plasticity) can drive malignant progression and resistance to therapy in multiple types of cancer, including lung adenocarcinoma (LUAD). The lineage specifying transcription factors FoxA1 and FoxA2 (FoxA1/2) regulate identity in NKX2-1/TTF1-negative LUAD. However, their role in NKX2-1-positive LUAD has not been systematically investigated. We find that *Foxa1/2* knockout severely impairs tumorigenesis in KRAS-driven genetically engineered mouse models and human cell lines. *Foxa1/2* deletion leads to the collapse of a dual identity state, marked by co-expression of pulmonary and gastrointestinal transcriptional programs, which has been implicated in LUAD progression. Mechanistically, loss of FoxA1/2 leads to aberrant NKX2-1 activity and genomic localization, which inhibits tumorigenesis and drives alternative cellular identity programs associated with non-proliferative states. This work demonstrates that FoxA1/2 expression is a novel lineage-specific vulnerability in NKX2-1-positive LUAD and identifies mechanisms of response and resistance to targeting FoxA1/2 in this disease.

## INTRODUCTION

The behavior of a cancer cell is initially constrained by the specific epigenetic state of its cell of origin. Changes in cancer cell identity (often referred to as lineage switching, lineage plasticity, or histological transformation) have become increasingly well characterized as a mechanism of cancer progression and acquired resistance to targeted therapies (Boumahdi and de Sauvage, 2020; Le Magnen et al., 2018; Quintanal-Villalonga et al., 2020). Lung cancer is the leading cause of cancer death worldwide, and lung adenocarcinoma (LUAD) is the most frequently diagnosed subtype of this disease (Meza et al., 2015). In patients, LUAD exhibits significant heterogeneity in cell identity and differentiation state (Travis et al., 2011); these characteristics correlate directly with prognosis, response to available therapies, and acquisition of drug resistance (Campos-Parra et al., 2014; Niederst et al., 2015; Russell, 2011; Sun and Yang, 2006). Although epigenetic changes have been implicated in cell identity heterogeneity, the field lacks a comprehensive understanding of factors governing LUAD cell identity and how perturbations within these networks alter malignant potential (LaFave et al., 2020; Marjanovic et al., 2020; Tavernari et al., 2021).

Previous work from our lab and others has shown that the transcription factors FoxA1, FoxA2, and NKX2-1/TTF1 coordinately regulate cellular identity in LUAD (Camolotto et al., 2018; Li et al., 2015; Snyder et al., 2013; Watanabe et al., 2013; Winslow et al., 2011). The homeodomain transcription factor NKX2-1/TTF1 is a master regulator of normal lung morphogenesis and differentiation (Bingle, 1997; Bohinski et al., 1994; Little et al., 2019; Little et al., 2021; Liu et al., 2002; Yi et al., 2002) as well as LUAD identity(Snyder *et al*., 2013; Watanabe *et al*., 2013; Winslow *et al*., 2011). NKX2-1 activity is dependent on differential interactions with many transcription factors, co-activators and co-repressors in a gene-specific manner, including FoxA1 and FoxA2 in both normal and malignant lung tissue. FoxA1/2 physically interact with NKX2-1, which enhances the expression of NKX2-1 target genes that have both NKX2-1 and FoxA binding motifs within the promoter (Minoo et al., 2007). FoxA1 and FoxA2 (FoxA1/2) are closely related paralogs in the forkhead box transcription factor family (Kaestner, 2010). They share many cellular roles including the establishment and maintenance of cellular identity in multiple endodermal-derived tissues (Bochkis et al., 2012). FoxA1/2 are critical regulators of normal foregut differentiation; embryonic deletion of *Foxa1/2* in the foregut endoderm blocks branching morphogenesis of the lung (Wan et al., 2005), hepatic specification (Kaestner, 2005; Lee et al., 2005), and pancreatogenesis (Gao et al., 2008). In the adult mouse lung, FoxA1/2 and NKX2-1 are co-expressed in the tracheal epithelium, bronchiolar epithelium, and Type 2 Alveolar (AT2) cells (Besnard et al., 2004; Paranjapye et al., 2020).

FoxA1 and FoxA2 play context-specific roles in multiple types of cancer (Gao et al., 2020). For example, FOXA1 functions as an oncogene in breast and prostate cancer by virtue of genomic amplifications and point mutations within the forkhead box DNA binding domain (Bernardo and Keri, 2012; Robinson et al., 2013; Teng et al., 2021). In a p53-proficient LUAD genetically engineered mouse model (GEMM), we have previously shown that NKX2-1 regulates FoxA1/2 function by dictating whether they bind pulmonary or gastric marker genes (Snyder *et al*., 2013). In the absence of NKX2-1, FoxA1/2 drive a gastric differentiation program, reflecting their role in normal foregut differentiation. *Nkx2-1*-deleted murine tumors closely resemble a subtype of human LUAD termed invasive mucinous adenocarcinoma (IMA), which also exhibits gastric differentiation (Cha and Shim, 2017). In contrast, FoxA1/2 bind to regulatory elements of pulmonary marker genes in both human (Boggaram, 2009; Watanabe *et al*., 2013) and murine (Snyder *et al*., 2013) NKX2-1-positive LUAD. FoxA1/2 and NKX2-1 exhibit extensive genomic co-localization in LUAD, suggesting they may have cooperative functions in cancer that mirror their functions in normal tissue. *NKX2-1* is genomically amplified in ∼12% of LUAD (Weir et al., 2007), and *FOXA1* is frequently co-amplified with *NKX2-1* (∼10% of human LUAD cases in TCGA, PanCancer Atlas, data not shown). In a functional screen of nuclear receptors and their co-regulators, FOXA1 was identified as pro-tumorigenic in a subset of human LUAD cell lines analyzed. Those with the highest *FOXA1* copy number or expression exhibit varied levels of dependence on the gene, whereas cell lines with low levels were generally *FOXA1*-independent (Hight et al., 2020). In contrast, a subset of human LUAD downregulate FOXA2, but FOXA1 and NKX2-1 levels were not evaluated in this study (Basseres et al., 2012).

Despite these observations, FoxA1 and FoxA2 have not been systematically characterized in NKX2-1-positive LUAD, which comprises ∼75% of human LUAD cases overall (Bejarano et al., 1996). Here we use complementary GEMMs, organoid cultures, and human cell lines to demonstrate that FoxA1/2 are critical regulators of growth and cellular identity in NKX2-1-positive LUAD.

## RESULTS

### FoxA1/2 are required for growth of NKX2-1-positive lung adenocarcinoma

We first evaluated expression of NKX2-1, FoxA1 and FoxA2 in primary human LUAD via immunohistochemistry (IHC) on whole tumor sections and tumor microarrays (n=132 independent tumors). We found that all NKX2-1 positive tumors evaluated (n=87) express FoxA1 and/or FoxA2. In contrast, 17% (8/45) of NKX2-1-negative tumors have no detectable FoxA1 or FoxA2 expression (Supplemental Figure 1A-B, p<0.0001, Fisher’s Exact test). These initial observations reveal that NKX2-1 and FoxA1/2 are co-expressed in human LUAD. NKX2-1 and FoxA1/2 are also co-expressed in the LUAD GEMM driven by KRAS^G12D^ and deletion of *Trp53* (“KP” hereafter; Figure 1A, left panels). These data suggest that there is selective pressure for NKX2-1 positive LUAD tumors to retain FoxA1/2, and that NKX2-1-positive disease may be dependent on FoxA1/2 activity. Consistent with this possibility, stochastic NKX2-1 loss typically precedes FoxA2 loss in the KP model (Li *et al*., 2015).

**Figure 1:**
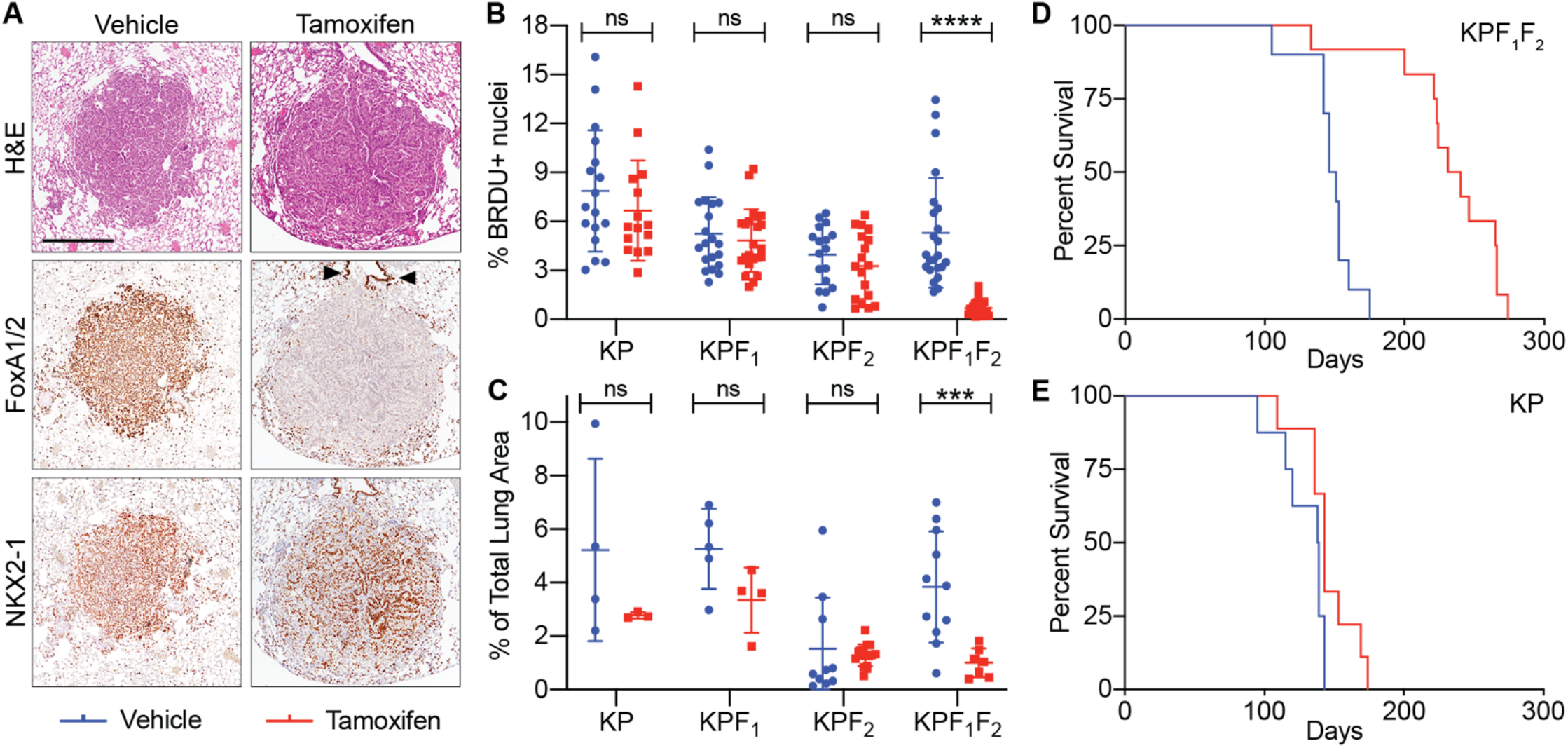
FoxA1/2 are required for growth of NKX2-1-positive lung adenocarcinoma in vivo. A) Representative images of KPF_1_F_2_ tumors (i.e. in *Kras^FSF-G12D/+^*; *Trp53^Frt/Frt^*; *Rosa26^FSF-CreERT2^; Foxa1^F/F^*; *Foxa2^F/F^* mice) 8 weeks post-initiation with adenoviral CMV-FlpO. Tamoxifen or vehicle was administered 6 weeks post-initiation. IHC for FoxA1/2 and NKX2-1 and Hematoxylin and Eosin (H&E) staining shown. Arrowheads in center right panel mark FoxA1/2-positive normal bronchiolar epithelium, confirming tumor-specific loss of FoxA1/2 (scale bar 1000 μm). B) BRDU IHC quantification of individual KP, KPF_1_, KPF_2_, and KPF_1_F_2_ tumors initiated with adenoviral CMV-FlpO, injected with tamoxifen (red) or vehicle (blue) 6 weeks later, and sacrificed at 8 weeks (1 hour post BRDU injection; ****p<0.0001, unpaired t-test) C) Tumor burden quantification of individual mice from the experiment shown in B (***p=0.001014, unpaired t-test) D) KPF_1_F_2_ survival study in which tamoxifen (red, n=12) or vehicle (blue, n=10) was administered 10 weeks post-initiation with lentiviral EFS-FlpO (vehicle median survival of 21.21 weeks; tamoxifen median survival of 34.71 weeks; p<0.0001, Log-rank Mantel-Cox test). E) KP survival study in which tamoxifen (red, n=9) or vehicle (blue, n=8) was administered 10 weeks post-initiation with lentiviral EFS-FlpO (vehicle median survival of 19.79 weeks; tamoxifen median survival of 20.43 weeks; p=0.0645, Log-rank Mantel-Cox test).

To investigate the role of FoxA1/2 in NKX2-1-positive LUAD, we developed a sequential recombination system to delete *Foxa1* and/or *Foxa2* in existing KRAS driven, NKX2-1 positive lung tumors in vivo. We generated *Kras^FSF-G12D/+^*; *Trp53^Frt/Frt^*; *Rosa26^FSF-CreERT2^; Foxa1^F/F^*; *Foxa2^F/F^* mice, as well as control mice harboring conditional alleles of either *Foxa1* (Gao et al., 2008) or *Foxa2* (Sund et al., 2000) alone. In these mice, delivery of the FlpO recombinase to the lung epithelium activates oncogenic *Kras^G12D^* (Young et al., 2011), inactivates the tumor suppressor *Trp53* (Lee et al., 2012), and induces transcription of Cre^ERT2^ from the *Rosa26* locus (Schonhuber et al., 2014). Upon tamoxifen treatment, Cre^ERT2^ enters the nucleus and deletes *Foxa1* and/or *Foxa2* (KPF_1_, KPF_2_, or KPF_1_F_2_ hereafter) specifically in LUAD cells (Supplemental Figure 1C-D). We first used this model system to determine the effect of *Foxa1/2* deletion at an early timepoint in tumor development. Lung tumors were initiated and allowed to develop for 6 weeks before deletion of *Foxa1/2* via tamoxifen treatment. Eight weeks after tumor initiation, lung tissue was collected for histopathologic evaluation (Figure 1A). Heterozygous deletion of a single copy of *Foxa1* and *Foxa2* (KPF_1_^F/+^F_2_^F/+^; shortened to KP hereafter), as well as homozygous deletion of either *Foxa1* (KPF_1_) or *Foxa2* (KPF_2_) alone had no effect on tumor proliferation (Figure 1B) or overall tumor burden (Figure 1C). Simultaneous deletion of both *Foxa1* and *Foxa2* caused a significant decrease in both proliferation (7.8-fold decrease) and overall tumor burden at an early timepoint (Figure 1B-C), demonstrating that low-grade LUAD tumors are dependent on FoxA1/2. There was no increase in levels of the apoptosis marker cleaved caspase-3 (CC3) at this timepoint (Supplemental Figure 1E). Concomitant *Foxa1/2* deletion also caused a subtle but highly reproducible change in tumor morphology, characterized by smaller, more compact nuclei (Figure 1A). In contrast, we observed no specific changes in morphology after deletion of either *Foxa1* or *Foxa2* alone (Supplemental Figure 1D). We next wanted to discern whether *Foxa1/2* deletion simply slows the rate of growth or if it causes tumor regression. Using weekly μCT scans of tumor bearing mice, we found that tamoxifen treatment caused an initial decrease in tumor burden, followed by disease stasis over the course of five weeks. In contrast, vehicle treated mice had a continuous increase in tumor burden over the same period of time (Supplemental Figure 1F).

To evaluate long term consequences of *Foxa1/2* deletion, we performed a survival study in which tumors developed for 10 weeks prior to tamoxifen or vehicle injections. At this timepoint, a subset of tumors in the KP mouse model have progressed to higher grade, macroscopic lesions (Caswell et al., 2014). Concomitant deletion of both *Foxa1* and *Foxa2* in established tumors caused a substantial increase in survival, extending median lifespan by 13.5 weeks (Figure 1D). In contrast, deletion of a single allele of both *Foxa1* and *Foxa2* (Figure 1E), or *Foxa2* alone had no impact on median overall survival, and deletion of *Foxa1* alone lead to a slight (1.5 week) increase in median survival (Supplemental Figure 1G-H). We observed high levels of recombination in tamoxifen treated KPF_1_, KPF_2_ and KPF_1_F_2_ mice, although incomplete recombinants could also be detected in most mice by IHC, particularly KPF_1_F_2_ mice that survived for extended periods of time (data not shown). *Foxa1/2* deletion in established tumors also increased survival in a LUAD model driven by BRAF^V600E^ (*BRAF^FSF-V600E/+^; Trp53^Frt/Frt^*)(Dankort et al., 2007), demonstrating that FoxA1/2 are required for the growth of LUAD harboring distinct driver mutations (Supplemental Figure 1I-J). Of note, we detected a higher proportion of incomplete recombinants in tamoxifen treated BPF_1_F_2_ survival mice than KPF_1_F_2_ mice. This is likely due the use of lentivirus in KPF_1_F_2_ mice rather than adenovirus, which may have led to a higher recombination rate of the *Rosa26^FSFCreERT2^* allele. Detailed histological analysis of all survival studies can be found in Supplemental Table 1. Together, these results establish that FoxA1 and FoxA2 are necessary for in vivo growth of NKX2-1-positive LUAD.

We derived cell lines from individual macroscopic KPF_1_F_2_ tumors under either standard 2D (3311) or three-dimensional, Matrigel-based organoid (1027B, 1027D, and 1292B) culture conditions. Treatment with 4-hydroxytamoxifen (4-OHT) generates isogenic pairs of organoids and cell lines that differ solely in the presence or absence of FoxA1/2 (Figure 2A, Supplemental Figure 2A). In agreement with our in vivo data, in vitro deletion of *Foxa1/2* in both cell lines and organoids lead to a significant decrease in proliferation (Figure 2B-C). We detected a transient increase in apoptosis 72 hours after the start of 4-OHT treatment, but did not observe any durable, long term apoptosis induction (Supplemental Figure 2D). In subcutaneous tumors, deletion of *Foxa1/2* initially caused modest but significant tumor regression (p<0.003), followed by slower growth, a pattern which closely matches in vivo MicroCT analysis (Figure 2D, see also 7G). Upon histopathologic analysis, we found that the tamoxifen treated subcutaneous tumors were predominantly FoxA1/2-negative (∼65-100%) and identified two distinct morphologies in *Foxa1/2*-deleted tumors: a moderate to poorly differentiated adenocarcinoma component (similar to controls) and a distinct sarcomatoid/quasi-mesenchymal component, which exhibited moderately lower NKX2-1 levels (Supplemental Figure 2B-C).

**Figure 2:**
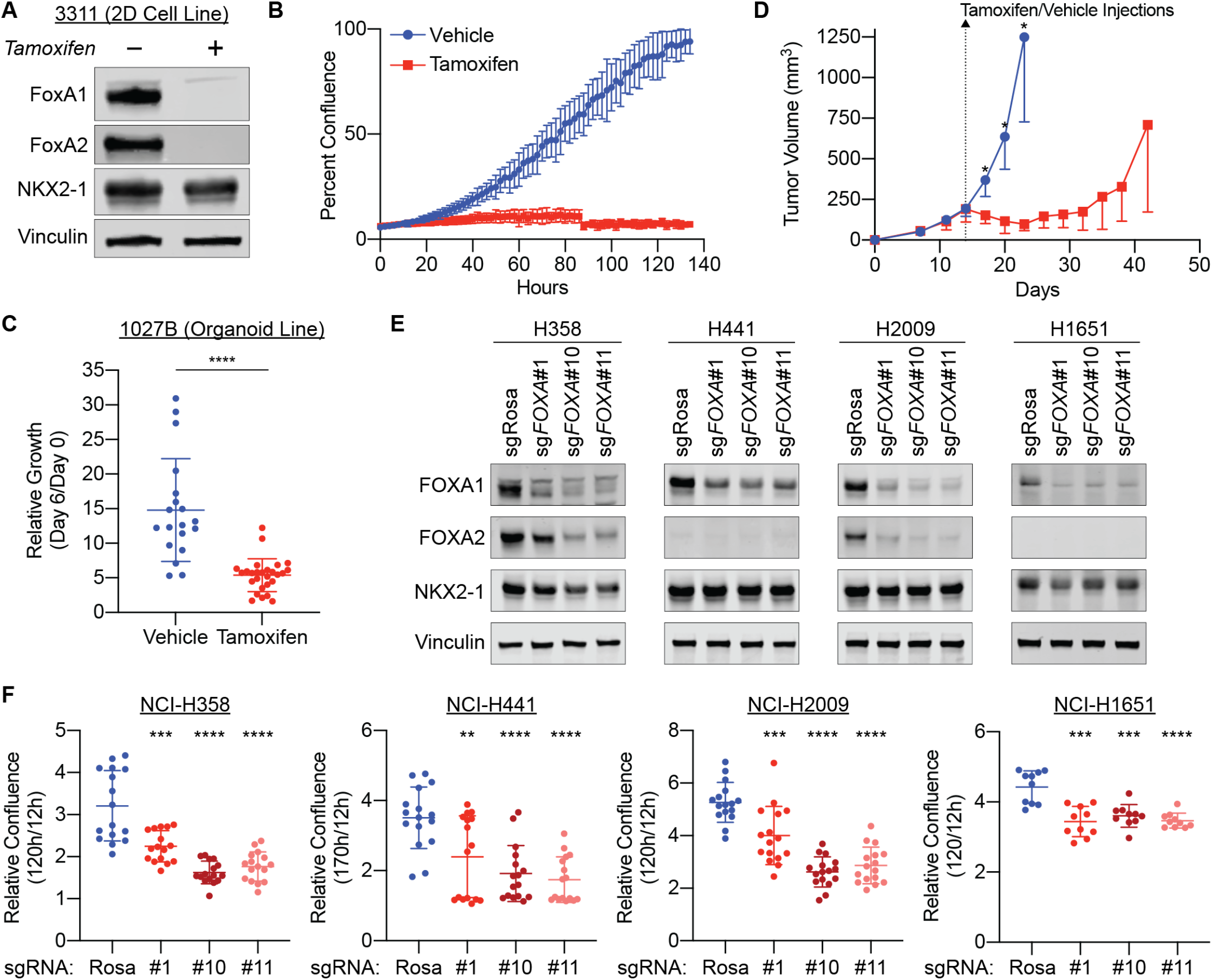
FoxA1/2 are required for growth of NKX2-1-positive lung adenocarcinoma *in vitro*. A) Western blot of protein extract from 2D KPF_1_F_2_ cell line (3311) collected 3 days post 48-hour 4-hydroxytamoxifen treatment demonstrating complete loss of FoxA1/2. B) Incucyte-based confluency growth assay of 3311 treated with vehicle (blue) or 4-OHT (red) one week before seeding 7,000 cells/well (one biological replicate shown; representative of n=3 biological replicates with 8 technical replicates each; p<0.0001 unpaired t-test of endpoint values). C) Presto Blue proliferation assay of 1027B KPF_1_F_2_ organoid line 6 days post 48-hour treatment with vehicle or 4-OHT (Day 6 reads normalized to Day 0 reads; p<0.0001 unpaired t-test). D) Subcutaneous tumors (3311 cell line) were allowed to grow to an average of 200mm^3^ before treating with either vehicle (n=5) or tamoxifen (n=5) and progressed until one tumor from the treatment cohort surpassed 1000mm^3^ (unpaired t-test day 17 p=0.0027; day 20 p=0.0006; day 23 p=0.0012; representative of one of two independent experiments shown; Graph indicates mean + S.D.) E) Western blot of protein extract from NCI-H358, NCI-H441, NCI-H2009, and NCI-H1651 cell lines transduced with three separate lentivectors that simultaneously express unique sgRNAs targeting both FOXA1 and FOXA2 (sg*FOXA*#1, sg*FOXA*#10, and sg*FOXA*#11), as well as a dual sgRNA control targeting the Rosa locus (sgRosa). Protein collected concurrently with proliferation assay seeding (6 days post-lentiviral sgRNA transduction/2 days post-selection). F) Relative increase in percent confluence in Incucyte-based growth assay: percent confluence at 120 or 170 hours normalized to percent confluence 12 hours post-seeding (n=2 biological replicates with 8 or 5 technical replicates each; unpaired t-test relative to sgNT **p<0.005, ***p<0.001, ****p<0.0001).

We next examined FOXA1/2 dependence in NKX2-1-positive human LUAD cell lines by knocking out *FOXA1/2* via CRISPR/Cas9. To accomplish this, we selected three cell lines harboring *KRAS* mutations that express NKX2-1, as well as one or both of the FoxA1/2 proteins: NCI-H358, NCI-H441, and NCI-H2009. We also targeted *FOXA1/2* in NCI-H1651, an NKX2-1-positive lung adenocarcinoma cell line that co-expresses pulmonary and gastric markers. Cells stably expressing Cas9 were transduced with three independent lentivectors that express unique sgRNAs targeting both *FOXA1* and *FOXA2* (sg*FOXA*#1, sg*FOXA*#10, and sg*FOXA*#11), as well as a non-targeting dual sgRNA control (sgNT). In all four cell lines, *FOXA1/2* knockout inhibited proliferation (Figure 2E-F). The three sgRNA combinations have varied levels of efficiency in targeting *FOXA1/2* in the *KRAS* mutant lines: sg*FOXA*#1 yields the smallest decrease in FOXA1/2 protein levels, while both sg*FOXA*#10 and sg*FOXA*#11 reduce FOXA1/2 protein levels to a much greater extent (Figure 2E). The difference in *FOXA1/2* targeting efficiency correlates directly with decrease in proliferation: in *KRAS* mutant lines the least efficient sg*FOXA*#1 yields an average of 40% decrease in proliferation, while the more efficient sg*FOXA*#10 and sg*FOXA*#11 cause a 94% and 90% decrease, respectively (Figure 2F). Combined, these murine and human results demonstrate that FoxA1/2 are required for the growth of NKX2-1 positive LUAD in vivo and in vitro.

### FoxA1/2 enforce GI-related and Alveolar Type 2 programs in NKX2-1-positive lung adenocarcinoma

We next sought to determine whether FoxA1/2 regulate cellular identity programs associated with malignant progression in NKX2-1-positive LUAD. Recent single cell analyses of the KP LUAD GEMM, including in vivo lineage recording of cancer progression, defined multiple transcriptional programs associated with distinct cellular identities that emerge as tumors progress (LaFave *et al*., 2020; Marjanovic *et al*., 2020; Yang et al., 2021). Early low grade tumor cells most closely resemble normal AT2 cells, as evidenced by high levels of *Nkx2-1* and its canonical AT2 targets. As tumors progress (12-30 weeks after initiation), subsets of higher grade LUAD cells upregulate distinct gastrointestinal (GI) transcriptional programs despite the continued expression of *Nkx2-1*. (For simplicity, we use the term “dual identity” to refer to cells in which the AT2 program is simultaneously expressed with GI programs.) In late stage disease, a sub-population of cells reach a terminal poorly differentiated/EMT-like transcriptional state that lacks both AT2 and GI differentiation. Transition through dual identity states is a key component of LUAD malignant progression, but the mechanisms driving these lineage changes are not fully defined.

In order to determine which cellular identities are regulated by FoxA1/2 in NKX2-1-positive LUAD, we performed scRNAseq on in vivo KPF_1_F_2_ and KP tumors initiated with Ad5mSPC-FlpO (to specifically target SPC-positive epithelial cells). Mice were injected with tamoxifen 10 weeks post tumor-initiation. Twelve weeks after tumor initiation, we macrodissected individual tumors, digested into single cell suspensions, depleted CD45+ and CD31+ cells via magnetic activated cell sorting (MACS) to increase representation of tumor cells, and sequenced cells both pre-depletion and post-depletion (n=2 mice per genotype, multiple tumors per mouse). We visualized single cells in reduced dimensionality using uniform manifold approximation and projection (UMAP) and identified 25 major clusters between both genotypes (Supplemental Figure 3A). Of the 25 clusters, 10 were identified as tumor cells based on expression of FlpO-induced *Cre^ERT2^*transcripts and low levels of stromal specific marker genes (Supplemental Figure 3B; Supplemental Table 2). The remaining 15 clusters were identified as immune cells, endothelial cells, fibroblasts, and normal lung epithelium (Supplemental Figure 3C, 5A;Supplemental Table 2). Analysis of only the tumor cell population identified 13 major clusters (Figure 3A), of which six are largely KP specific (Clusters 1, 3, 4, 5, 6, and 12), four are KPF_1_F_2_ specific (Clusters 2, 7, 8, and 9), and three contain cells from both KP and KPF_1_F_2_ mice (Clusters 0, 10, and 11; Figure 3B, Supplemental Figure 3D-E). Upon closer inspection, we found that the cells from KPF_1_F_2_ samples that cluster with the KP tumor cells (Clusters 0, 10, and 11) are *Foxa1/2*-positive incomplete recombinants, based on equivalent levels of the *Foxa1* exon flanked by loxP sites in both KP and KPF_1_F_2_ cells (Figure 3C). In contrast, cells in KPF_1_F_2_ specific clusters appear to be complete recombinants based on much lower levels of this exon compared to KP specific clusters (Supplemental Figure 3F). The observation that multiple clusters contain either normal stromal cells or unrecombined tumor cells from mice of both genotypes suggests that cell type, rather than genotype per se, is the primary driver of clustering. Thus, the near-complete separation of FoxA1/2-negative and FoxA1/2-positive tumor cells shows that in vivo *Foxa1/2* deletion has a major impact on the transcriptome of LUAD cells. To complement this in vivo analysis, we also performed bulk RNAseq on isogenic KPF_1_F_2_ organoid pairs. Utilizing DESeq2 we identified 1,914 differential genes (padj<0.05). Additional filtering for the scale of fold change (minimum 1.5-fold, or log2FC>±0.585) identified 1,692 differentially expressed genes: 918 in FoxA1/2-positive organoids and 774 in FoxA1/2-negative organoids (Supplemental Table 3).

**Figure 3:**
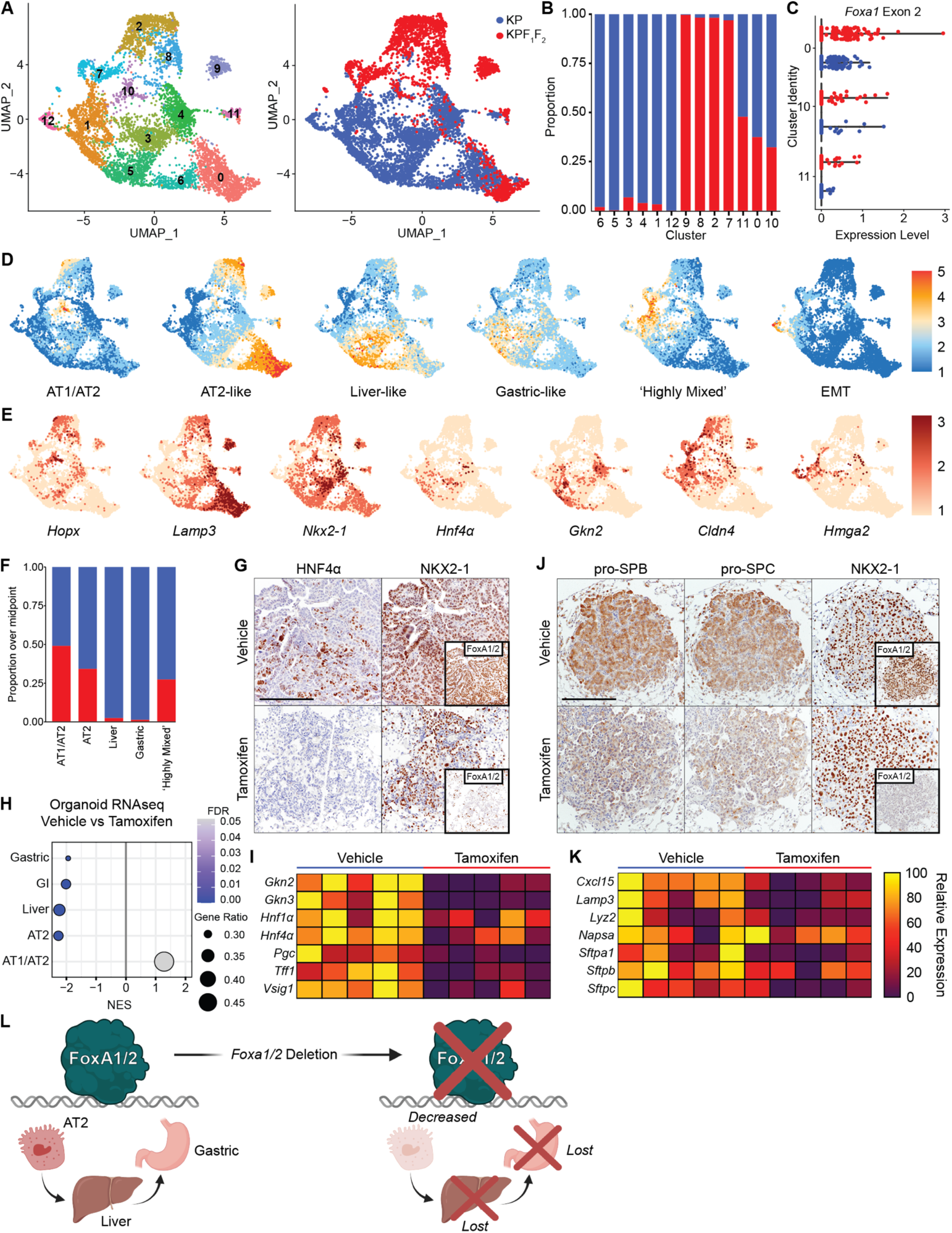
FoxA1/2 enforce AT2 and GI-related transcriptional programs in NKX2-1-positive LUAD. A) UMAP of cells from KP (4646 cells) and KPF_1_F_2_ (2481 cells) autochthonous tumors. Left panel colored by cluster identity and right panel colored by genotype. B) Proportion of cells in each cluster from KP or KPF_1_F_2_ tumors. C) Expression level of the second exon of *Foxa1* that should be excised by Cre-ERT2. Detection of this exon within KPF_1_F_2_ cells indicates retained expression of *Foxa1/2* due to incomplete Cre-mediated recombination. D) UMAPs of gene module scores of dual-identity transcriptional programs associated with tumor progression as defined by Marjanovic et al. E) UMAPs of relative expression of genes corresponding to above programs (*Hopx*/AT1-AT2; *Lamp3* and *Nkx2-1*/AT2; *Hnf4α* and *Gkn2*/Liver and Gastric; *Cldn4*/Highly Mixed; *Hmga2*/EMT). F) Proportion of FoxA1/2-positive (blue) and FoxA1/2-negative (red) cells with gene module scores at a level at or over the midpoint of the range of scores for the indicated transcriptional programs. G) Representative IHC images of gastric transcription factor HNF4α and pulmonary transcription factor NKX2-1 in FoxA1/2-positive and FoxA1/2-negative tumors, illustrating the co-expression of both endodermal and pulmonary transcription factors in KP tumors and the loss of the gastric lineage upon *Foxa1/2* deletion (scale bar 500 μm). H) Normalized Enrichment Score (NES) as determined by GSEA of organoid RNAseq for AT2, AT1/AT2, and GI-related transcriptional programs as defined by Marjanovic et al, illustrating that the AT2, Liver, Gastric, and GI transcriptional programs are all enriched in FoxA1/2-positive organoids. Positive NES indicates enrichment in FoxA1/2-negative organoids and negative NES indicates enrichment in FoxA1/2-positive organoids. Gene ratio representing the proportion of genes within a gene signature detected in the DEG list. I) Relative expression of select gastric marker genes in FoxA1/2-positive (vehicle) and FoxA1/2-negative (tamoxifen) organoid cultures. J) Representative IHC images of AT2 markers pro-SPB, pro-SPC, and pulmonary transcription factor NKX2-1 in FoxA1/2-positive and FoxA1/2-negative tumors, illustrating the decrease in AT2-like lineage upon *Foxa1/2* deletion (scale bar 500 μm). K) Relative expression of select AT2-specific genes in FoxA1/2-positive (vehicle) and FoxA1/2-negative (tamoxifen) organoid cultures. L) In summary, FoxA1/2 are required for the full expression of AT2, gastro-intestinal, and EMT identities associated with LUAD progression. Deletion of *Foxa1/2* causes a distinct loss of liver, gastric, and EMT differentiation states and partially decreases the AT2 transcriptional program (image created in BioRender).

We first evaluated the expression of critical transcriptional programs that arise throughout LUAD development as described by (Marjanovic *et al*., 2020). Across the KP clusters we identified cells expressing the entire continuum of identities that arise throughout tumor evolution (Figure 3D-E; Supplemental Figure 3F). Cluster 0 is strongly associated with the AT2-like program expressed by cells at an early timepoint in tumor evolution (*Lamp3* and *Nkx2-1*). Cluster 10 expresses high levels of a mixed AT1/AT2-like pulmonary identity (*Hopx*). Clusters 3, 4, 5, and 6 retain expression of *Nkx2-1* and the AT2-like program, but also gain expression of the Embryonic Liver-like program and Gastric-like programs (*Hnf4a* and *Gkn2*); these clusters contain cells with a critical dual-identity state that emerges as KP LUAD tumors evolve. Cells in Cluster 1 express both Gastric-like and GI epithelium-like programs, as well as the “Highly Mixed” transcriptional program associated with the high-plasticity cell state described by (Marjanovic *et al*., 2020) (*Cldn4*). Finally, in Cluster 12 we isolated a small set of cells from KP mice that express the terminal, highly aggressive EMT transcriptional program that is found only in advanced, high grade adenocarcinomas (*Hmga2*). As expected, cells in the G2/M phase cluster separately from other tumor cell populations (Cluster 11; Supplemental Figure 3F).

Because KP control tumors contain the entire spectrum of cellular identities that emerge throughout LUAD progression in this model, we can examine differences between the KP and KPF_1_F_2_ clusters to determine which LUAD differentiation states are dependent on *Foxa1/2* expression. We found that FoxA1/2-negative cells still express markers of the combined AT1/AT2 identity (as shown by expression of *Hopx* in Clusters 2, 8, and 9) and the Highly Mixed transcriptional state (as shown by expression of *Cldn4* in Clusters 2, 7 and 9) (Figure 3D-F). However, the data reveal that FoxA1/2 are required for full activation of the AT2-like and Gastric/Liver-like states.

Our most striking observation is that the dual-identity, simultaneously pulmonary and gastric cellular state is completely dependent on FoxA1/2. Embryonic Liver-like and Gastric-like transcriptional programs are not expressed in *Foxa1/2*-deleted clusters (*Hnf4a* and *Gkn2*; Figure 3D-E, Supplemental Figure 3F,H). Nearly 100% of the cells with high expression of the Liver and Gastric programs come from KP-specific clusters (Figure 3F). This loss of a gastric identity is consistent at the protein level in tumors and organoids; a subset of FoxA1-2-positive cells are positive for both NKX2-1 and HNF4α, but FoxA1/2-negative cells are completely negative for HNF4α (Figure 3G, Supplemental Figure 3G). In addition, HNF4α levels decrease following FOXA1/2 inhibition in the dual identity human LUAD cell line NCI-H1651 (Supplemental Figure 3I). In organoid bulk RNAseq data, we see an overall enrichment for the Gastric, GI epithelium, and Liver transcriptional programs in FoxA1/2-positive organoids (Figure 3H), and at an individual gene level, canonical gastric and endodermal markers are significantly decreased following *Foxa1/2* deletion (Figure 3I). These data show that FoxA1/2 are critical drivers of the gastric/endodermal transcriptional programs that are a critical phase of LUAD evolution.

FoxA1/2 are also important drivers of the AT2-like cellular identity, albeit to a lesser extent than the Gastric and Liver identities. The AT2-like program is very low in KPF_1_F_2_ Cluster 7, and is expressed in Clusters 2, 8 and 9 at a lower level than the KP AT2-like Cluster 0 (*Lamp3* and *Nkx2-1*; Figure 3D-E; Supplemental Figure 3F). At the individual gene level, we find that the decline in AT2 identity following *Foxa1/2* deletion is a consequence of partial rather than complete downregulation of many AT2 marker genes in Clusters 2, 8 and 9 when compared to the AT2-like Cluster 0 (Supplemental Figure 3H,J). Moreover, there are specific AT2 marker genes in each cluster (such as *Napsa* in Clusters 8 and 2) that are expressed at approximately the same level as cluster 0 (Supplemental Figure 3H). Consistent with these observations, we observe a decrease, but not complete loss, of AT2 markers pro-SPB and pro-SPC by IHC after *Foxa1/2* deletion (Figure 3J). In vitro, we also see an overall enrichment of the AT2 gene signature in FoxA1/2-positive organoids (Figure 3H). However, at an individual gene level we see that some AT2 markers are significantly decreased following *Foxa1/2* deletion (*Cxcl15, Lamp3, Lyz2*, *Sftpa1* and *Sftpc*) while others are only slightly decreased (*Sftpb*) or virtually unchanged (*Napsa*; Figure 3K). Additionally, expression of AT2 marker genes *SFTPA1* and *SFTPB* significantly decreases following FOXA1/2 knock down in human LUAD cell line NCI-H441 (Supplemental Figure 3K). These data show that FoxA1/2 are needed for maximal expression of the AT2-like transcriptional program, but other pulmonary transcription factors are able to act independently of FoxA1/2 to activate some elements of AT2 identity.

Altogether, we find that FoxA1/2 are critical drivers of the dual-identity state that emerges as LUAD tumors evolve. Upon *Foxa1/2* deletion, tumors decrease their expression of an AT2-like identity and completely lose gastric and endodermal transcriptional programs (Figure 3L).

### FoxA1/2 suppress alternative pulmonary and stratified squamous transcriptional programs in NKX2-1-positive lung adenocarcinoma

We next sought to understand the transcriptional state(s) to which LUAD cells equilibrate in the absence of FoxA1/2. We first evaluated relative levels of transcriptional programs associated with normal lung epithelium in more depth using gene signatures of normal AT1 and AT2 cells derived from the same study (Travaglini et al., 2020). We found that the Travaglini AT2 signature is detectable in FoxA1/2-negative Clusters 2, 8, and 9 (Figure 4A), but at a lower level than Cluster 0 FoxA1/2-positive cells, consistent with our analysis of the Marjanovic AT2-like signature (Figure 3D-E, Supplemental Figure 3J). The Travaglini AT1 signature is readily detectable in Clusters 2 and 7 (Figure 4B), as are an AT1 signature from the LungGENS database(Du et al., 2015) (Supplemental Figure 4A) and AT1 genes uniquely dependent on NKX2-1 as defined by Little et al(Little *et al*., 2019) (Figure 4C). Close inspection of Cluster 2 reveals an inverse correlation between AT2 and AT1 identities within this cluster (Supplemental Figure 4B-C). FoxA1/2-mediated inhibition of AT1 differentiation is also evident in LUAD organoids. Using Gene Set Enrichment Analysis (GSEA), we found that *Foxa1/2* deletion in organoids impairs AT2 differentiation while promoting AT1 differentiation (Figure 3H, 4D). At the protein level, a higher proportion of LUAD tumors are positive for the AT1 marker HOPX following *Foxa1/2* deletion (Figure 4E). Together these data suggest that loss of FoxA1/2 promotes an AT1 cellular state.

**Figure 4:**
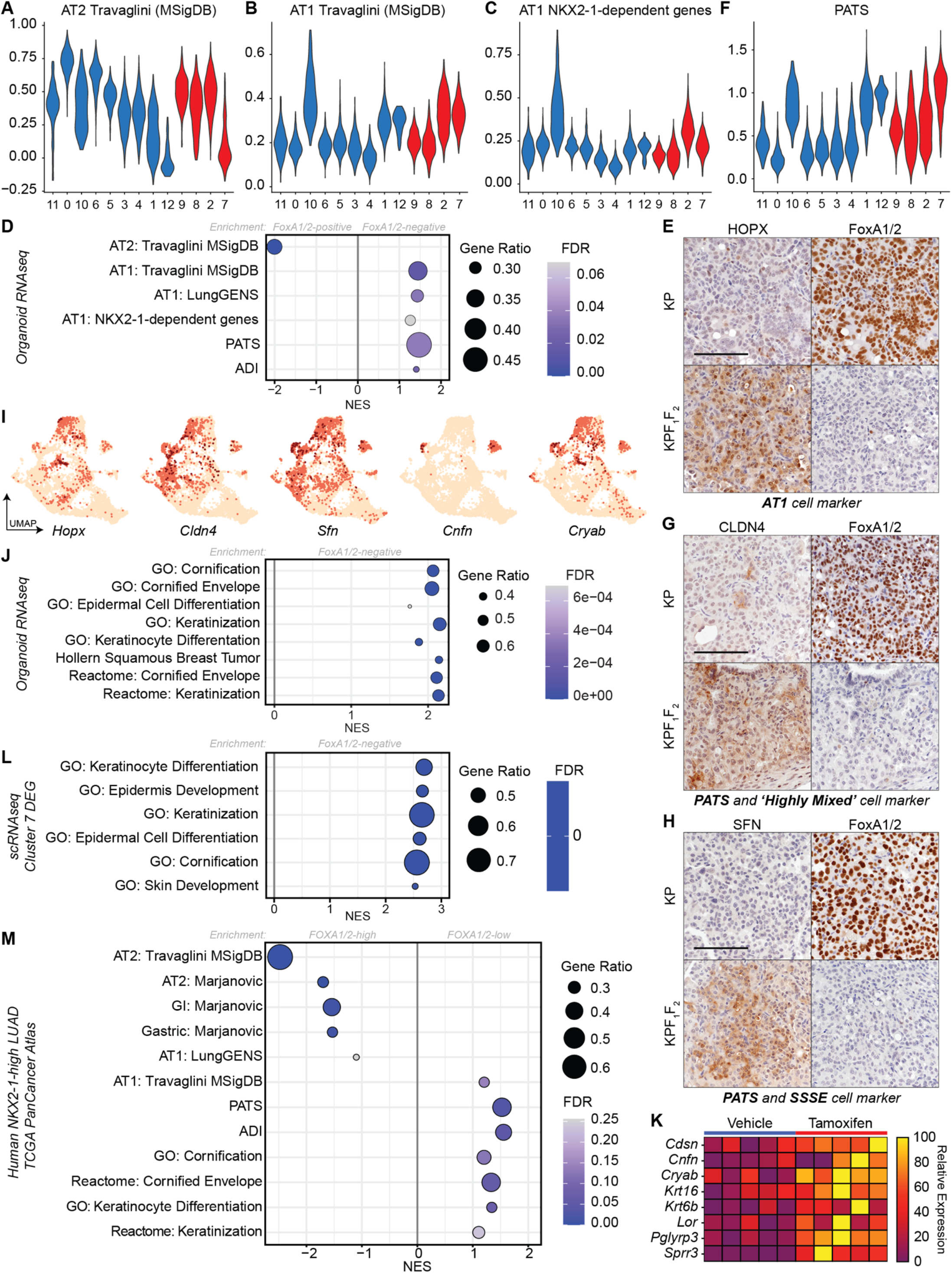
FoxA1/2 suppress alternative pulmonary and stratified squamous transcriptional programs in NKX2-1-positive lung adenocarcinoma. A) Violin plot representing the expression of the Travaglini AT2 transcriptional program in each tumor cell cluster. The y-axis indicates gene module score. The x-axis indicates cluster identity. B) Violin plot representing the expression of the Travaglini AT1 transcriptional program in each tumor cell cluster. C) Violin plot representing the expression of NKX2-1-dependent AT1 genes in each tumor cell cluster. D) Normalized Enrichment Score (NES) as determined by GSEA of KPF_1_F_2_ organoid RNAseq for listed gene sets. Positive NES indicates enrichment in FoxA1/2-negative organoids and negative NES indicates enrichment in FoxA1/2-positive organoids. E) Representative IHC images for AT1 marker HOPX in KP and KPF_1_F_2_ tumors, illustrating an increase in HOPX protein levels following *Foxa1/2* deletion (Scale bar 250 μm). F) Violin plot representing the expression of the PATS transcriptional program in each tumor cell cluster. G) Representative IHC images for PATS and ‘Highly Mixed’ marker CLDN4, illustrating an increase in CLDN4 protein levels following Foxa1/2 deletion (Scale bar 250 μm). H) Representative IHC images for PATS and SSSE marker SFN, illustrating an increase in SFN protein levels following Foxa1/2 deletion (Scale bar 250 μm). I) UMAP representing relative expression of AT1, PATS, and SSSE marker genes in tumor cells (AT1: *Hopx* and *Cryab*; PATS: *Cldn4* and *Sfn*; SSSE: *Sfn, Cnfn,* and *Cryab*) J) NES of representative gene sets associated with a squamous differentiation from GSEA intersection of the differentially expressed genes identified in KPF_1_F_2_ organoid RNAseq with all available MSigDB gene set collections. Positive NES indicates enrichment in FoxA1/2-negative organoids. K) Relative expression of select SSSE genes in FoxA1/2-positive (vehicle) and FoxA1/2-negative (tamoxifen) organoid cultures. L) NES of gene ontology signatures associated with keratinocyte differentiation from GSEA analysis of FoxA1/2-negative Cluster 7. Positive NES indicates enrichment in Cluster 7 cells relative to all other cells. M) GSEA analysis of differentially expressed genes identified between *FOXA1/2*-high and *FOXA1/2*-low cohorts in in *NKX2-1*-high human LUAD patients (TCGA Pan Cancer Atlas). Gene sets representing normal AT1, AT2, Gastric, PATS, ADI, and SSSE cell types were analyzed. We found significant enrichment for AT2 and Gastric programs in the *FOXA1/2*-high patient cohort (negative NES score) and significant enrichment for AT1, PATS, ADI, and SSSE cell signatures in the *FOXA1/2*-low patient cohort (positive NES score).

Like the AT1/AT2-like LUAD program, the ‘Highly Mixed’/*Cldn4*-high program from (Marjanovic *et al*., 2020) is found in both FoxA1/2-positive and FoxA1/2-negative clusters (Cluster 1 vs. 7, Figure 3 D-F). Interestingly, three recent publications have described a distinct *Cldn4*-high alveolar cell state that emerges as AT2 cells differentiate into AT1 cells in response to lung injury, which (Kobayashi et al., 2020) have termed Pre-Alveolar type-1 Transitional cell State (PATS; also described by (Strunz et al., 2020) as Alveolar Differentiation Intermediate - ADI, and (Choi et al., 2020) as Damage-Associated Transient Progenitors - DATPs). They found that this intermediate alveolar cell type is not simply undergoing a gradual loss of AT2 characteristics while acquiring AT1 characteristics; rather this is a unique transitional cell population that emerges in response to injury and is marked by high *Cldn4* levels and activation of signaling pathways such as NFκB and TGFβ. The PATS and ADI signatures are both strongly expressed in a subset of KP and KPF_1_F_2_ LUAD cells and overlap with the ‘Highly Mixed’ LUAD signature in our dataset (Figure 4F; Supplemental Figure 4D). We see expression of PATS/ADI markers CLDN4 and SFN following *Foxa1/2* deletion in tumors at the protein level (Figure 4G-H; gene expression UMAPs 4I). Additionally, GSEA reveals enrichment of these gene sets in organoids upon loss of FoxA1/2 in vitro (Figure 4D). This suggests that the LUAD cells previously described as ‘Highly Mixed’ may have activated the transcriptional program of PATS/ADI transitional cells that emerge in response to lung injury. We note that in scRNA-seq data, levels of both PATS/ADI and AT1 signatures are not necessarily higher in specific *Foxa1/2*-deleted clusters compared to controls, but that a higher fraction of cells express these programs, which likely explains their relative enrichment in bulk RNA-seq data.

Global pathway analysis (GSEA) of organoid RNAseq provided additional insights into the cellular identities adopted by FoxA1/2-negative cells. We find consistent enrichment of gene sets canonically expressed in maturing stratified squamous epithelium in FoxA1/2-negative organoids across multiple MSigDB collections. Genes sets associated with keratinocyte differentiation, the cornified envelope, and more broad categorizations of squamous tumors and epidermal cell differentiation are all enriched in FoxA1/2-negative organoids (Figure 4J-K, Supplemental Table 5). GSEA of differentially expressed genes in each cluster (Supplemental Table 3) revealed that a similar array of gene sets associated with stratified squamous differentiation were highly enriched in FoxA1/2-negative Cluster 7 (Figure 4L). Specific genes highly expressed in FoxA1/2-negative cells in vivo include *Sfn*, a marker of keratinocytes and PATS/ADI cells, *Cnfn*, a marker of the superficial layers of the stratified squamous epithelium (hereafter SSSE), and *Cryab*, a reported marker of SSSE and AT1 cells (Figure 4I). At the protein level, the majority of FoxA1/2-negative tumors are positive for SFN (>75%), while most FoxA1/2-positive tumor lacks SFN staining (Figure 4H; positive staining in SSSE Supplemental Figure 4E). IHC for alpha crystallin-β chain (the *Cryab* gene product), reveals expression specifically in a subpopulation of *Foxa1/2* deleted tumor cells, but not controls. Of note, in our analysis this protein was detectable in normal esophageal SSSE but not AT1 cells (Supplemental Figure 4F). Despite induction of several SSSE marker genes, ΔNp63 is undetectable in FoxA1/2-negative tumor cells (Supplemental Figure 4G). ΔNp63 is a master regulator of squamous differentiation that is normally expressed in the basal layer of stratified squamous epithelia, including the stem cell compartment, and can function as an oncogene in squamous cell carcinoma (SCC) (Moses et al., 2019). Thus, *Foxa1/2* deletion causes an incomplete shift toward a squamous differentiation program in a subset of LUAD cells, but the absence of ΔNp63 may prevent full squamous transdifferentiation.

We next wanted to assess whether these transcriptional patterns exist in human disease, so we analyzed *NKX2-1*-high human LUAD samples from TCGA’s PanCancer Atlas (n=402/516) and grouped patients into *FOXA1/2*-high and *FOXA1/2*-low cohorts (n=114 and 23, respectively). We performed differential gene expression analysis between these cohorts using DESeq2 and then assessed the enrichment of relevant transcriptional programs via GSEA (Supplemental Table 3). In support of our in vivo and in vitro murine RNAseq data, we find that AT2 and Gastric identities are significantly enriched in tumors with high *FOXA1/2* expression. In contrast, tumors with low *FOXA1/2* levels exhibit a significant enrichment of AT1, PATS/ADI, and Keratinocyte differentiation states (Figure 4M).

Taken together, these data show that FoxA1/2 inhibit alternative alveolar and stratified squamous differentiation states in NKX2-1-positive LUAD. Specifically, *Foxa1/2* deletion causes a partial loss of AT2-like identity in LUAD, with a concomitant increase in the percentage of cells expressing markers of alternative alveolar cell types, such as AT1 or PATS. Moreover, a subset of LUAD cells also de-repress transcriptional programs associated with the SSSE after *Foxa1/2* deletion.

### Simultaneous deletion of both *Foxa1* and *Foxa2* is required for cellular identity transformations

In order to understand the individual contributions of FoxA1 and FoxA2 to the cellular differentiation phenotypes described in Figures 3 and 4, we performed RNAseq on isogenic pairs of KPF_1_ and KPF_2_ organoids (n=3 and n=2 independent organoid lines, respectively). *Foxa1* deletion led to 363 differentially expressed genes (DEGs, padj<0.05). With additional filtering for the scale of fold change (minimum 1.5-fold, or log2FC>±0.585) we found 284 DEGs: 244 in FoxA1-positive organoids and 40 in FoxA1-negative organoids. *Foxa2* deletion yields 893 DEGs. After filtering for a 1.5-fold change, we identified 760 DEGs: 435 in FoxA2-positive organoids and 325 in FoxA2-negative organoids. In contrast, simultaneous *Foxa1* and *Foxa2* deletion led to 1692 DEGs (padj<0.05 and log2FC>±0.585).

We then intersected DEGs from each genotype and found that 75% of gene expression changes (1269 of 1692 genes) observed in KPF_1_F_2_ isogenic organoid pairs are unique to this genotype. Thus, the large majority of observed transcriptional changes require loss of both FoxA1 and FoxA2. Of the remaining 25% of genes, 19% are shared between simultaneous *Foxa1/2* deletion and deletion of *Foxa2* alone (322 of 1692 genes), 2.7% are shared between deletion of *Foxa1/2* and deletion of *Foxa1* alone (45 of 1692 genes), and 3.3% are shared between all three genotypes (56 of 1692 genes; Supplemental Figure 4H). In addition to yielding fewer DEGs overall, single deletion of either *Foxa1* or *Foxa2* results in gene expression changes that are significantly lower in magnitude than deletion of both. DEGs following *Foxa1* deletion exhibit a range of -1.8 to 1.7 Log2 fold change while *Foxa2* deletion results in DEGs with a range of -4.4 to 4.1 Log2 fold change. Simultaneous deletion of *Foxa1/2* yields increases in gene expression as large as 9.1 Log2 fold change and decreases up to 5.4 Log2 fold change (Supplemental Figure 4I). These data reveal that FoxA1 plays a notably smaller role in transcription than FoxA2, but loss of both paralogs is required for maximal transcriptional changes.

Although there are substantially fewer differentially expressed genes following individual deletion of *Foxa1* or *Foxa2*, we proceeded to investigate the individual contribution of each paralog in regulating cellular identity. Deletion of either *Foxa1* or *Foxa2* alone, does not inhibit gastric or AT2 identities at the gene expression or protein level. Expression levels of most GI and AT2 genes do not significantly change following deletion of either *Foxa1* or *Foxa2* alone (Supplemental Figure 4J). The GI and AT2 genes that decrease following individual deletion do not exhibit the same magnitude of change as dual deletion (KPF_1_: *Lamp3* and *Sftpa1*; KPF_2_: *Lamp3*, *Tff1*, *Cxcl15*, and *Sftpc*). We also do not observe changes in HNF4α or pro-SPB protein levels following individual *Foxa* deletion in vivo (Supplemental Figure 4K-L). Additionally, individual deletion does not result in induction of the SSSE differentiation as seen with combined FoxA1/2 loss (Supplemental Figure 4J). Together, these data illustrate the functional redundance of FoxA1 and FoxA2; concomitant loss of both transcription factors is required to observe changes in tumor growth and progression (Figure 1B-E; Supplemental Figure 1 G-H) and changes in differentiation state.

### FoxA1/2 loss leads to an M1-polarization in resident alveolar macrophages and increases neutrophil recruitment to the tumor microenvironment

To determine whether deletion of *Foxa1/2* in tumor cells has an effect on the tumor microenvironment (TME), we profiled each stromal cell population in our scRNAseq dataset (Supplemental Figure 3A-C, Supplemental Figure 5A). While we did not identify major differences in the transcriptional profiles of T cells, B cells, fibroblasts, or endothelial cells between the two genotypes (not shown), we did identify significant transcriptional differences within two cell compartments: cells of the mononuclear phagocyte system (MPS) and neutrophils. Analysis of the MPS cells (monocytes, macrophages and dendritic cells (DCs)) revealed multiple transcriptionally defined clusters corresponding to four established cell types based on expression of canonical marker genes (*Siglecf, Pparg,* and *Marco* for tissue resident alveolar macrophages*, Mafb* and *C1q* for interstitial macrophages, *Flt3* and *Cd11c* for DCs, *Ly6c2* and *Ace* for other monocytes; Supplemental Figure 5B-E, Supplemental Table 2) (Casanova-Acebes et al., 2021; Maier et al., 2020). While most MPS clusters overlapped between genotypes, we identified two distinct clusters of alveolar macrophages that were largely unique to either KPF_1_F_2_ tumors (Cluster 1) or KP tumors (Cluster 0). Interestingly, the top differentially expressed gene between these two clusters was *Il1b*, a pro-inflammatory cytokine that marks M1-polarized macrophage populations (Supplemental Figure 5E,G, Supplemental Table 4). Macrophages expressing an M1-like transcriptional signature suppress tumorigenesis in various cancers, including in LUAD (Yuan et al., 2015). We found *Il1b* and several additional M1-associated genes enriched in Cluster 1 compared to Cluster 0 (Supplemental Figure 5G). In addition, GSEA enrichment analysis on the differentially expressed gene list between Cluster 1 and Cluster 0 revealed pro-inflammatory TNFα signaling as the top statistically enriched pathway (Supplemental Figure 5F). Together, these results indicate that alveolar macrophages in KPF_1_F_2_ tumors adopt a pro-inflammatory profile, which could contribute to slower tumor growth in these mice. Intriguingly, *Il1b* secreted from macrophages also promotes the PATS/DATP intermediate cell state in alveolar regeneration models (Choi *et al*., 2020). Thus, macrophages in the TME of KPF_1_F_2_ tumors may be poised to activate and/or reinforce the PATS/DATP transcriptional state that is enriched in tumor cells upon *Foxa1/2* deletion (Figure 4).

In our immune analysis, we also noted an enrichment in the neutrophil recruitment chemokines *Cxcl1* and *Cxcl2* in Cluster 1 relative to Cluster 0 (Supplemental Figure 5G). This was intriguing given the recent observation that *Nkx2-1* deletion LUAD accelerates transdifferentiation from LUAD to lung squamous cell carcinoma by enhancing neutrophil recruitment to the TME (Mollaoglu et al., 2018). Because AT2 identity is also partially lost upon *Foxa1/2* deletion in the present study, we hypothesize that in the KPF_1_F_2_ model both macrophages and tumor cells may coordinately recruit neutrophils to the TME where they might similarly impact tumor cell state transitions. Indeed, we found that in addition to the increased *Cxcl1* and *Cxcl2* expression in KPF_1_F_2_ alveolar macrophages above, KPF_1_F_2_ tumor cells express higher levels of *Cxcl1, Cxcl2*, and *Cxcl5* than KP tumor cells (Supplemental Figure 5H).

Analysis of the neutrophil population (i.e. the *Ly6g*, *S100a8*, *S100a9*, and *Mmp9* positive cluster of cells identified in Supplemental Figure 5A) revealed cellular subtypes with differences in the expression of *Siglecf*, an established marker of pro-tumorigenic neutrophils in the KP model (Engblom et al., 2017; Zilionis et al., 2019) (Supplemental Figure I-K). In our analysis, Siglecf^high^ neutrophils expressed additional genes associated with the pro-tumorigenic subtype including *Clec4n*, *Xbp1*, and *Car4* while the Siglecf^low^ population expressed higher levels of canonical neutrophil markers including *S100a9, S100a8, Mmp8, and Ly6g* (Engblom *et al*., 2017; Zilionis *et al*., 2019) (Supplemental Figure 5L-M). While the Siglecf^high^ population was more evenly distributed between genotypes, the Siglecf^low^ cluster was almost entirely derived from KPF_1_F_2_ samples (Supplemental Figure 5O). Of note, the Cxc chemokine receptor 2 gene (*Cxcr2)*, was found to be expressed at higher levels in the KPF1F2-specific Siglecf^low^ cluster compared to the Siglecf^high^ cells (Supplemental Figure 5N). This may indicate specific recruitment of this potentially functionally distinct Siglecf^low^ neutrophil subset to the TME by *Cxcl1/2/5*-expressing tumor cells and macrophages upon loss of FoxA1/2 in tumor cells (Supplemental Figure 5N). It remains to be determined whether the change in neutrophil recruitment to the TME reinforces the SSSE program upon loss of FoxA1/2 *in vivo*.

Finally, we asked whether there was evidence that the increase in M1 macrophage markers and neutrophil chemoattractant genes might reflect distinct multicellular interactions within the tumor-myeloid axis specifically in KPF_1_F_2_ tumors. To model distinct patterns of intercellular communication between major immune cell subtypes in our scRNAseq dataset, we utilized the CellChat computational package (Jin et al., 2021). CellChat analysis inferred stronger interactions between neutrophils, tumor cells, and MPS cells in KPF_1_F_2_ (red) vs. KP (blue) tumors (Supplemental Figure 5P). We predict that increased communication between these populations could reinforce PATS/ADI and SSSE differentiation programs in KPF_1_F_2_ tumor cells *in vivo*, while FoxA1/2 loss concomitantly induces pro-inflammatory programs that specifically hinder LUAD tumor growth. Nevertheless, we also observe induction of these transcriptional programs in vitro (Figure 4D, J), suggesting that the TME is not strictly required for acquisition of these programs, but rather reinforces them in vivo.

### NKX2-1 is responsible for a subset of gene expression changes induced by *Foxa1/2* deletion

NKX2-1 has been implicated in the activation of both AT2 and AT1 cell fates and regulates a distinct set of genes in each cell type (Little *et al*., 2019; Little *et al*., 2021). We therefore decided to investigate whether changes in NKX2-1 activity might drive the changes in pulmonary identity programs caused by *Foxa1/2* deletion.

We first sought to determine which gene expression changes induced by *Foxa1/2* deletion can be attributed to NKX2-1 transcriptional activity. In order to answer this question, we performed RNAseq on organoid cultures transduced with shNKX2-1 or shScramble after treatment with 4-OHT or vehicle (n=2 biological replicates, Figure 5A). Intersection of differentially expressed genes from each of the 4 conditions (Figure 5B; Supplemental Table 6) reveals that NKX2-1 is required for ∼25% of gene expression changes caused by *Foxa1/2* deletion. Specifically, 22% of the genes induced by *Foxa1/2* deletion (86/396) and 31% of genes that decline after *Foxa1/2* deletion are NKX2-1-dependent (208/663; Figure 5C).

**Figure 5:**
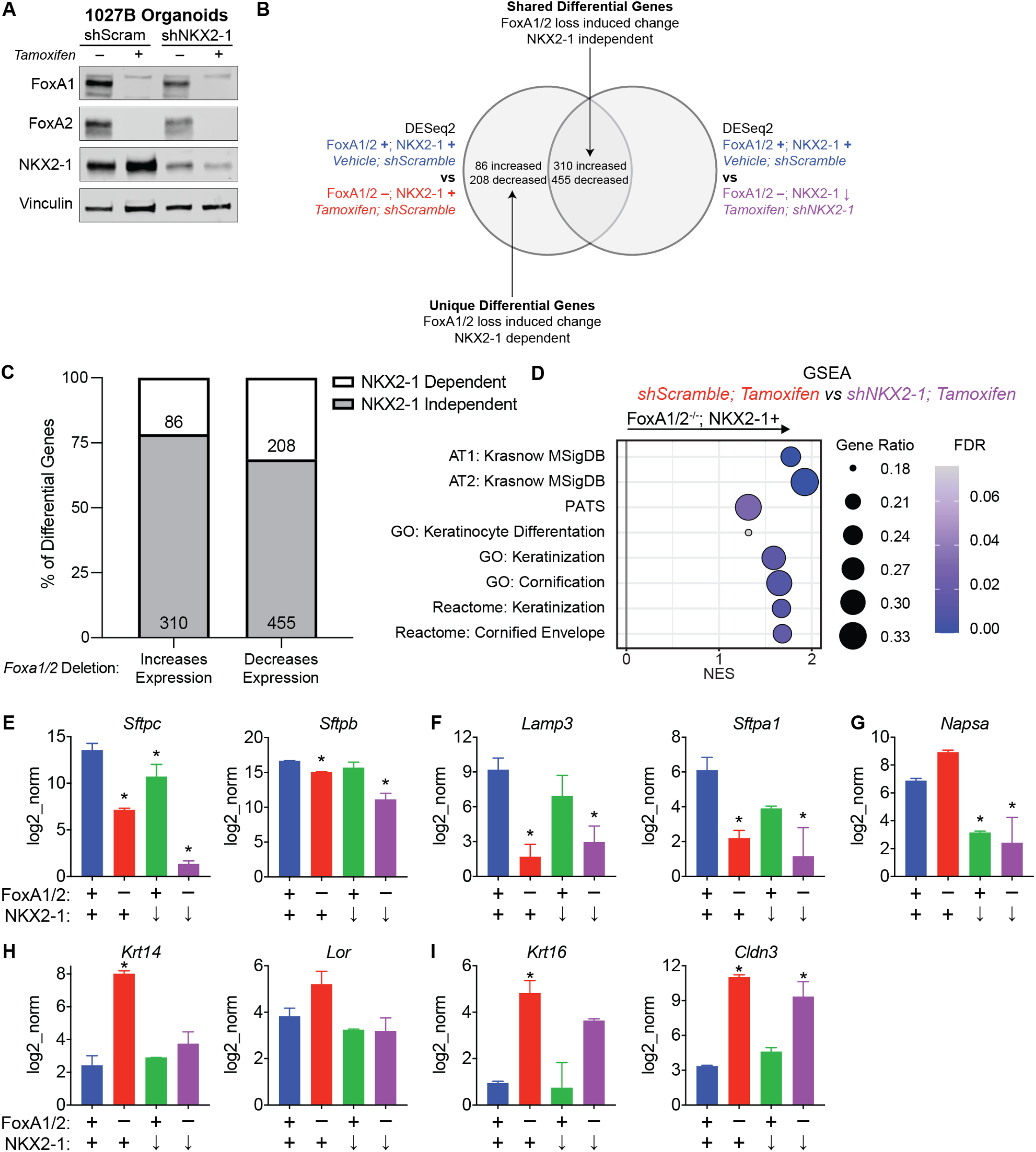
NKX2-1 is required for a subset of gene expression changes induced by FoxA1/2 deletion. A) Western blot of 1027B organoids transduced with shScramble and shNKX2-1 and then treated with 4-OHT/vehicle demonstrating successful knock down of NKX2-1. B) Graphical representation of relevant comparisons for data shown in Figure 5C. See methods section for detailed description of bioinformatic analysis. C) Number of genes increased or decreased following *Foxa1/2* deletion further categorized by NKX2-1 dependence. D) GSEA intersecting relevant gene sets with the ranked gene list generated by differential expression analysis (DESeq2) comparing FoxA1/2-negative; NKX2-1-positive and FoxA1/2-negative; NKX2-1-low organoid cultures. Positive normalized enrichment score (NES) shows enrichment in the NKX2-1-positive cells, indicating that those gene sets are functionally dependent on NKX2-1 in a FoxA1/2-negative context. E) Normalized Log2 expression values as determined by DESeq2 for AT2 genes that show additive FoxA1/2 and NKX2-1 function: *Sftpc* and *Sftpb* (decrease relative to shScramble; vehicle *padj<0.007) F) Normalized Log2 expression values as determined by DESeq2 for AT2 genes primarily dependent on FoxA1/2 function: *Lamp3* and *Sftpa1* (decrease relative to shScramble; vehicle *padj<2×10^-5^) G) Normalized Log2 expression values as determined by DESeq2 for *Napsa,* an AT2 gene that is dependent on NKX2-1 and not FoxA1/2 (decrease relative to shScramble; vehicle *padj<7.22×10^-10^) H) Normalized Log2 expression values as determined by DESeq2 for SSSE genes that show dependence on NKX2-1: *Krt14* and *Lor* (increase relative to shScramble; vehicle *padj<7.22×10^-17^) I) Normalized Log2 expression values as determined by DESeq2 for SSSE genes that are independent of NKX2-1: *Krt16* and *Cldn3* (increase relative to shScramble; vehicle *padj<0.006)

We then investigated how NKX2-1 contributes to the activation of the AT1, AT2, and PATS identities at the program level. All three alveolar transcriptional signatures are enriched in the FoxA1/2-negative/NKX2-1-positive organoids in comparison to the FoxA1/2-negative/NKX2-1-knock-down organoids. This indicates that overall, NKX2-1 is important for both the maintenance of residual AT2 identity and the activation of the de novo AT1 and PATS phenotypes after *Foxa1/2* deletion (Figure 5D). At the individual gene level, we observed nuanced transcriptional dynamics of key AT2 markers. The expression of hallmark surfactant proteins critical for AT2 function, *Sftpc* and *Sftpb,* are regulated by both FoxA1/2 and NKX2-1 in an additive manner. Deletion of *Foxa1/2* and knock down of NKX2-1 independently decrease the expression of these genes, but a combination of both of these conditions yields the maximal decrease in transcription (Figure 5E). In contrast, expression of *Lamp3* and *Sftpa1* is maximally decreased upon *Foxa1/2* deletion, and *Nkx2-1* knockdown inhibits their expression only in FoxA1/2-positive cells, but not FoxA1/2-negative cells (Figure 5F). These data indicate that NKX2-1 can partially activate some, but not all, AT2 marker genes in the absence of FoxA1/2. Conversely, *Napsa* expression exhibits dependence solely on NKX2-1. Loss of FoxA1/2 does not decrease expression, and NKX2-1 knockdown decreases *Napsa* transcription to the same extent in both FoxA1/2-positive and FoxA1/2-negative cells (Figure 5G). The complex transcriptional hierarchy between these transcription factors at AT2 marker genes explains why deleting *Foxa1/2* decreases, but does not completely ablate, the AT2 signature in NKX2-1-positive LUAD. In addition to investigating pneumocyte cell fate regulation, we also asked whether NKX2-1 contributes to the activation of the SSSE in the absence of FoxA1/2. We found that SSSE gene sets are also enriched in the FoxA1/2-negative/NKX2-1-positive organoids when compared to the FoxA1/2-negative/NKX2-1-knock-down organoids, indicating that NKX2-1 is important for full activation of the SSSE phenotype induced by *Foxa1/2* deletion (Figure 5D). At the individual gene level, a subset of SSSE genes are NKX2-1-dependent (e.g. *Krt14* and *Lor*, Figure 5H), whereas others are NKX2-1 independent (e.g. *Krt16* and *Cldn3,* Figure 5I). These data show that NKX2-1 contributes to, but is only partially responsible for, activation of the SSSE identity induced by *Foxa1/2* deletion, suggesting that there are other factors that coordinate with NKX2-1 for full expression of the observed SSSE transcriptional program.

### FoxA1/2 regulate cell type-specific NKX2-1 binding in lung adenocarcinoma

To further elucidate the mechanism by which NKX2-1 drives unique transcriptional programs after *Foxa1/2* deletion, we performed Chromatin Immunoprecipitation followed by massively parallel DNA sequencing (ChIPseq) for NKX2-1 in FoxA1/2-positive and FoxA1/2-negative organoids (n=4 biological replicates of each condition). We identified a total of 36,958 individual NKX2-1 peaks: 16,911 peaks in FoxA1/2-positive organoids and 32,358 peaks in FoxA1/2-negative organoids via MACS2 (Supplemental Table 7). Initial HOMER transcription factor motif analysis of these two sets of peaks showed that, as expected, NKX binding motifs are enriched in both conditions, whereas FOX motifs are exclusively enriched in FoxA1/2-positive NKX2-1 ChIP peaks (Supplemental Figure 6A). Both sets of peaks also share similar genomic occupancy profiles, with a plurality of peaks found in distal intergenic (enhancer) regions, followed by promoters and introns (Supplemental Figure 6B). We also performed ChIPseq for FoxA2 in FoxA1/2-positive organoids (n=4 biological replicates) and identified 6770 individual FoxA2 peaks. We performed HOMER transcription factor motif analysis of FoxA2 peaks and found enrichment for FOX binding motifs, as well as HNF4 and NKX, FOS/JUN, ETS, KLF, and GATA family transcription factors (Supplemental Figure 6C).

Recent publications have shown that in addition to club cells and AT2 cells, NKX2-1 is also expressed in normal AT1 cells, and that differential NKX2-1 localization and transcriptional activity drives alveolar cell fate(Little *et al*., 2019; Little *et al*., 2021). NKX2-1 is expressed in FoxA1/2-negative tumors (Figure 1A), and FoxA1/2-negative tumors retain residual expression of an AT2 identity while also activating an AT1 program (Figure 4A-F). We therefore investigated whether differential NKX2-1 chromatin binding drives a shift in pulmonary cell identity in the absence of FoxA1/2. (Little *et al*., 2021) performed NKX2-1 ChIPseq in normal mouse AT1 and AT2 cells identified AT1-specific and AT2-specific NKX2-1 binding sites. In alignment with our gene expression data, there is an overall increase in NKX2-1 localization to its AT1-specific binding sites in the absence of FoxA1/2 (Figure 6A), whereas global NKX2-1 localization at AT2-specific sites is unchanged (Figure 6B). Together these data suggest that FoxA1/2 regulate NKX2-1 genomic occupancy, which in turn dictates the specific pulmonary identity adopted by tumor cells.

**Figure 6:**
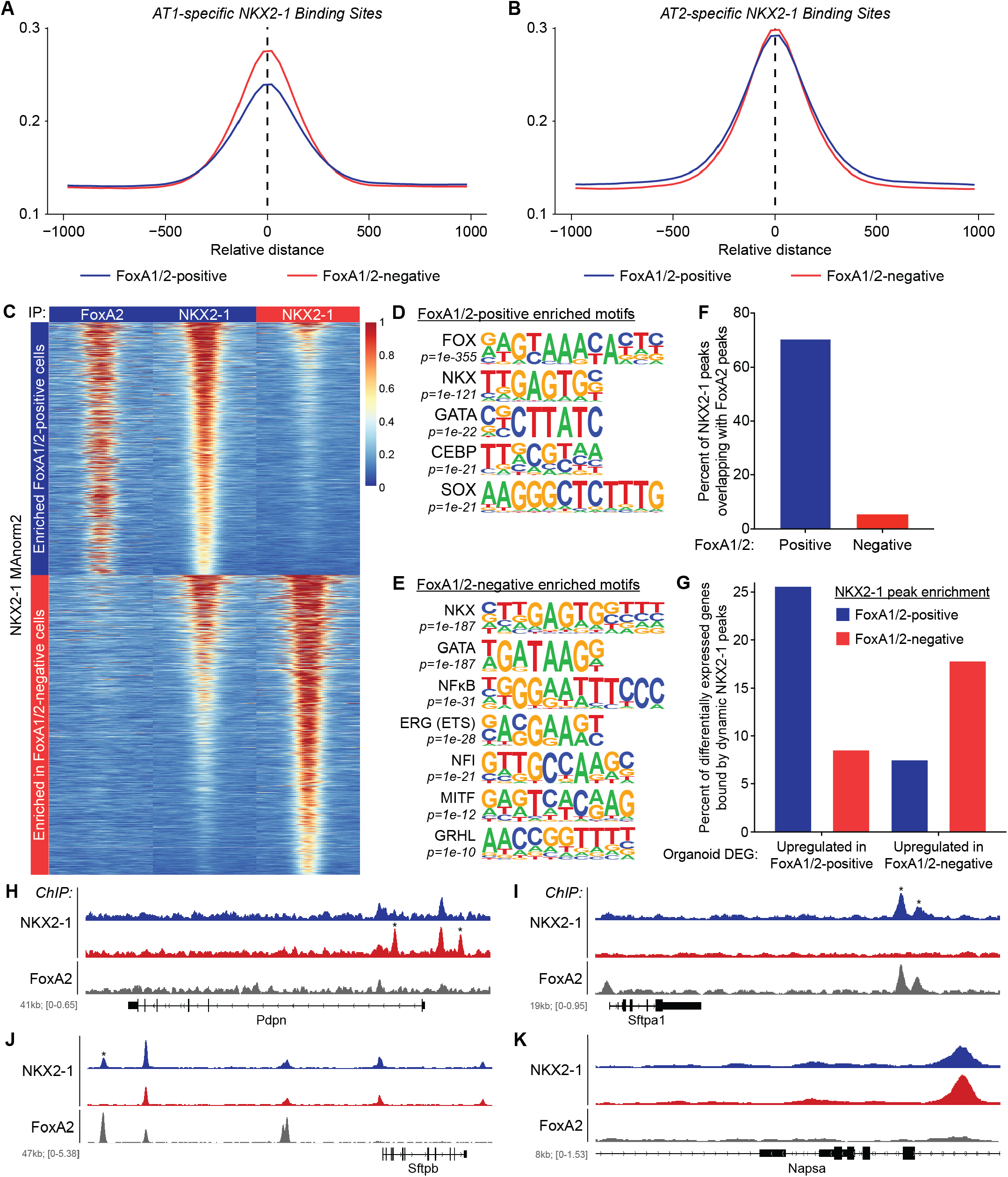
FoxA1/2 regulate global NKX2-1 binding in lung adenocarcinoma. A-B) Mean peak profile of FoxA1/2-positive and FoxA1/2-negative NKX2-1 ChIP peaks (fragments) over all A) AT1-specific and B) AT2-specific NKX2-1 binding sites as defined by NKX2-1 ChIPseq in healthy adult mouse AT1 and AT2 lung cells^23^. Representative of four biological replicates of NKX2-1 ChIPseq in FoxA1/2-positive and FoxA1/2-negative LUAD organoids (1027B). C) Heatmap representing FoxA2 peaks (left panels) and NKX2-1 peaks at the loci of of 2,758 differential NKX2-1 binding sites as determined by MAnorm2 in FoxA1/2-positive (center panels) and FoxA1/2-negative (right panels) LUAD organoids. 1,262 NKX2-1 peaks enriched in a FoxA1/2-positive context (top panels) and 1,496 NKX2-1 peaks enriched in a FoxA1/2-negative context (bottom panels) with a significance cutoff of padj<0.05. D-E) Select transcription factor motifs detected in NKX2-1 peaks enriched in FoxA1/2-positive (D) or (E) FoxA1/2-negative organoids. F) Quantification of NKX2-1 peaks in FoxA1/2-positive and FoxA1/2-negative organoids that overlap with a FoxA2 peak in FoxA1/2-positive organoids. G) Graphical representation of significant association between differential NKX2-1 binding and differential gene expression (p<0.0001, Fishers exact test). Differential NKX2-1 peaks were assigned to protein-coding genes and then intersected with genes significantly changed after *Foxa1/2* deletion as determined by RNAseq of two biological replicates of isogenic 1027B organoids. 25.7% (170/662) of genes enriched in FoxA1/2-positive 1027B organoids are associated with a FoxA1/2-positive enriched NKX2-1 binding site. 8.6% (57/662) of genes enriched in FoxA1/2-positive 1027B organoids are associated with a FoxA1/2-negative enriched NKX2-1 binding site. 7.6% (30/396) of genes enriched in FoxA1/2-negative 1027B organoids are associated with a FoxA1/2-positive enriched NKX2-1 binding site. 18.0% (71/396) of genes enriched in FoxA1/2-negative 1027B organoids are associated with a FoxA1/2-negative enriched NKX2-1 binding site. H-M) FoxA2 and NKX2-1 ChIP peaks in FoxA1/2-positive and FoxA1/2-negative organoids for AT1 marker *Pdpn* (H) and AT2 markers Sftpa1 (I), *Sftpb* (J), and *Napsa* (K). Scale for each gene indicated in individual figures.

Because FoxA2 is a pioneer factor that promotes AT2 identity in LUAD, we asked whether FoxA2 binding correlates with cell type-specific chromatin accessibility in the lung. We performed differential chromatin accessibility analysis on published ATAC-seq data from purified AT1 and AT2 cells (Little *et al*., 2021) to identify differential vs. shared open chromatin regions. We intersected our FoxA2 peaks with these open chromatin regions, and found that 56% of FoxA2 peaks are found in AT2-specific open chromatin regions. Conversely, only 4% of FoxA2 peaks are located within AT1-specific open chromatin regions. Of the remaining FoxA2 peaks, 13% are found within open chromatin regions shared between AT1 and AT2 cells, and 26% are within regions not identified as accessible in either AT1 or AT2 cells (Supplemental Figure 6D).

To investigate the impact of FoxA1/2 loss on NKX2-1 genomic localization in more depth, we utilized MAnorm2 (Tu et al., 2021) to identify a total of 2,758 high confidence differential NKX2-1 binding sites: 1,262 in FoxA1/2-positive cells and 1,496 in FoxA1/2-negative cells (Figure 6C; Supplemental Table 7). When compared to common peaks, a smaller percentage of differential peaks are found in promoters, which is accompanied by an increase to >50% of peaks localized to distal intergenic (enhancer) regions (Supplemental Figure 6B). We next intersected differential NKX2-1 peaks with FoxA2 peaks. There is striking colocalization between FoxA2 and NKX2-1 in FoxA1/2-positive cells; 71% of differential NKX2-1 peaks in FoxA1/2-positive cells overlap with FoxA2 peaks. Conversely, only 5% of differential NKX2-1 peaks in FoxA1/2-negative cells overlap with FoxA2 peaks from control cells (Figure 6C, F).

HOMER analysis (Supplemental Table 7) of differential (Figure 6D-E) and common peaks (Supplemental Figure 6E) identified two interesting patterns beyond the expected FOX motif and NKX motif distribution. (1) Differential AT2 transcription factor distribution: CEBP-family proteins, NFI-family proteins, and GRHL-family proteins are all transcription factors that play a role in AT2 differentiation (Bachurski et al., 2003; Herriges et al., 2012; Little *et al*., 2021; Varma et al., 2012). CEBP motifs are enriched in FoxA1/2-positive and static NKX2-1 peaks, but not in FoxA1/2-negative NKX2-1 peaks. Instead, we find alternative AT2 transcription factor motifs (NFI and GRHL) in a FoxA1/2-negative context. These data suggest that *Foxa1/2* deletion could alter the specific AT2-regulating transcription factors with which NKX2-1 physically interacts and/or binds chromatin. (2) NFκB enrichment in FoxA1/2-negative NKX2-1 ChIP peaks: activation of the NFκB pathway is a hallmark of the PATS/ADI cell population that emerges in response to alveolar injury (Kobayashi *et al*., 2020; Strunz *et al*., 2020). The presence of an NFκB motif in *Foxa1/2* deleted cells raises the possibility of cooperation between NKX2-1 and NFκB signaling in promoting a ADI-like identity.

We also intersected FoxA2 peaks and differential NKX2-1 peaks with genes that significantly change in response to *Foxa1/2* deletion. FoxA2 peaks are associated with 12.5% of differentially expressed genes (12.7% of genes enriched in FoxA1/2-positive cells and 12.1% of genes enriched in FoxA1/2-negative cells). Notably, we find that direct binding by differential NKX2-1 peaks is more significantly associated with gene activation than repression (p<0.0001, Fishers Exact Test; Figure 6G; Supplemental Table 7). Specifically, genes upregulated in FoxA1/2-negative organoids are more strongly associated with NKX2-1 peaks specific to FoxA1/2-negative organoids (∼18% bound) than with peaks specific to FoxA1/2-positive organoids (∼7% bound). For example, there are two unique NKX2-1 peaks in *Foxa1/2* deleted cells within the promoter region of the AT1 marker Podoplanin (*Pdpn*; Figure 6H). Similarly, genes differentially expressed in FoxA1/2-positive organoids are more strongly associated with NKX2-1 peaks specific to FoxA1/2-positive organoids, including a subset of AT2 marker genes. Although there is no global change in NKX2-1 localization to unique AT2 binding sites (Figure 6B), there are specific AT2 marker genes that require FoxA1/2 for NKX2-1 localization. For example, two NKX2-1 peaks in the promoter of *Sftpa1*, which co-localize with FoxA2, are only present in FoxA1/2-positive cells (Figure 6I). Corresponding with the loss of NKX2-1 peaks, *Foxa1/2* deletion is sufficient for maximal reduction in *Sftpa1* transcript levels (Figure 5F). In contrast, we find both differential and common NKX2-1 peaks near *Sftpb*, regardless of the presence or absence of an overlapping FoxA2 peak (Figure 6J). This likely explains why NKX2-1 can partially activate *Sftpb* transcription in the absence of FoxA1/2 (Figure 5E). Expression of the AT2 marker *Napsa* is not impacted by *Foxa1/2* deletion and does not have a FoxA2 peak, but expression does decrease upon NKX2-1 knock down. Correspondingly, NKX2-1 binds *Napsa* independent of FoxA1/2 status (Figure 5G, 6K).

Taken together, these data underscore the complex transcriptional relationship between FoxA1/2 and NKX2-1 in regulation of pulmonary identity. FoxA1/2 regulate cell type-specific localization of NKX2-1 in lung adenocarcinoma, and the differential NKX2-1 binding sites observed in the context of *Foxa1/2* deletion are correlated with differential gene expression leading to the activation of AT1 and AT2 cellular identities.

### NKX2-1 impairs tumor growth in the absence of FoxA1/2

After we established the role of NKX2-1 in regulation of LUAD cellular identity changes following *Foxa1/2* deletion, we next wanted to determine whether de novo NKX2-1 transcriptional activity and genomic localization contribute to the growth arrest caused by FoxA1/2 loss. Stochastic NKX2-1 loss can facilitate progression of KP LUAD (Winslow *et al*., 2011). We therefore predicted that if NKX2-1 restrains the growth of *Foxa1/2*-deleted tumors, there would also be selection for NKX2-1 loss in *Foxa1/2*-deleted tumors over time. Histopathologic analysis of tamoxifen treated mice in the survival study (Figure 1D) revealed two classes of tumors that grew to macroscopic size despite the absence of FoxA1/2. These macroscopic tumors exhibited morphologies that are highly distinct from the majority of FoxA1/2-negative tumors, which remained microscopic and were indistinguishable from FoxA1/2-negative tumors evaluated two weeks after deletion (Figure 1A-C). The first class, found in 80% of tamoxifen treated KPF_1_F_2_ mice (n=8/10 mice histologically evaluated), is characterized by a keratinizing squamous cell carcinoma (SCC) morphology (Supplemental Table 1). These SCCs express squamous lineage specifier ΔNp63 and retain expression of NKX2-1, albeit at reduced levels compared to adenocarcinomas (Supplemental Figure 7A). The second class, found in 30% of tamoxifen treated KPF_1_F_2_ mice (n=3/10), consists of poorly differentiated, high grade tumors that are NKX2-1-negative (Figure 7A). In contrast, all poorly-differentiated, NKX2-1-negative tumors observed in vehicle treated KPF_1_F_2_ mice (n=9/10) expressed FoxA1 and/or FoxA2, suggesting that the combined absence of NKX2-1 and FoxA1/2 is specifically found in the context of *Foxa1/2* genetic deletion. Because *Foxa1/2* deletion does not directly lead to loss of NKX2-1 or gain of ΔNp63 expression (Figure 1A and Supplemental Figure 4E), these observations point to specific stochastic events that might enable LUAD to escape *Foxa1/2* deletion over time. Consistent with the first possibility, a stochastic decrease in NKX2-1 levels correlated with sarcomatoid morphology and eventual outgrowth of subcutaneous tumors following *Foxa1/2* deletion (Supplemental Figure 2B, right column).

**Figure 7:**
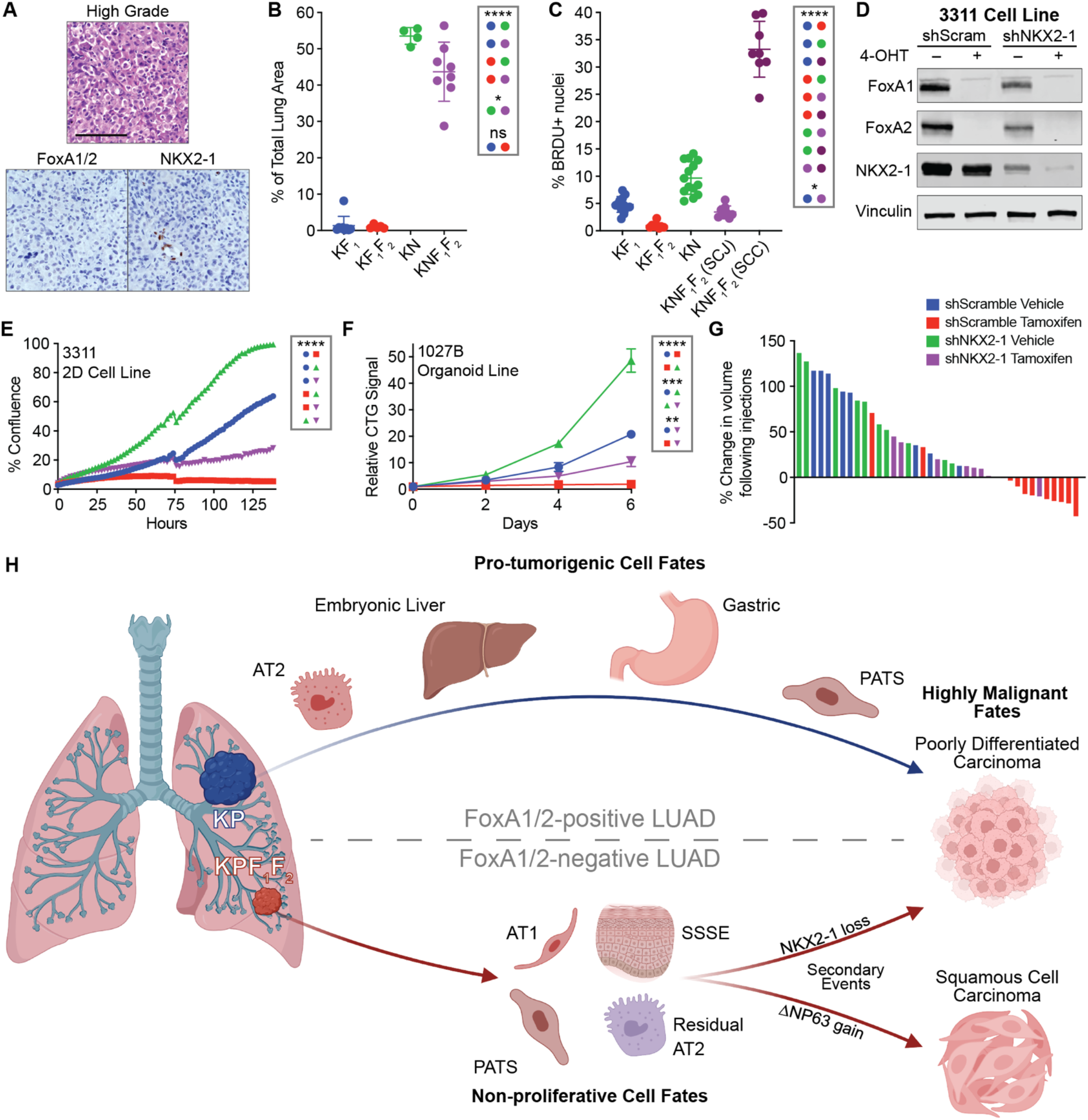
NKX2-1 restrains tumor growth in the absence of FoxA1/2. A) Representative images of high grade, NKX2-1-negative tumors that emerged in the FoxA1/2-negative cohort of long-term survival studies. Top panel H&E, bottom panels IHC (scale bar 250 μm). B) Tumor burden quantification of KF_1_, KF_1_F_2_, KN, and KNF_1_F_2_ tumors initiated with Ad5mSPC-Flp, injected with tamoxifen 6 weeks later, and sacrificed at 12 weeks (multiple unpaired t-tests; *p=0.0424; ****p<0.0001). C) BRDU IHC quantification experiment shown in B (multiple unpaired t-tests; *p=0.0366; ****p<0.0001) D) Western blot of protein extract from a KPF_1_F_2_ cell line (3311) transduced with shScramble and shNKX2-1 collected 3 days post 48-hour 4-OHT/vehicle treatment demonstrating successful knock down of NKX2-1. E) Incucyte-based confluency growth assay of cells shown in D one week after 4-OHT/vehicle treatment. One representative experiment shown of two biological replicates with 8 technical replicates each (multiple unpaired t-tests of endpoint values; ****p<0.0001 for all comparisons). F) Timecourse experiment using the CellTiter-Glo 3D cell viability assay of cells shown in Figure 5A. CTG assay was performed on the day of seeding, as well as 2, 4, and 6 days post-seeding. Data shown relative to day 0 luminescence reads for each condition. One representative experiment shown of two biological replicates with 3 technical replicates each (multiple unpaired t-tests of endpoint values; **p=0.0016; ***p<0.0005; ****p<0.0001). G) Waterfall plot showing the change in individual 3311 subcutaneous tumor volumes. Percent change representing the first measurement after completion of tamoxifen/vehicle injections relative to the measurement immediately preceding treatment. H) FoxA1/2 are required for the activation of the dual identity, combined pulmonary and gastro-intestinal identity that is a key step in the malignant evolution of this disease. In the absence of FoxA1/2, LUAD tumors shed a portion of their AT2 identity while activating an AT1-like identity and maintaining a PATS/ADI pulmonary state. Discrete subpopulations within both of these pulmonary states also express gene sets associated specifically with the superficial layers of the stratified squamous epithelium. We have shown that NKX2-1 acts in a tumor-suppressive manner in a FoxA1/2-negative context. In the absence of FoxA1/2, NKX2-1 localization dynamically changes, and these alterations are correlated with global gene expression changes. We have shown that NKX2-1 transcriptional activity is at least partially responsible for the gene expression changes following FoxA1/2 deletion, including both the emergence of AT1 and SSSE identities as well as a decrease in AT2 identity. We speculate that this transition to non-proliferative cellular identities upon *Foxa1/2* deletion, as aided by de novo NKX2-1 activity, plays a major role in the growth inhibition observed in FoxA1/2-negative LUAD. Over time, FoxA1/2-negative lesions can overcome this growth inhibition by either progressing into a fully malignant, keratinized squamous tumor or stochastically downregulating NKX2-1 expression and transforming into a high grade, poorly-differentiated tumor (Image created with Biorender).

Based on these correlations, we sought to determine whether loss of NKX2-1 is sufficient to rescue *Foxa1/2* deletion using multiple independent approaches. First, we initiated *Kras^G12D^*-driven p53-proficient tumors, allowed tumors to develop for 6 weeks, and then administered tamoxifen to delete *Foxa1* alone (KF_1_), *Foxa1* and *Foxa2* (KF_1_F_2_), *Nkx2-1* alone (KN), or *Nkx2-1, Foxa1* and *Foxa2* (KNF_1_F_2_). Tumors progressed for an additional 6 weeks before tissue was collected. *Nkx2-1* deletion significantly rescued the effects of *Foxa1/2* deletion on tumor growth, resulting in a comparable overall tumor burden in KNF_1_F_2_ and KN mice (Figure 7B). As expected (Camolotto *et al*., 2018; Snyder *et al*., 2013), tumor burden in KN was highest overall. We have previously shown that KNF_1_F_2_ mice develop two distinct tumor types (squamocolumnar junction (SCJ)-like and keratinizing squamous cell carcinoma (SCC), also pictured in Supplemental Figure 7B)(Camolotto *et al*., 2018). Both subtypes are more proliferative than both KF_1_ and KF_1_F_2_ tumors and therefore contribute to the overall increase in tumor burden of KNF_1_F_2_ mice (tumor burden of individual subtypes quantified in Supplemental Figure 7C). SCC lesions have a higher BrdU incorporation rate than SCJ-like lesions, consistent with the fact that SCC can form macroscopic tumors whereas SCJ-like lesions remain microscopic in size (Figure 7C).

To determine whether NKX2-1 loss can also rescue *Foxa1/2* deletion in a p53-deficient context, we introduced an shRNA against NKX2-1(Winslow *et al*., 2011) to KPF_1_F_2_ cell lines and organoids in order to decrease NKX2-1 levels alongside deletion of *Foxa1/2* via 4-OHT treatment (Figure 7D; Supplemental Figure 5A). Knockdown of NKX2-1 partially rescues the proliferation defect observed upon *Foxa1/2* deletion in both 2D cell lines (Figure 7E) and organoid cultures (Figure 7F). As expected from previous work(Snyder *et al*., 2013; Winslow *et al*., 2011), NKX2-1 knockdown also enhances proliferation in parental FoxA1/2-positive cell lines and organoids. In order to determine whether NKX2-1 knockdown prevents tumor regression upon FoxA1/2 loss, we subcutaneously injected NSG mice with the 3311 cell line carrying shNKX2-1 and treated mice with either vehicle or tamoxifen when tumors reached an average volume of 200mm^3^. We found that NKX2-1 knockdown increases the rate of growth in both the tamoxifen and vehicle treatments. Importantly, NKX2-1 knock down largely prevents the tumor regression following *Foxa1/2* deletion seen in the shScramble tamoxifen treated cohort (Figures 2D and 7G, Supplemental Figure 7D-E). Consistent with these rescue experiments, CRISPR-Cas9 mediated *FOXA1/2* deletion in two NKX2-1-negative human LUAD cell lines (A549 and NCI-H2122) has no significant effect on growth (Supplemental Figure 7F-G). Together, these data demonstrate that induction of aberrant NKX2-1 activity is one mechanism by which FoxA1/2 loss inhibits LUAD growth.

## DISCUSSION

LUAD progression is driven by epigenetic changes that lead to increased plasticity relative to normal cells of origin, which ultimately enables cancer cells to undergo profound changes in cell identity that enhance malignant potential (Boumahdi and de Sauvage, 2020; Campos-Parra *et al*., 2014; LaFave *et al*., 2020; Le Magnen *et al*., 2018; Marjanovic *et al*., 2020; Niederst *et al*., 2015; Quintanal-Villalonga *et al*., 2020; Russell, 2011; Sun and Yang, 2006; Tavernari *et al*., 2021; Travis *et al*., 2011). Here we show that FoxA1/2 regulate critical aspects of lineage plasticity and infidelity, coordinately driving multiple cell identity programs that are essential for LUAD progression (Figure 7H, top). In the absence of FoxA1/2, LUAD cells undergo lineage switching to alternative identities normally associated with low levels of proliferation (AT1 and SSSE). However, a subset of FoxA1/2-negative LUAD cells stochastically escape this proliferative arrest over time, undergoing an additional lineage switch (to either SCC or poorly differentiated, NKX2-1-negative carcinoma) that enables them to regain their full malignant potential (Figure 7H, bottom).

### FoxA1/2 can activate pulmonary and non-pulmonary identities within a single cell in lung adenocarcinoma

In NKX2-1-positive, p53-deficient LUAD, FoxA1/2 are required for expression of multiple cellular identity programs, including an AT2-like program and programs associated with endodermal/GI-like differentiation states. Moreover, we find that FoxA1/2 can activate both AT2 and GI-like differentiation programs within the same cancer cell. This was particularly striking because our previous work in p53-proficient LUAD mouse models showed that FoxA1/2 are restricted to pulmonary activity when NKX2-1 is expressed, and can only activate GI programs upon NKX2-1 loss (Snyder *et al*., 2013). Specific mechanisms that promote expanded FoxA1/2 transcriptional activity in p53-deficient LUAD are unknown and will require additional investigation. It is likely that a combination of altered histone modification and DNA methylation profiles, as well as changes in cofactor relationships, could enable FoxA1/2 to access non-pulmonary targets in KP LUAD that it cannot bind in p53-proficient LUAD (or normal lung epithelium). Moreover, we have recently found that oncogenic signaling can modulate cellular identity in NKX2-1-negative LUAD (Zewdu et al., 2021). We speculate that the increased signaling downstream of KRAS^G12D^ observed during progression in KP LUAD (Feldser et al., 2010) may also contribute to the capacity of FoxA1/2 to activate GI-related differentiation programs despite the expression of NKX2-1.

### FoxA1/2 promote LUAD growth by multiple mechanisms, including regulation of NKX2-1 activity

NKX2-1-positive LUAD growth is highly dependent on FoxA1/2. Deletion of *Foxa1/2* in autochthonous lung tumors, murine organoids, and human LUAD cell lines severely impedes proliferation, which in turn significantly increases survival. This intense dependence on FoxA1/2 is likely due to a combination of at least two distinct mechanisms, the first being the loss of the dual identity state associated with progression into high grade disease. Additional studies will be needed to determine how co-activation of GI and pulmonary programs in NKX2-1-positive disease promotes tumor progression. However, we have previously shown that HNF4α, which is a direct FoxA1/2 target in non-pulmonary tissues, promotes the initiation of p53-proficient, NKX2-1-negative LUAD (Snyder *et al*., 2013). This raises the possibility that the growth of p53-deficient, NKX2-1-positive LUAD might also be augmented by HNF4α and other endodermal lineage specifiers, many of which may be direct FoxA1/2 targets outside the lung.

Second, our data suggest that FoxA1/2-dependence in LUAD is also due to novel NKX2-1 activity that, in the absence of FoxA1/2, triggers the activation of transcriptional programs associated with non-proliferative cell types, including AT1 cells and maturing cells of the superficial stratified squamous epithelium. In KP LUAD, FoxA1/2 regulate NKX2-1 localization, resulting in the expression of AT2-specific targets. Upon *Foxa1/2* deletion, NKX2-1 exhibits de novo transcriptional activity that contributes to the activation of AT1 and SSSE identities. Under normal conditions, both of these cell types are non-proliferative, and typically require pools of other cell types to differentiate and replace them upon injury or death (AT2/PATS cells and basal epithelial stem cells, respectively). In further support of this hypothesis, concomitant loss of NKX2-1 alongside FoxA1/2 prevents the significant antiproliferative impact of *Foxa1/2* deletion alone, as well as the emergence of these non-proliferative cellular identities. These data suggest a model in which novel NKX2-1 transcriptional activity after FoxA1/2 loss restrains tumor growth through the activation of alternative, non-proliferative cell identity programs. Nevertheless, there are likely to be additional mechanisms by which de novo NKX2-1 activity restrains LUAD growth following *Foxa1/2* deletion. One additional mechanism may be induction of “oncogene overdose” by de novo NKX2-1 binding that impairs growth. *Ror1* is induced by *Foxa1/2* deletion and exhibits de novo NKX2-1 binding in FoxA1/2-null organoids. In EGFR mutant LUAD, *Ror1* is a putative NKX2-1 target and mediates receptor tyrosine kinase (RTK) signaling (Yamaguchi et al., 2012). High level RTK signaling can inhibit the growth of KRAS-driven LUAD by hyperactivation of oncogenic pathways (Unni et al., 2015). Defining the mechanism(s) by which NKX2-1 actively inhibits growth following FoxA1/2 loss is an important direction for future work.

### FoxA1/2-negative tumors can overcome growth arrest via additional lineage switching

Despite the severe antiproliferative consequences of *Foxa1/2* deletion, we have identified two categories of tumors that stochastically escape this fate over extended periods of time: fully keratinized, NKX2-1 positive SCCs or high grade, poorly differentiated NKX2-1 negative carcinomas. This raises the question of whether all *Foxa1/2*-deleted cells are equally capable of progressing to these two fates, or if only a subset of cells are poised to undergo the stochastic lineage switch required to resume proliferation in a FoxA1/2-negative state. Although our data cannot answer this question directly, we speculate that cells within Cluster 7 may be the most poised to overcome *Foxa1/2* deletion-induced growth arrest over time. These cells retain expression of the ‘Highly Mixed’ transcriptional program, which is associated with increased tumorigenic potential in KP LUAD (Marjanovic *et al*., 2020) and might be permissive for dedifferentiation into the NKX2-1 negative, poorly differentiated carcinoma. Furthermore, a subgroup of cells within Cluster 7 adopted an SSSE-like transcriptional identity. Those cells that co-express SSSE and ‘Highly Mixed’ programs may be the most likely to give rise to the fully keratinized squamous cell carcinoma escaper tumors in response to stochastic upregulation of ΔNp63. Technical innovations enabling *Foxa1/2* deletion in specific subpopulations of KP LUAD will help to address this question in the future.

### Translational implications of FoxA1/2 dependency in NKX2-1-positive LUAD

In this publication we have laid the groundwork for several future directions. (1) This work clearly shows that FoxA1/2 are potential therapeutic targets. Whether it be through targeted degradation or the development of drugs that selectively inhibit FoxA proteins, our data show that blocking their transcriptional activity could improve outcomes for LUAD patients with NKX2-1-positive disease. (2) Since we have also shown that tumors can develop resistance to FoxA1/2 inhibition, it will be important to develop combinatorial strategies. We have shown that NKX2-1 actively inhibits growth following Foxa1/2-deletion in LUAD. It will be important to define the mechanism(s) by which NKX2-1 restrains tumor progression and ultimately identify therapeutic agents that might further augment the anti-tumor effects of NKX2-1 after FoxA1/2 inhibition. (3) We have identified immune cell populations unique to FoxA1/2-negative tumors (such as M1-polarized alveolar macrophages) that may contribute to growth arrest. If functional studies demonstrate a role for these cells in producing an anti-tumor inflammatory response, it would be valuable to test candidate therapeutic strategies that might further augment the anti-tumor response of the TME (for example, by further promoting M1 macrophage polarization (Wang et al., 2021)). (4) Our work does not address the potential impact of FoxA1/2 inhibition on normal adult tissues, so it is also important to consider FoxA1/2 targets that activate GI transcriptional programs critical for tumor progression. Investigating these downstream targets, some of which may be directly druggable, in enforcing a dual-identity state is an important future direction and could yield a more clinically-actionable target.

Identifying the master transcriptional regulators of cellular identity and investigating the impact of their inactivation is critical for understanding of mechanisms of cancer progression and will contribute to the development of differentiation state-specific therapeutic strategies. Although transcription factors have been historically viewed as undruggable, recent advances in targeted degradation have the potential to make transcription factor inhibition a pharmacologically tractable strategy in cancer(Samarasinghe and Crews, 2021). This study suggests that targeting FoxA1/2 would have a major impact on NKX2-1-positive LUAD. We also identify candidate mechanisms of escape from FoxA1/2 inhibition, which will likely require co-targeting in order to forestall resistance and achieve a durable response to this potential therapeutic approach.

## Acknowledgements

We are grateful to members of the Snyder lab for suggestions and comments. We thank Brian Dalley for his sequencing expertise, Jay Gertz for his ChIPseq expertise, James Marvin for his FACS expertise, Kaylyn Bauer for their immunology expertise, and the University of Utah core facilities (PRR, BMP, DNA sequencing, Genomics/Bioinformatics, Flow Cytometry). The results published here are in part based upon data generated by the TCGA Research Network: https://www.cancer.gov/tcga. ELS was supported grants from NIH (R01CA212415 and R01CA240317), a Career Award for Medical Scientists from the Burroughs Wellcome Fund, and institutional funds (Department of Pathology and Huntsman Cancer Institute/Huntsman Cancer Foundation, University of Utah). CRV was supported by Pershing Square Sohn Cancer Research Alliance, the Cold Spring Harbor Laboratory and Northwell Health Affiliation, the National Cancer Institute 5P01CA013106-Project 4 and 1R01CA229699, the Thompson Family Foundation, the Simons Foundation, and a Career Development Award from the Pancreatic Cancer Action Network–American Association for Cancer Research 16-20-25-VAKO. GO was supported by the NIH (F31CA243427) and the Eunice Kennedy Shriver National Institute of Child Health & Human Development of the National Institutes of Health (T32HD007491). OK was supported by the Deutsche Forschungsgemeinschaft (DFG) Research Fellowship (KL 3228/1-1). Research reported in this publication utilized shared resources (including Flow Cytometry, High Throughput Genomics, Bioinformatics, and Biorepository and Molecular Pathology) at the University of Utah and was supported by the National Cancer Institute of the National Institutes of Health under Award Number P30CA042014. Work in the flow cytometry core was also supported by the National Center for Research Resources of the National Institutes of Health under Award Number 1S20RR026802. The content is solely the responsibility of the authors and does not necessarily represent the official views of the NIH.

## Author contributions

GO and ELS designed experiments. GO, GF, AJ, KG, WO, and RT performed experiments. GO, GF, CS, BL, TP, BTS, and ELS analyzed data. ELS performed histopathologic review. OK, CRV and KK provided essential tools and reagents. GO and ELS wrote the manuscript. All authors discussed results, reviewed and revised the manuscript.

## Supplemental Tables

Table 1: Detailed histological analysis of KP, KPF_1_, KPF_2_, KPF_1_F_2_, and BPF_1_F_2_ survival studies

Table 2: Mouse Cell Atlas analysis of scRNAseq data

Table 3: Differentially expressed genes in KPF_1_F_2_, KPF_1_, and KPF_2_, organoid isogenic pairs and TCGA Pan Cancer Atlas *NKX2-1*-high LUAD samples as determined by DESeq2

Table 4: scRNAseq tumor and stromal cell barcodes, DEG lists, and tumor clusters GSEA

Table 5: Significantly enriched gene sets in KPF_1_F_2_ organoid isogenic pairs as determined by GSEA

Table 6: Differentially expressed genes in KPF_1_F_2_ shScramble and shNKX2-1 organoid isogenic pairs as determined by DESeq2

Table 7: NKX2-1 and FoxA2 ChIPseq (all peaks, differential analysis, gene intersection, HOMER analysis)

## RESOURCE AVAILABILITY

### Lead Contact

Further information and requests for resources and reagents should be directed to and will be fulfilled by the lead contact, Eric Snyder

### Materials availability

Novel murine cell lines and organoids are available upon request. The newly generated pCDH-EFS-FlpO lentivector will be deposited at Addgene.

### Data and code availability

Single-cell RNAseq, bulk RNAseq, and ChIPseq data have been deposited at Gene Expression Omnibus (GSE188438) and are publicly available as of the date of publication. This paper does not report original code. Any additional information required to reanalyze the data reported in this paper is available from the lead contact upon request.

## EXPERIMENTAL MODEL AND SUBJECT DETAILS

### Animal studies

Mice harboring *Kras^FSF-G12D^* ^((Young *et al*., 2011))^, *Braf^FSF-V600E^* ^((Shai et al., 2015))^, *p53^frt^* ^(^(Lee *et al*., 2012)^)^, *Rosa-FSF-Cre^ERT2^* ^(^(Schonhuber *et al*., 2014)^)^*, Foxa1^flox^* ^((Gao^ *^et al.^*^, 2008))^, and *Foxa2^flox^* ^((Sund^ *^et al.^*^, 2000))^ have been previously described. All animals were maintained on a mixed 129/B6 background. All experimental mice were between 2 and 6 months of age at intubation. Mice of both sexes were used throughout each study. Animal studies were approved by the IACUC of the University of Utah, conducted in compliance with the Animal Welfare Act Regulations and other federal statutes relating to animals and experiments involving animals, and adhered to the principles set forth in the Guide for the Care and Use of Laboratory Animals, National Research Council (PHS assurance registration number A-3031-01).

### Cell lines and primary cultures

All primary murine organoid cultures (see Key Resources Table) were established within Matrigel (Corning or Preclinical Research Shared Resource core facility) submerged in recombinant organoid medium for approximately two weeks (Advanced DMEM/F-12 supplemented with 1X B27 (Gibco), 1X N2 (Gibco), 1.25mM nAcetylcysteine (Sigma), 10mM Nicotinamide (Sigma), 10nM Gastrin (Sigma), 100ng/ml EGF (Peprotech), 100ng/ml R-spondin1 (Peprotech), 100ng/ml Noggin (Peprotech), and 100ng/ml FGF10 (Peprotech). After organoids were established, cultures were switched to 50% L-WRN conditioned media(Miyoshi and Stappenbeck, 2013).

A549, 3311, and HEK293T cells were cultured in DMEM/10% FBS (Gibco). NCI-H358, NCI-H441, NCI-H2009, and NCI-H2122 were cultured in RPMI/10% FBS (Gibco). NCI-H1651 was cultured in Advanced DMEM F-12/10% FBS (Gibco). All cell lines were tested periodically for mycoplasma contamination. To maintain cell cultures mycoplasma free, all culture media were supplemented with 2.5 ug/ml Plasmocin.

## METHOD DETAILS

### Tumor initiation, tamoxifen administration, and BrdU administration in vivo

Autochthonous lung tumors were initiated by administering viruses via intratracheal intubation. Adenoviruses were obtained from University of Iowa Viral Vector Core. Lentivirus was generated as described in **Lentiviral production and transduction** methods section.

| **Experiment** | **Virus** | **PFU/mouse** |
| --- | --- | --- |
| Figure 1A-C; Supplemental Figure 1D-E | Adeno-CMV-FlpO | $4 \times 10^7$ |
| Supplemental Figure 1I | | $1 \times 10^7$ |
| Figure 1D-E, 7A; Supplemental Figure 1 G-H, 7A | Lenti-EFS-FlpO | $1 \times 10^4$ |
| Supplemental Figure 1F,J | Adeno-mSPC-FlpO | $2.5 \times 10^8$ |
| Figures 3-4 | | $5 \times 10^8$ |
| Figure 7B-C; Supplemental Figure 7 B-C | | $1 \times 10^9$ |

Tumor-specific activation of Cre^ERT2^ nuclear activity was achieved by intraperitoneal injection of tamoxifen (Sigma) dissolved in corn oil at a dose of 120mg/kg. Mice received 4 injections over the course of 5 days. For survival experiments, mice were additionally given pellets supplemented with 500mg/kg tamoxifen (Envigo) for 7 days following injections. BrdU incorporation was performed by injecting mice at 40mg/kg (Sigma) intraperitoneally 1 hour prior to tissue collection. Mice in survival studies were monitored for lethargy and respiratory distress, at which time animals were euthanized.

### Analysis of human lung adenocarcinoma

#### IHC

Whole sections of de-identified formalin fixed, paraffin-embedded (FFPE) LUAD (n=42) were obtained from the Intermountain Biorepository, which collects samples in accordance with protocols approved by the Intermountain Healthcare Institutional Review Board. An additional 92 de-identified tumors were evaluated by FFPE tissue microarrays obtained from US Biomax (BC04115c and BCS04017).

#### TCGA

We first filtered all TCGA Pan Cancer Atlas lung adenocarcinoma samples based on high *NKX2-1* expression (*NKX2-1* z-score>-0.2, n=402/516 LUAD samples as of May 2022). The 402 *NKX2-1*-high samples were further filtered on *FOXA1* and *FOXA2* expression. We selected patients that had below average expression of both *FOXA1* and *FOXA2* for the *FOXA1/2*-low cohort (z-score<0 for both genes, n=23/402) and patients with above average expression of both *FOXA1* and *FOXA2* for the *FOXA1/2*-high cohort (z-score>0 for both genes; n=114/402). We identified differentially expressed genes between the *FOXA1/2*-high and *FOXA1/2*-low cohorts using DESeq2 and used the resultant log2FCs to run GSEA with our selected gene sets.

### Histology and immunohistochemistry

All tissues were fixed in 10% formalin overnight and when necessary, lungs were perfused with formalin via the trachea. Organoids were first fixed in 10% formalin overnight and then mounted in HistoGel (Thermo Fisher Scientific). Mounted organoids and tissues were transferred to 70% ethanol, embedded in paraffin, and four-micrometer sections were cut. Immunohistochemistry (IHC) was performed manually on Sequenza slide staining racks (Thermo Fisher Scientific). Sections were treated with Bloxall (Vector Labs) followed by Horse serum 536 (Vector Labs) or Rodent Block M (Biocare Medical), primary antibody, and HRP-polymer-conjugated secondary antibody (anti-Rabbit, Goat and Rat from Vector Labs; anti-Mouse from Biocare. The slides were developed with Impact DAB (Vector Labs) and counterstained with hematoxylin. Slides were stained with antibodies to FoxA1 (1:4000, Abcam 10881-14), FoxA2 (1:1200, Abcam 4466), Murine NKX2-1 (1:2000, Abcam EP1584Y), Human NKX2-1 (1:2000, Abcam 133638), BRDU (1:400, Abcam BU1/75), CC3 (CST 9664S 1:800), proSP-B (1:3000, Millipore AB3430), proSP-C (1:4000, Millipore AB3786), HNF4α (1:500, CST C11F12), ΔNp63 (1:100, Biocare [clone BC28]). Images were taken on a Nikon Eclipse Ni-U microscope with a DS-Ri2 camera and NIS-Elements software. Tumor Burden and BrdU quantitation and histological analyses were performed on hematoxylin and eosin-stained and IHC-stained slides using NIS-Elements software. All histopathologic analysis was performed by a board-certified anatomic pathologist (E.L.S.).

### MicroCT imaging and analysis

Mice were anesthetized with isoflurane and imaged using a small animal Quantum GX2 microCT (Perkin Elmer). Images were acquired with 2 minute scans at 90 kV and 88-µA current, and reconstructed at a 90-µm voxel size. Resulting images were processed with Analyze 12.0 software (Analyze Direct) as described in Mollaoglu et al. 2017(Mollaoglu et al., 2017).

### Establishing primary murine LUAD cell lines and organoids

Five months after tumor initiation in KF_1_F_2_ mice (2D cell lines; Lenti-CA2-FlpO-sh*p53*) and KPF_1_F_2_ mice (3D organoid lines; Adeno-mSPC-FlpO), tumor bearing mice were euthanized and lungs were isolated. Individual macroscopic tumors were removed from lungs, minced under sterile conditions, and digested at 37°C for 30 minutes with continuous agitation in a solution of Advanced DMEM/F12 containing the following enzymes: Collagenase Type I (Thermo Fisher Scientific, 450U/ml), Dispase (Corning, 5U/ml), DNaseI (Sigma, 0.25mg/ml). Enzyme reactions were stopped by addition of cold DMEM/F-12 with 10% FBS. The digested tissue was repeatedly passed through a 20-gauge syringe needle, sequentially dispersed through 100μm, 70 μm, and 40μm cell strainers, and treated with erythrocyte lysis buffer (eBioscience) to obtain a single cell suspension.

Standard cultures were established by seeding tumor cells in adherent culture flasks. Organoid cultures were established by seeding 1×10^5^ tumor cells in 50ul of Matrigel (Corning) and plated in 24-well plates. For the first 1-2 weeks of organoid initiation, Matrigel droplets were overlaid with recombinant organoid medium: Advanced DMEM/F-12 supplemented with 1X B27 (Gibco), 1X N2 (Gibco), 1.25mM nAcetylcysteine (Sigma), 10mM Nicotinamide (Sigma), 10nM Gastrin (Sigma), 100ng/ml EGF (Peprotech), 100ng/ml R-spondin1 (Peprotech), 100ng/ml Noggin (Peprotech), and 100ng/ml FGF10 (Peprotech). After organoids were established, cultures were switched to 50% L-WRN conditioned media(Miyoshi and Stappenbeck, 2013).

Organoid cultures were screened via immunohistochemistry and qPCR, and lines that uniformly expressed FoxA1, FoxA2, and NKX2-1 in all cells were selected for subsequent analysis. Standard culture cell lines were heterogeneous for NKX2-1, with both NKX2-1-positive and NKX2-1-negative cells identified within the same parental culture. In order to generate uniform cultures for downstream analysis our 2D cell lines were single cell cloned. The clonal populations were screened via immunoblotting for FoxA1, FoxA2, and NKX2-1, and the cell line with strongest expression of all three transcription factors was selected for subsequent analysis. The full name of the 2D cell line in this paper labeled 3311 is 3311-Tumor 3-Subclone 1, which was shortened for clarity.

### In vitro 4-hydroxytamoxifen treatment

Cells were transiently treated with 2μM 4-hydroxytamoxifen (Cayman Chemical Company, dissolved in 100% Ethanol) or vehicle for 48 (standard culture) or 72 (organoid culture) hours to activate CreER^T2^ nuclear activity and generate isogenic pairs.

### Immunoblotting

Cells were lysed on ice for 20 minutes in RIPA buffer (50mM Tris HCl pH 7.4, 150 mM NaCl, 0.1 % (w/v) SDS, 0.5% (w/v) sodium deoxycholate, 1% (v/v) Triton X-100) plus complete protease phosphatase inhibitor cocktail (A32961, Thermo Fisher Scientific). Cellular debris was pelleted for 15 minutes at 4 °C and protein concentration was quantitated with the Coomassie (Bradford) Protein Assay (ThermoFisher Scientific). A total of 15μg (organoids) or 30μg (cell lines) of protein lysates were separated on Tris-Glycine (TGX) precast gels (BIO RAD) and transferred to nitrocellulose membranes (Thermo Fisher Scientific). Membranes were probed overnight with antibodies to FoxA1 (1:1000, Abcam 23738), FoxA2 (1:1000, Abcam 108422), NKX2-1 (1:2000, Abcam 133638), CC3 (CST 9664S 1:1000), and Vinculin (1:20000, Abcam 129002). The next day membranes were probed with IRDye 800CW Goat anti-Rabbit IgG Secondary Antibody (1:20000, LI-COR) and imaged with a LI-COR Odyssey CLx and Image Studio Software.

### Generating a single cell suspension from organoid cultures

Matrigel droplets containing organoid cells were broken down via repeated pipetting in Cell Recovery Solution (Corning, 500μl per Matrigel droplet). Cell Recovery Solution containing organoids was transferred to sterile conical tubes and submerged in ice for 20-30 minutes before centrifugation at 4°C (300-500G). Cell Recovery Solution supernatant was removed and the cell pellet was washed via resuspension in PBS followed by centrifugation. Cells were then resuspended in pre-warmed TrypLE Express Enzyme (Thermo Fisher Scientific) and incubated for 5-7 minutes at 37°C. TrypLE reaction was quenched via dilution with cold Splitting Media (Advanced DMEM/F-12 [Gibco], 10 mM HEPES [Invitrogen], 1X Penicillin-Streptomycin-Glutamine [Invitrogen]). Cells were centrifuged and then resuspended in a pre-warmed DNase solution (L-WRN media supplemented to a final concentration of 200U/ml DNase [Worthington], 2.5 mM MgCl_2_, 500 μM CaCl_2_) and incubated for 5-7 minutes at 37°C. Cells were centrifuged and washed in PBS before use.

### Cell viability and growth assays

#### Presto Blue Assay

Organoids were broken down into a single cell suspension and seeded at equal density with 5ul of Matrigel per well in a solid wall, clear bottom 96 well plate with 100ul of LWRN per well. One day after seeding, a baseline measurement was taken before beginning a 48 hour treatment with 4-OHT (or Ethanol). 10ul of PrestoBlue™ HS Cell Viability Reagent was added to each well and incubated at 37C for 1 hour. After incubation, fluorescent emission was quantified on a Synergy HTX plate reader (Excitation: 528/20, Emission: 590/20, Read: 560 emission). Presto blue reagent was removed and wells were washed with warm PBS before adding fresh LWRN media. Measurements were taken every other day until organoids reached confluency.

#### Incucyte Live-Cell Imaging

Standard culture cells were seeded at a density of 7,000 cells per well (3311), 10,000 cells per well (H1651, H2009 and A549), or 20,000 cells per well (H358, H441, and H2122) in a 96-well plate and grown within an IncuCyte Live Cell Imaging System. Wells were imaged every two hours. Percent phase confluence was determined using IncuCyte Zoom software. Culture media was changed as needed and the assay was ended once one of the conditions reached 100% confluency.

#### CellTiter-Glo Assay

Organoid cells were broken down into a single cell suspension. Four identical 48-well plates were seeded with three wells per condition (5000 cells/well in 15 ul of Matrigel). The following protocol was performed on the day of seeding, and every other day for 6 days following seeding. CellTiter-Glo 3D reagent (Promega) and 50% L-WRN media were warmed to room temperature. A working solution was prepared with a ratio of 1 part CTG:5 parts L-WRN. Media was removed from each well, 330 μl of CTG working solution was added to each well containing organoids and one empty well before shielding from light and incubating at room temperature for 30 minutes on a plate shaker at 500 RPM. After incubation CTG working solution was transferred to a clear-bottom, solid-wall 96 well plate (100ul/well; 3 wells/replicate; 3 replicates/sample). Luminescence was measured using an EnVision 2105 Multimode Plate Reader (Perkin Elmer). Luminescent signal for each condition was normalized to the corresponding Day 0 luminescence read.

### Subcutaneous allografts

For subcutaneous allograft experiments, a single cell suspension of 3×10^5^ standard culture cells were mixed in a 1:1 volume with 50 μl of Matrigel. Cells were subcutaneously injected into the flank of NOD/SCID-gamma chain deficient mice (NSG). Tumor dimensions were measured with calipers, and tumor volume was calculated using the (L x W^2^)/2 formula. When the average volume of the tumors surpassed 200mm^3^, mice were randomized into Corn Oil or Tamoxifen cohorts (120mg/kg Tamoxifen). Mice received daily injections for 4 days. Tumor volume was monitored and measured every 3 days, and mice were euthanized once one tumor within the cohort surpassed 1000mm^3^ (Figure 1) or individually as their tumor volume surpassed 1000mm^3^ (Figure 4/4S).

### Lentiviral production and transduction

#### Cloning of EFS-FlpO lentivector

We generated a pCDH-EFS-FlpO lentiviral vector by PCR amplifying the EFS (EF1a) promoter from the pCDH-Cre plasmid(Han et al., 2014), digesting the purified PCR product with XbaI, and cloning into SnaBI-XbaI sites of pCDH-CMV-FlpO(Camolotto *et al*., 2018) via ligation with T4 DNA ligase. Correct identity and orientation of the construct was confirmed via Sanger sequencing

#### In vivo use

HEK293T cells were transfected with EFS-FlpO-encoding lentiviral vector, Δ8.9 packaging vector, and VSV-G envelope vector mixed with TransIT-293 (Mirus). Virus-containing supernatant was collected 36, 48, 60, and 72 hours after transfection, and then ultracentrifuged at 25,000 RPM for 2 hours to concentrate virus for in vivo infection. The concentration of the final viral stock was determined using the mouse fibroblast FlpO-GFP reporter cell line 3TZ to determine Plaque Forming Units/μl (PFU).

#### Cloning of human dual sgRNA constructs

Dual targeting vectors were generated by inserting a sgRNA1-scaffold-bovineU6-sgRNA2 cassette (synthesized as gene-blocks by IDT) into the LRG2.1-GFP-P2A-BlastR vector using Gibson assembly. The scaffold sequence for sgRNA was generated by altering the stem-stem loop region of the LRG2.1 scaffold(Shi et al., 2015) based on previously described CRISPRi sgRNA scaffolds(Adamson et al., 2016).

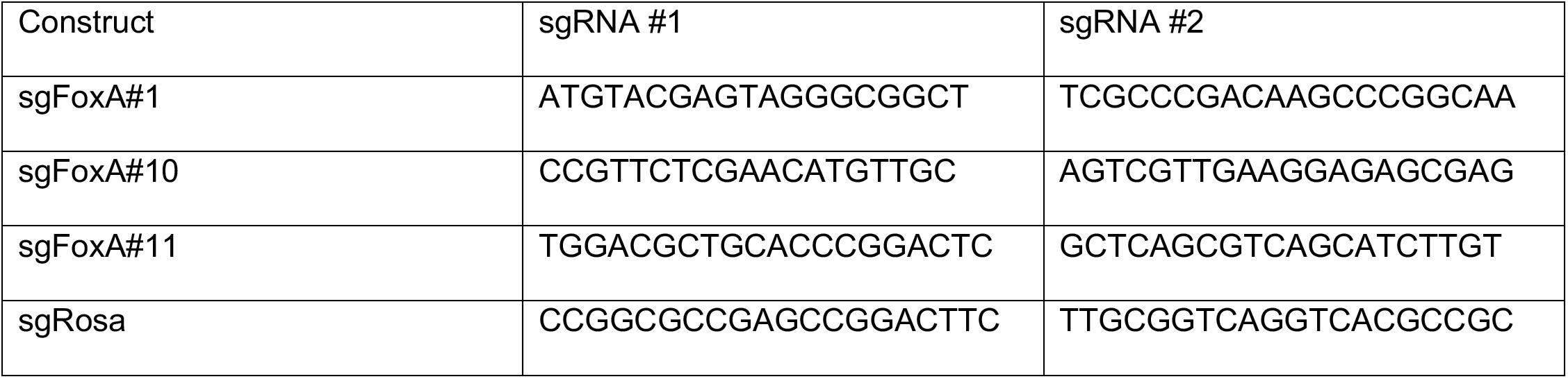

#### In vitro use

HEK293T cells were transfected with lentiCRISPRv2 (Addgene Plasmid #98290), the above dual sgRNA constructs, pLKO.shNkx2-1 (Addgene Plasmid #32400) or pLKO.shScramble (Addgene Plasmid #1864) lentiviral vectors, Δ8.9 packaging vector, and VSV-G envelope vector mixed with TransIT-293 (Mirus). Virus-containing supernatant was collected 48, 60, and 72 hours after transfection, centrifuged to pellet floating HEK293T cells, and filtered using 0.45μm filters before storing long term at -80°C.

#### Human CRISPR/Cas9

Human cell lines were transduced with lentiCRISPRv2 by culturing with undiluted lentiviral media containing 8 μg/ml polybrene for 48 hours total, refreshing the media and polybrene at 24 hours. Three days after transduction ceased, cells were subjected to Puromycin selection to produce stable cell lines expressing Cas9. Cells were then transduced with lentiviral sgNT, sgFoxA#1, sgFoxA#10, or sgFoxA#11 and selected with Blasticidin in the same manner. After three days of selection, cells were seeded for an Incucyte proliferation assay, protein was collected for immunoblotting, and RNA was collected for qRT-PCR analysis.

#### shRNA

Standard culture cell lines were transduced by culturing with undiluted lentiviral media containing 8 μg/ml polybrene for 48 hours total, refreshing the media and polybrene at 24 hours. Three days after transduction ceased, cells were subjected to Puromycin selection to produce stable lines. For stable transduction of isogenic organoid pairs post-4OHT/Ethanol treatment, Matrigel domes containing organoids were resuspended in ice cold Splitting Media and centrifuged at 300G 4°C twice. Cells were then resuspended in TrypLE Express Enzyme (Thermo Fisher Scientific), incubated for 5 minutes at 37°C, and quenched with ice cold Splitting Media. Cell pellets were resuspended in a 1:1 mixture, by volume, of 50% L-WRN and lentiviral media with a final concentration of 8 μg/ml polybrene, 1X Y-27632 (ROCK inhibitor), 1X A83-01 (ALK inhibitor), and 1X SB431542 (ALK inhibitor). The cell suspension was then transferred to a 6-well plate and centrifuged at 1700 RPM for 1 hour at room temperature. After centrifugation, the plate was incubated at 37°C for 2 hours. Cells were collected (scraping the bottom of the plate when necessary), centrifuged, resuspended in Matrigel, seeded in 24-well plates, and cultured in 50% L-WRN supplemented with 1X Y-27632, 1X A83-01, and 1X SB431542 for 72 hours. Three days after transduction, inhibitor media was washed away and cells were subjected to Puromycin selection to produce stable lines.

### RNA Sequencing

RNA was collected from biological replicates of isogenic organoid cultures 1027B, 1027D, and 1292B 3 weeks after 4-OHT/Ethanol treatment. RNA was isolated via Trizol-chloroform extraction followed by column-based purification. The aqueous phase was brought to a final concentration of 50% ethanol, and RNA was purified using the PureLink RNA Mini kit according to the manufacturer’s instructions (ThermoFisher Scientific). Library preparation was performed using the Illumina TruSeq Stranded mRNA Library Prep with UDI (Illumina; poly(A) selection). Sequencing was performed using the NovaSeq 6000 (50 x 50 bp paired-end sequencing; 25 million reads per sample).

### RNAseq Data Processing and Analysis

The mouse GRCm38 genome and gene feature files were downloaded from Ensembl release 102 and a reference database was created using STAR version 2.7.6a(Dobin et al., 2013). Optical duplicates were removed from NovaSeq runs via Clumpify v38.34(Bushnell, 2021). Reads were trimmed of adapters and aligned to the reference database using STAR in two pass mode to output a BAM file sorted by coordinates. Mapped reads were assigned to annotated genes using featureCounts version 1.6.3(Liao et al., 2019). Raw counts were filtered to remove features with zero counts and features with five or fewer reads in every sample. Differentially expressed genes were identified using a 5% false discovery rate with DESeq2 version 1.34.0(Love et al., 2014). For each genotype, we ran a separate model using Condition + Line in the design formula to compare 4-OHT vs EtOH while controlling for cell line effects on gene expression.

GSEA-Preranked was run with the differential gene list generated from DESeq2 and the following MSigDB gene sets: c2, c3, c4, c5, c6, c8, and Hallmarks on May 4, 2022. Gene sets smaller than 15 and larger than 500 were excluded from analysis. Marjanovic et al Clusters and Program gene lists and ADI/PATS signatures were downloaded from their supplemental materials.

#### Evaluation of NKX2-1 dependence

We first performed differential gene expression analysis (DESeq2) between the following pairs of samples: (1) shScramble vehicle treated (FoxA1/2-positive; NKX2-1-positive) vs shScramble tamoxifen treated (FoxA1/2-negative; NKX2-1-positive) (2) shScramble vehicle treated (FoxA1/2-positive; NKX2-1-positive) vs shNKX2-1 tamoxifen treated (FoxA1/2-negative; NKX2-1-knock down). We then intersected the differential gene lists for these two comparisons. Gene expression changes found in both comparisons were deemed NKX2-1 independent because the expression change is induced upon *Foxa1/2* deletion regardless of NKX2-1 levels. Gene expression changes unique to comparison (1) were deemed NKX2-1 dependent because the expression change induced upon *Foxa1/2* deletion only occurs when NKX2-1 is fully expressed.

### Single Cell RNA Sequencing

Mice were injected with tamoxifen 10 weeks after intubation and single cells were collected 12 weeks after intubation as follows. Lungs and heart were perfused with PBS. Individual macroscopic tumors were removed from the lungs and broken down into single cell suspensions as described in *Establishing primary murine LUAD cell lines and organoids* methods. Pre-depletion cells were viably cryopreserved in 5% DMSO/FBS. The remaining cells were depleted of CD45-positive and CD31-positive cells using MACS with Miltenyi microbeads (CD45: 130-052-301; CD31: 130-097-418) and LD columns (130-042-901) following manufacturer recommendations. Post-depletion cells were viably cryopreserved in 5% DMSO/FBS.

Protocols used to generate scRNA-seq data with 10x Genomics Chromium platform can be found at https://support.10xgenomics.com/single-cell-gene-expression.

In brief, the Chromium Single Cell Gene Expression Solution with 3’ chemistry, version 3 (PN-1000075) was used to barcode individual cells with 16bp 10X barcodes and to tag cell specific transcript molecules with 10bp Unique Molecular Identifier (UMI) according to the manufacturer’s instructions. The following protocol was performed at the High-Throughput Genomics Shared Resource at Huntsman Cancer Institute, University of Utah. Single cells were suspended in phosphate buffered saline with 0.04% bovine serum albumin, and the cell suspension was passed through a 40 micron cell strainer. Viability and cell count were assessed on Countess II (Thermo Scientific). Suspensions were equilibrated to targeted cell recovery of 8000 cells. For the KP sample without stromal depletion, the targeted cell recovery was 7500 cells. 10x Gel Beads and reverse transcription reagents were added and cell suspensions were loaded to Chromium Single Cell A (PN-120236) to form Gel Beads-in emulsions (GEMs) - the nano-droplets. Within individual GEMs, cDNA generated from captured and barcoded mRNA was synthesized by reverse transcription at the setting of 53°C for 45 min followed by 85°C for 5 min. Subsequent A tailing, end repair, adaptor ligation and sample indexing were performed in bulk according to the manufacturer’s instructions. The resulting barcoding libraries were qualified on Agilent D1000 ScreenTape on Agilent Technology 2200 TapeStation system and quantified by quantification PCR using KAPA Biosystems Library Quantification Kit for Illumine Platforms (KK4842). Multiple libraries were then normalized and sequenced on NovaSeq 6000 with 2 × 150 PE mode.

### scRNAseq Data Processing and Analysis

#### Demultiplexing and data alignment

Single-cell RNA-seq data from both KP (n=2) and KPF_1_F_2_ (n=2) tumors were demultiplexed using the 10x cellranger mkfastq version 3.1.0 to create fastq files with the I1 sample index, R1 cell barcode+UMI, and R2 sequence. Reads were aligned to the mouse genome (mm10 with custom CRE-ERT2 and FoxA1/2 individual exon references) and UMIs were generated using cellranger count 3.1.0 with expected-cells set to 8000 per library. For the KP sample without stromal depletion, expected-cells was set to 7500. QC reporting, clustering, and dimension reduction were performed for initial data evaluation in 10x Genomics’ Cell Loupe Browser (v5.0). For the KP sample without stromal depletion, we captured 5,963 cells total with 48,090 mean reads per cell and 1,488 median genes per cell. For the KP sample with stromal depletion, we captured 8,536 cells total with 25,593 mean reads per cell and 736 median genes per cell. For the KPF_1_F_2_ sample without stromal depletion, we captured 4,662 cells total with 55,589 mean reads per cell and 1,552 median genes per cell. For the KPF_1_F_2_ sample with stromal depletion, we captured 4,415 cells total with 60,226 mean reads per cell and 1,339 median genes per cell. Additional details of the primary Cell Ranger data processing can be found at: https://support.10xgenomics.com/single-cell-gene-expression/software/pipelines/latest/algorithms/overview.

#### Quality control, clustering, and cell type identification

Single cell expression data was subjected to common Seurat workflows for initial quality control and clustering (https://satijalab.org/seurat/articles/pbmc3k_tutorial.html). Cells with unique feature counts over 7500 or less than 200 and over 20% mitochondrial counts were filtered out for downstream analysis. Counts of cells passing QC were then log normalized and scaled based on all genes using Seurat’s NormalizeData and ScaleData functions. PCA linear dimension reduction was performed and Seurat’s FindNeighbors function was employed to embed single cell profiles in a K-nearest neighbor (KNN) graph based on a (30 PC) PCA space. The FindClusters function was utilized to iteratively group cells together using Louvian algorithm modularity optimization techniques. Clustering was performed based on the top 30 dimensions using Seurat’s RunUMAP function. Differentially expressed genes for each cluster were identified using Seurat’s FindMarkers function using default setting. The Mouse Cell Atlas’ scMCA R package was run with the top differentially expressed genes for each cluster (Supplemental Table 2) to facilitate cell type identification. Following QC filtering and tumor cell identification, 4646 KP and 2481 KPF1F2 high quality tumor cells remained. For subsequent analyses, identified tumor cells from KP and KPF1F2 tumors were subsetted out and reclustered based on the top 25 dimensions. Cell barcodes identified as tumor cells that were used for downstream analyses are included in Supplemental Table 3. Reclustering of identified tumor cells in UMAP space revealed 13 clusters (Figure 3A) To identify KPF_1_F_2_ complete recombinants, expression of the floxed second exon of Foxa1 was visualized (Figure 3C).

#### Differential gene expression and signature score assignment

Differentially expressed genes in each of the tumor cell clusters were calculated using Seurat’s FindMarkers function. Differentially expressed genes of UMAP clusters from all KP and KPF_1_F_2_ tumor cells can be found in Supplemental Table 3. Gene module scores for several published gene signatures for AT1 cells AT2 cells(Du *et al*., 2015; Little *et al*., 2019; Marjanovic *et al*., 2020; Travaglini *et al*., 2020), and various additional cell types and states including stratified squamous epithelium (MSigDB) were determined per cell across the tumor cell and complete recombinant data sets using Seurat’s AddModuleScore function. Gene lists for each of these scores can be found in Supplemental Table 5.

### Chromatin Immunoprecipitation Sequencing

Organoids were broken down into a single cell suspension before following the manufacturer protocol v1.5 for Chromatrap ChIP-seq Protein A kits (500189) with the following modifications/optimizations:

<u>Step 1 (Chromatin preparation, fixation, collection):</u> Broken down, individual organoid cells were treated as suspension cells for crosslinking.
<u>Step 2 (Cell lysis and chromatin shearing):</u> Crosslinked cells were suspended in 100ul of Lysis Buffer/2,000,000 cells. Sonication was performed with 20 cycles of 30 seconds on:30 seconds off at 70% to reach fragmentation of 100-500bp lengths. Input DNA was purified with Zymo ChIP DNA Clean and Concentrator columns (D5205). DNA concentration was determined using Qubit dsDNA High Sensitivity Assay Kit (Q32851).
<u>Step 3 (Slurry preparation and immunoprecipitation):</u> NKX2-1 IP was performed with a sample:antibody ratio of 10ug sample:4.2ug antibody (Abcam ab133737). FoxA2 IP was performed with a sample:antibody ratio of 10ug sample: 5ug antibody (CST D56D6 custom high concentration formulation)
<u>Step 4 (Reverse cross-linking):</u> IP DNA was purified with Zymo ChIP DNA Clean and Concentrator columns (D5205).

IP and input samples were prepared for sequencing with NEBNext Ultra II DNA Library Prep Kit and sequenced on an Illumina Novaseq 6000 as paired-end 150 bp reads (HCI High-Throughput Genomics core).

### ChIPseq Data Processing and Analysis

Samples were de-multiplexed with Illumina BCL2Fastq using dual indexes and single bp mismatch. Fastq reads were aligned to the standard chromosomes of mouse genome, version mm10, using Novocraft novoalign (v4.03.01 http://www.novocraft.com) with provided tuned settings for Novaseq. Sequence adapter sequences were provided to Novoalign for masking during alignment. Alignment pairings were checked with samtools (v1.10) fixmate function.

Samples were run through the MultiRepMacsChIPSeq pipeline (v17 https://github.com/HuntsmanCancerInstitute/MultiRepMacsChIPSeq) with five independent samples. After comparison, four replicates were chosen for analysis, based on correlation metrics and number of peaks identified. Briefly, the pipeline consists of the following steps. Exclusion intervals representing high-copy genomic sequences were first generated by calling peaks on combined, unfiltered Input samples. Any alignments overlapping exclusion intervals were discarded from further analysis. Only properly-paired alignments were used; chimeric and singleton alignments were discarded. Duplicate alignments based on coordinates were randomly subsampled to a final rate of 5% across all samples from a mean of 24% (range 18-32%). Peaks were called individually for each replicate using MACS2 (v2.2.6 https://github.com/macs3-project/MACS) with peak calling parameters of qvalue 0.01, min-length 200 bp, max-gap 100 bp, and genome size (empirically determined) 2554209000 bp. Called peaks were first combined between replicates for each condition, EtOH and Tamoxifen treated, and then combined into a master peak list for both conditions using BEDtools (v2.28.0 https://bedtools.readthedocs.io). Mean fragment coverage tracks were generated using bam2wig (BioToolBox, v1.68 https://github.com/tjparnell/biotoolbox) by first depth-normalizing to 1 million paired-end fragments, then averaging between replicates. Multi-mapping alignments were scaled by the number of genomic hits reported by the aligner. Mean Log2 Fold Enrichment tracks were generated by using MACS2 bdgcmp function with depth-normalized ChIP coverage and generated mean lambda control coverage tracks, then converting to base 2 log values with manipulate_wig (BioToolBox). Count tracks were generated with bam2wig scaling to a total depth of 20M fragments, the lowest observed depth amongst all samples.

Peaks were scored using get_datasets (BioToolBox) with the depth-normalized count files generated by the MultiRepMacsChIPSeq pipeline. Fractional fragment counts were rounded to the nearest integer for simplicity. Differential peaks were identified by running MAnorm2(Tu *et al*., 2021) with recommended parameters, using “parametric” for fitting the mean variance curve and initial coefficients of 0.1 and 10. Differential peaks were identified by a padj value < 0.05. Data was collected for peaks by using BED files of the peaks of interest using the BioToolBox (v1.68) programs get_relative_data, using 50 bins of 40 bp surrounding the peak midpoint (±1 kb). Heat maps and plots were generated with custom R scripts using pHeatmap (https://cran.r-project.org/package=pheatmap) and ggplot2 (https://ggplot2.tidyverse.org). Peaks were annotated with ChIPseeker(Yu et al., 2015) and Ensembl gene annotation (release 98), restricted to expressed, protein-coding, Gencode transcripts. Motifs, both known and novel, were identified using the Homer package (http://homer.ucsd.edu/homer/).

## QUANTIFICATION AND STATISTICAL ANALYSIS

All graphing and statistical analysis was performed with PRISM software, with all graphs showing mean and standard deviation. The statistical details can be found in the corresponding figure legend. All NGS statistical analysis was performed according to published pipeline protocols cited, with a statistical significance cutoff of padj<0.05.

---

**Supplemental Figure 1, related to Figure 1.**
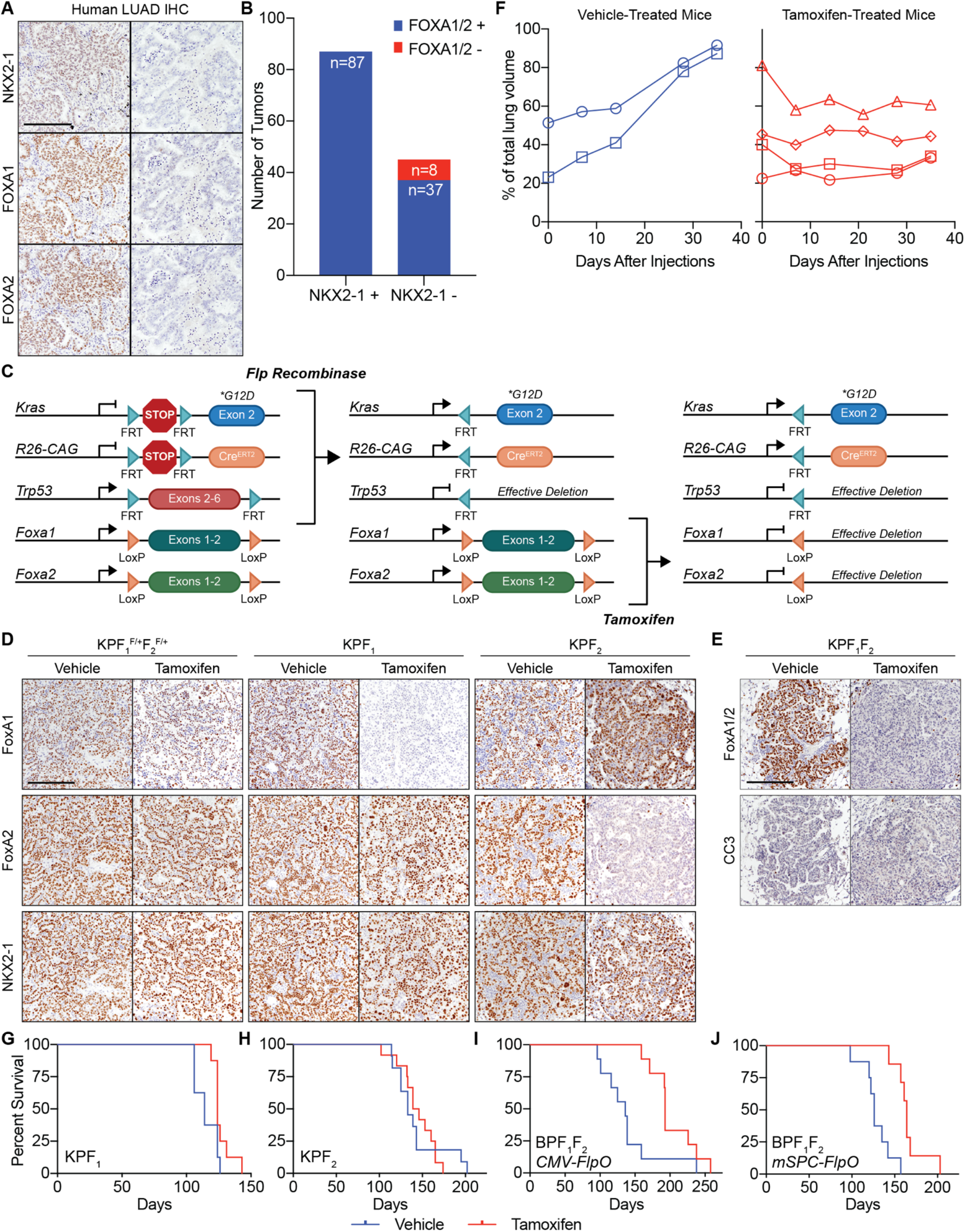
A) Representative IHC images from human tissue microarrays illustrating that NKX2-1-positive tumors retain FoxA1/2 expression while NKX2-1-negative tumors can lose FoxA1/2 expression (scale bar 500 μm). B) Histological analysis of human tumors. All NKX2-1 positive LUAD tumors evaluated (n=87) express FOXA1 and/or FOXA2. FOXA1 and/or FOXA2 were detected in 82% (37/45) of NKX2-1-negative LUAD tumors evaluated, while there was no detectable FOXA1 or FOXA2 in 18% (8/45) of NKX2-1-negative tumors. (p<0.0001, Fisher’s Exact test). C) KPF_1_F_2_ mouse model. Mice are intubated with viral FlpO recombinase, which activates the expression of oncogenic KRAS^G12D^ and Cre^ERT2^ and deletes *Trp53* in normal lung epithelial cells. This initiates the development of tumors in which Cre^ERT2^ is expressed, but sequestered in the cytoplasm. After tumors have developed, tamoxifen injection allows Cre^ERT2^ to enter the nucleus exclusively in tumor cells and delete *Foxa1* and *Foxa2*. D) Representative IHC images of KPF_1_^F/+^F_2_^F/+^ (KP), KPF_1_, and KPF_2_ mice from the experiment shown in Figure 1B-C illustrating specific deletion of FoxA1 and FoxA2 (scale bar 500 μm). E) Representative IHC images of KPF_1_F_2_ vehicle and tamoxifen treated mice from the experiment shown in Figure 1B-C, illustrating a lack of CC3 staining at this timepoint in either condition. F) Autochthonous lung tumors were initiated in KPF_1_F_2_ mice and tracked via μCT scans monthly until there was significant detectable/traceable lung tumor volume in the three-dimensional reconstruction from each mouse (14-17 weeks post-initiation), at which point mice were treated with either vehicle (n=2, left panel) or tamoxifen (n=4, right panel). Mice were scanned weekly to analyze tumor progression as a measurement of percentage of total lung volume. G) KPF_1_ survival study in which tamoxifen (red, n=8) or vehicle (blue, n=8) was administered 10 weeks post-initiation with lentiviral EFS-FlpO (vehicle median survival of 16.29 weeks; tamoxifen median survival of 17.71 weeks; p=0.0290, Log-rank Mantel-Cox test). H) KPF_2_ survival study in which tamoxifen (red, n=12) or vehicle (blue, n=11) was administered 10 weeks post-initiation with lentiviral EFS-FlpO (vehicle median survival of 19.00 weeks; tamoxifen median survival of 20.36 weeks; p=0.9499, Log-rank Mantel-Cox test). I) BPF_1_F_2_ survival study in which tamoxifen (red, n=9) or vehicle (blue, n=9) was administered 6 weeks post-initiation with Adenoviral CMV-Flp (vehicle median survival of 19.43 weeks; tamoxifen median survival of 27.57 weeks; p=0.0098, Log-rank Mantel-Cox test). J) BPF_1_F_2_ survival study in which tamoxifen (red, n=7) or vehicle (blue, n=8) was administered 6 weeks post-initiation with Adenoviral SPC-Flp (vehicle median survival of 18.00 weeks; tamoxifen median survival of 23.43 weeks; p=0.0006, Log-rank Mantel-Cox test).

**Supplemental Figure 2, related to Figure 2.**
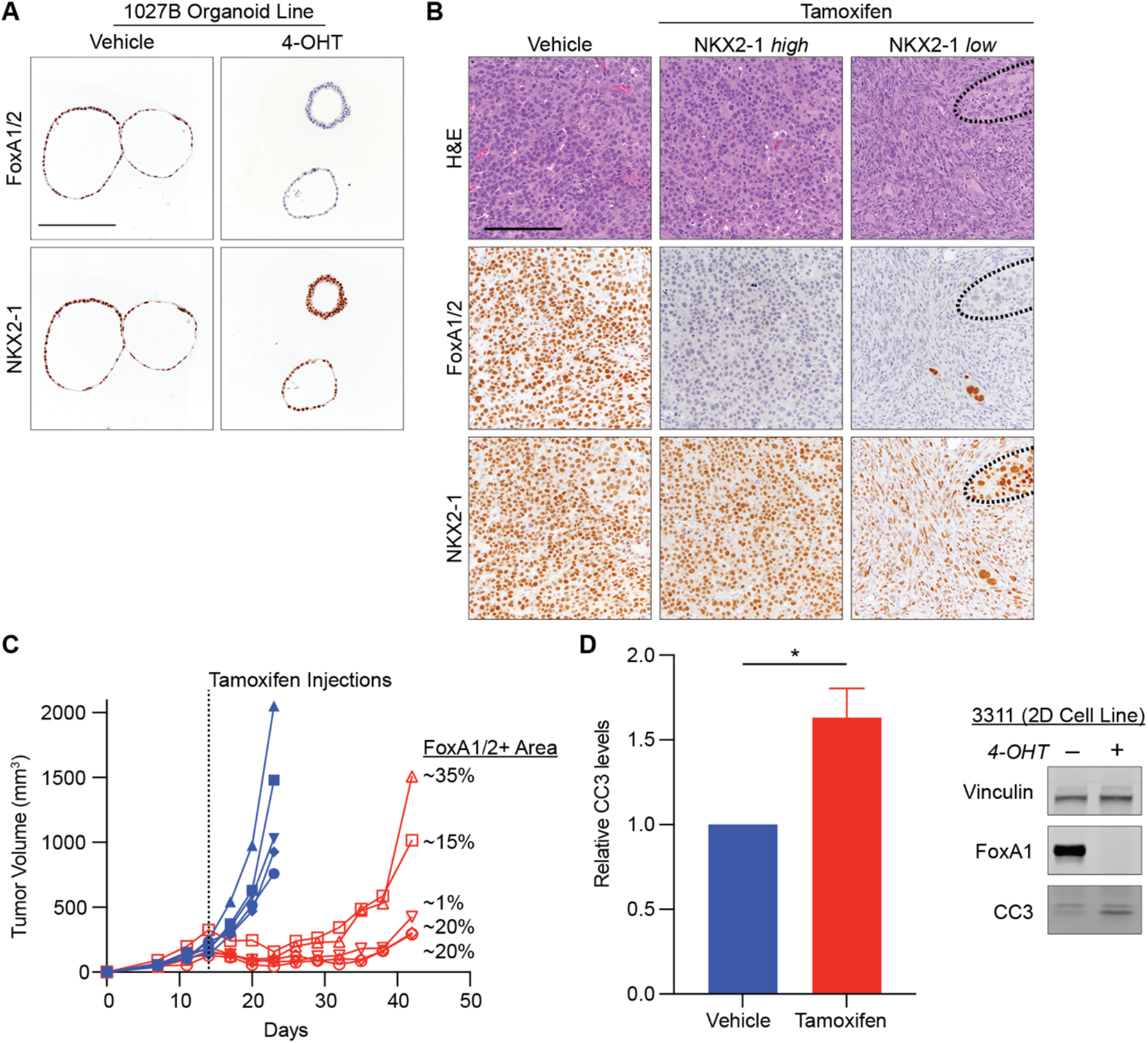
A) IHC for FoxA1/2 and NKX2-1 in KPF_1_F_2_ organoid line 1027B collected 3 days post 72-hour treatment with 4-OHT or vehicle demonstrating complete loss of FoxA1/2 (scale bar 500 μm). B) Representative images of subcutaneous tumors from experiment shown in E. H&E and IHC for FoxA1/2 and NKX2-1. Tamoxifen-treated tumors harbored two phenotypes: NKX2-1-high tumors with similar morphology to FoxA1/2-positive tumors and NKX2-1-low tumors with sarcomatoid/quasi-mesenchymal morphology (NKX2-1-high region outlined in dotted line; scale bar 500 μm). C) Individual tumor measurements for subcutaneous tumors in Figure 2D. Approximated percent of tumor for each tamoxifen-treated mouse that retained FoxA1/2 at time of tumor collection indicated on the right. D) Relative Cleaved Caspase 3 protein levels 3 days following initiation of 4-hydroxytamoxifen treatment. Western blots imaged with Licor system and quantified with ImageStudio (unpaired t-test *p=0.0359; one representative western blot shown of two biological replicates).

**Supplemental Figure 3, related to Figure 3.**
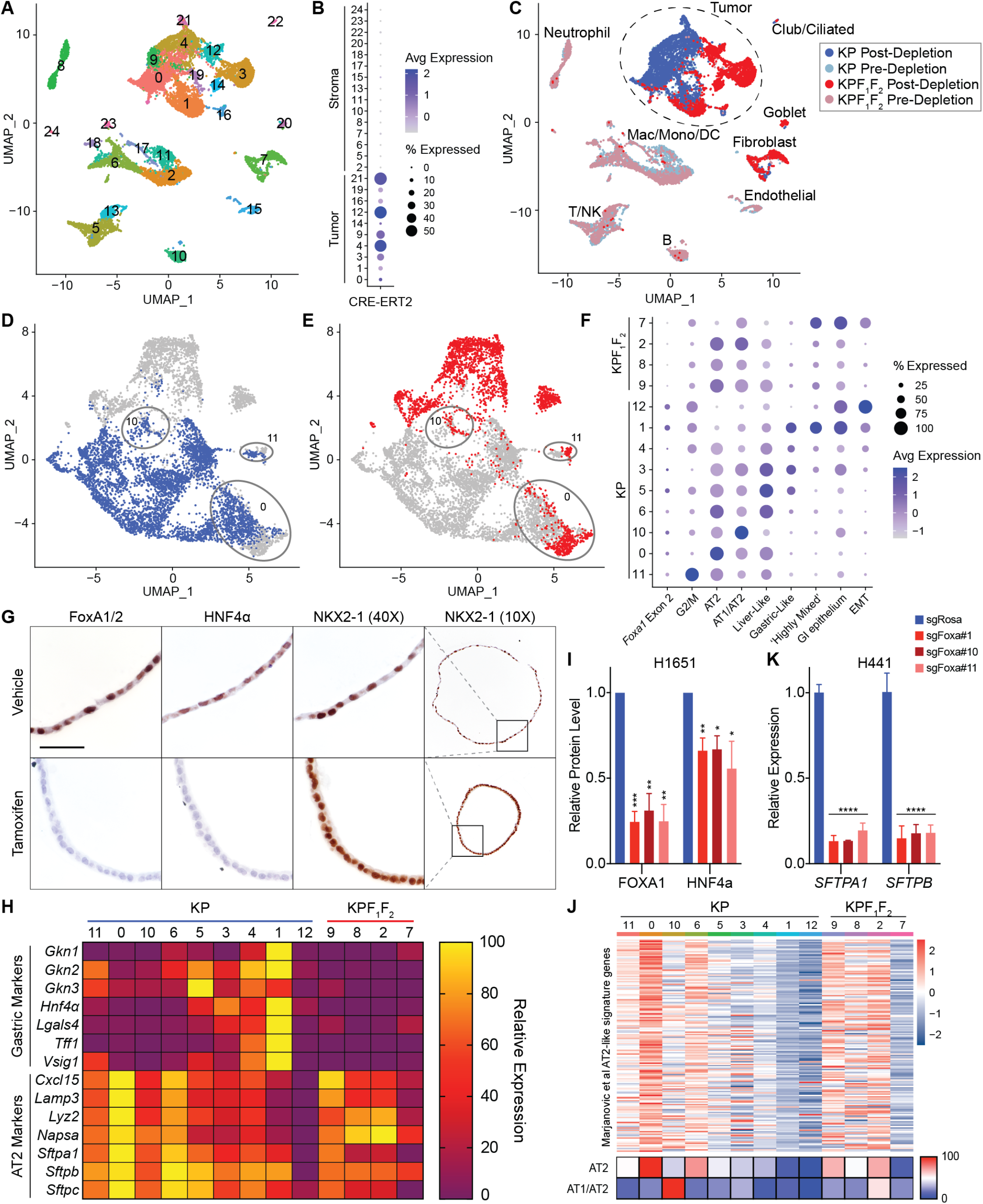
A) UMAP of QC-filtered cells from KP (8269 cells) and KPF_1_F_2_ (6482 cells) macro-dissected tumor regions, both pre- and post-MACS depletion of CD45+ and CD31+ cells. Cells colored by cluster identity. B) Dot plot quantifying the level of expression and proportion of cells expressing CRE-ERT2, a tumor-specific transcript, per cluster. CRE-ERT2 expression indicates that the cells within that cluster are tumor cells, while a complete lack of expression indicates a stromal identity. C) UMAP shown in A with cells colored by genotype and CD45/CD31 depletion status. Clusters included in the tumor cell reclustering (Figure 3A) circled with dashed line. D) UMAP of cells from KP (blue) and E) KPF_1_F_2_ samples (red) illustrating the level of overlap between the two genotypes in clusters 0, 10, and 11. KPF_1_F_2_ cells in these clusters retain expression of *Foxa1* Exon 2 indicating incomplete recombination. F) Dot plot quantifying *Foxa1* Exon2 expression and Marjanovic program module scores in the tumor cell UMAP of Figure 3A illustrating enrichment for *Foxa1* Exon2 and the mixed-lineage tumor progression within KP-specific clusters relative to KPF_1_F_2_ clusters. G) Representative IHC images of gastric transcription factor HNF4α and pulmonary transcription factor NKX2-1 in FoxA1/2-positive and FoxA1/2-negative 1027B organoids, illustrating the loss of gastric lineage upon *Foxa1/2* deletion (scale bar 100 μm). H) Row normalized average expression levels of gastric and AT2 genes in each cluster shown in Figure 3A demonstrating a distinct loss of gastric identity markers in FoxA1/2-negative clusters (2,7,8,9) relative to gastric-like FoxA1/2-positive clusters (1,3,4,5,6) and a decrease in AT2 identity markers in FoxA1/2-negative clusters relative to AT2-like FoxA1/2-positive Cluster 0. I) Relative protein levels of FOXA1 and HNF4a in H1651 following CRISPR-Cas9 deletion of *FOXA1/2*. Western blots imaged using Licor system and quantified using Image Studio (n=3 biological replicates; ***p<0.0001, **p<0.001, *p<0.009). J) Top: Heatmap representing average expression of the AT2-like signature in which each row is a gene of the AT2-like program and each column is a cluster’s average expression of an individual gene. Bottom: Heatmap of average AT2 and AT1/AT2 module scores in each tumor cell cluster K) Relative expression of *SFTPA1* and *SFTPB* in H441 cells lines (shown in Figure 2) following CRISPR-Cas9 deletion of *FOXA1/2* as determined by RT-qPCR (n=2 biological replicates with 3 technical replicates each; ****p<0.000001).

**Supplemental Figure 4, related to Figure 4.**
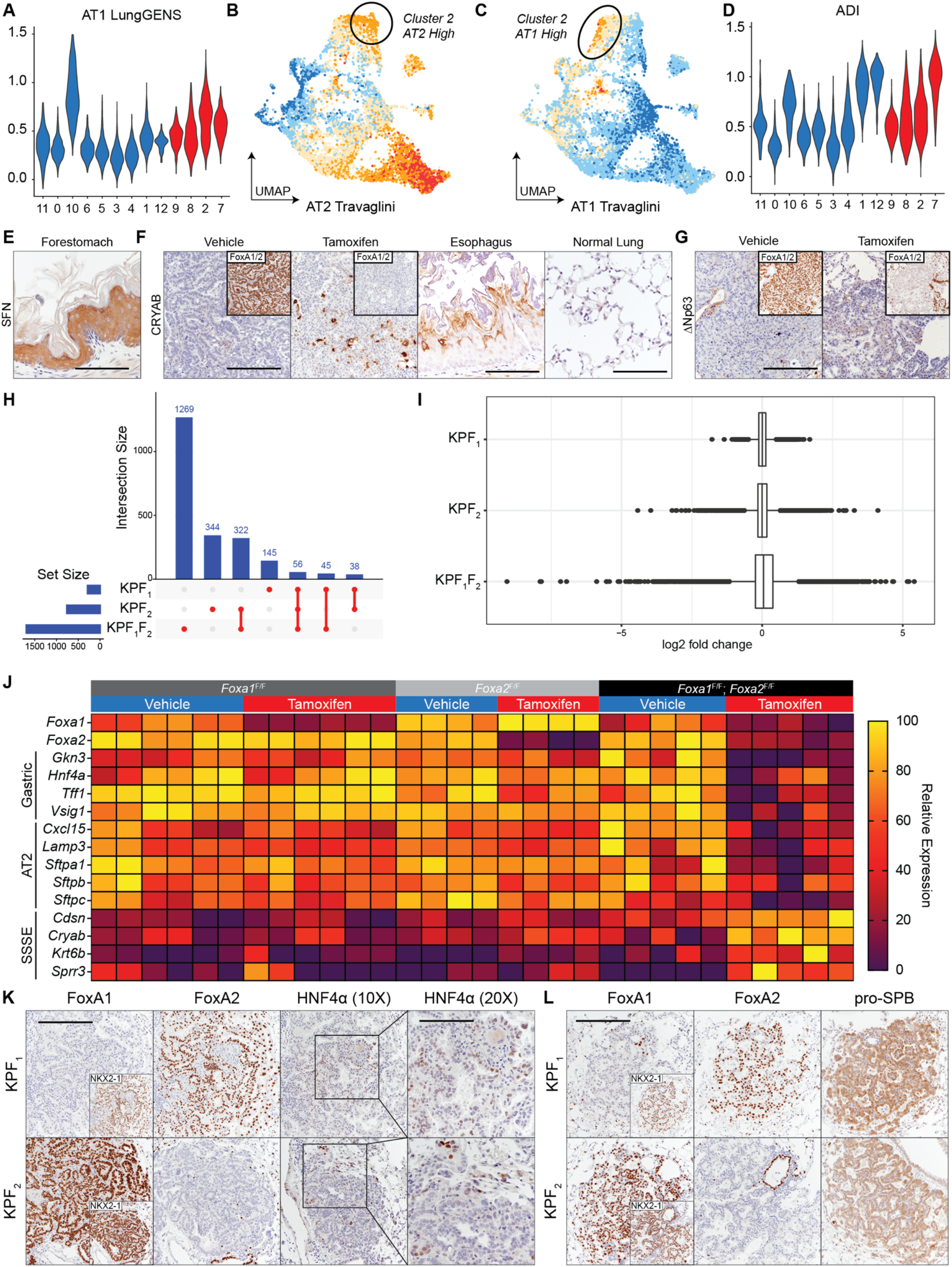
A) Violin plot representing the expression of the LungGENS AT1 transcriptional program in each tumor cell cluster. The y-axis indicates gene module score, and the x-axis indicates cluster identity. B) UMAP of gene module scores for AT2 and… C) AT1 Travaglini gene signatures, illustrating that there is an inverse correlation between AT2 and AT1 identity within the cells that make up KPF_1_F_2_-specific Cluster 2. D) Violin plot representing the expression of the ADI transcriptional program in each tumor cell cluster. E) Representative IHC image of SSSE/PATS marker Stratifin (SFN) in normal SSSE tissue found in the murine forestomach (Scale bar 250 μm). F) Representative IHC images of SSSE/AT1 marker Alpha Crystallin-B in normal SSSE (Esophagus) and FoxA1/2-positive, and FoxA1/2-negative tumors, confirming the induction of these identities upon *Foxa1/2* deletion. Of note, with this antibody we do not detect Alpha Crystallin-B in normal AT1 cells (tumor scale bar 500 μm, esophagus and lung scale bar 250 μm). G) Representative IHC images of ΔNp63 in FoxA1/2-positive and FoxA1/2-negative tumors, illustrating the lack of expression in both tumor types (scale bar 500 μm). H) Upset plot showing the intersection between DEGs in KPF_1_, KPF_2_, and KPF_1_F_2_ 4-OHT and vehicle-treated organoids. Set size bar chart indicating number of DEGs identified in each genotype. Intersection size bar chart indicates number of DEGs shared between gene lists indicated by red dots. I) Log2 fold change of each statistically significant gene identified in DEGs of KPF_1_, KPF_2_, and KPF_1_F_2_ 4-OHT and vehicle-treated organoids, illustrating a large increase in the magnitude of gene expression changes upon concomitant deletion of both *Foxa1* and *Foxa2* when compared to the range of log2 fold change values for deletion of either gene alone. J) Row normalized relative expression levels of *Foxa1*, *Foxa2*, and select gastric, AT2, and SSSE genes in each organoid RNAseq sample, illustrating that single deletion of *Foxa1* (left) and *Foxa2* (center) is not sufficient to alter expression of marker genes to the same extent that is observed upon deletion of both (right). K) Representative IHC images of gastric transcription factor HNF4α in tumors from tamoxifen-treated KPF_1_, KPF_2_ mice, illustrating retained expression following individual deletion (scale bar 500 μm; 20X scale bar 250 μm). L) Representative IHC images of AT2 marker pro-SPB in tumors from tamoxifen-treated KPF_1_, KPF_2_ mice, illustrating retained expression following individual deletion (scale bar 500 μm).

**Supplemental Figure 5, related to Figures 3 and 4.**
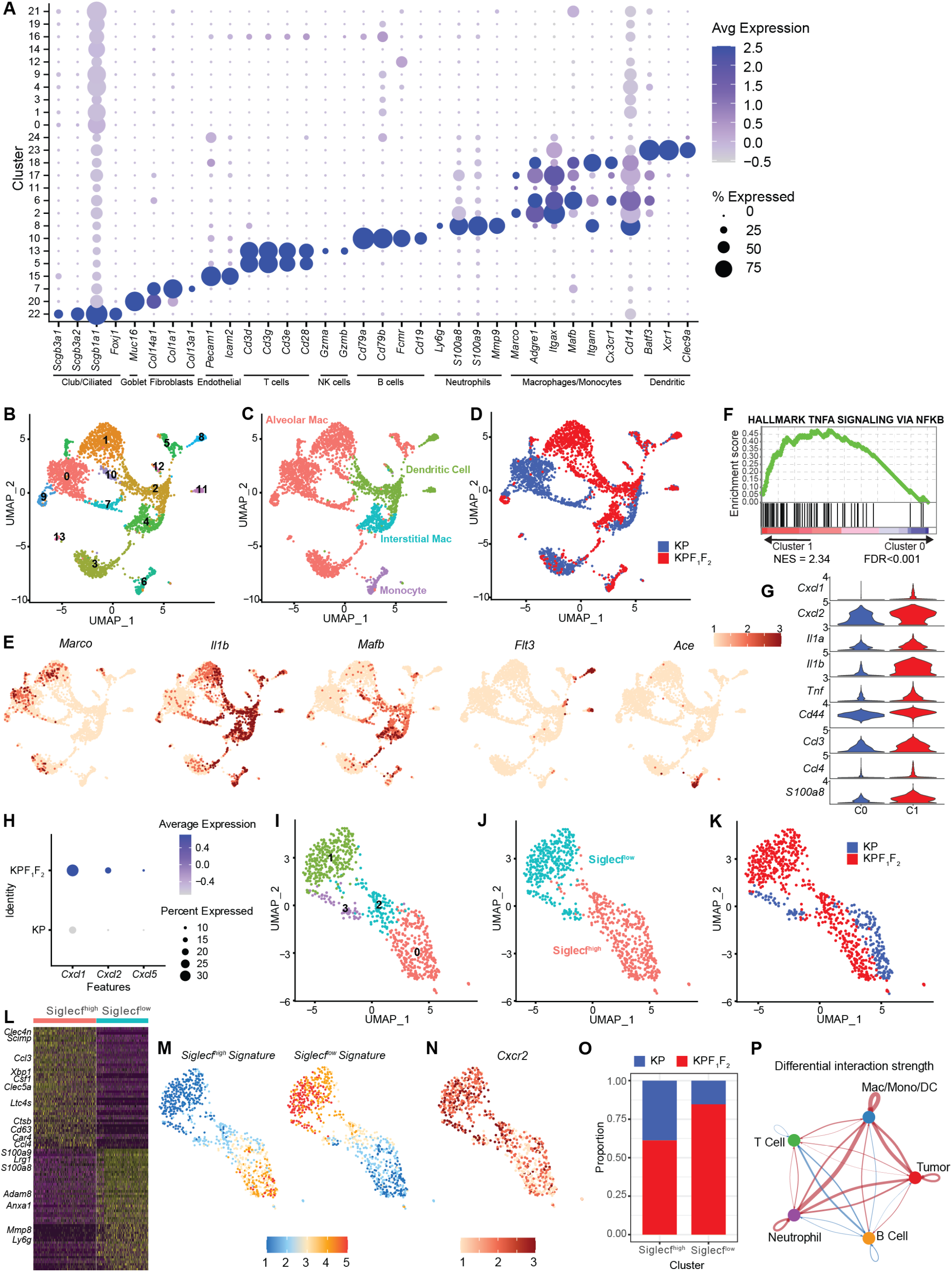
A) Dot plot quantifying expression levels of stromal marker genes in individual clusters (Supplemental Figure 3A). *Scgb3a1, Scgb3a2, Scgb1a1*, and *Foxj1* are markers of club/ciliated cells. *Muc16* is a marker of goblet cells. *Col14a1, Col1a1,* and *Col13a1* are markers of fibroblasts. *Pecam1* and *Icam2* are markers of endothelial cells. *Cd3d, Cd3g, Cd3e,* and *Cd28* are markers of T cells. *Gzma* and *Gzmb* are markers of NK cells. *Cd79a, Cd79b, Fcmr,* and *Cd19* are markers of B cells. *Ly6g, S100a8, S100a9,* and *Mmp9* are markers of neutrophils. *Marco, Adgre1, Itgax, Mafb, Itgam, Cx3cr1,* and *Cd14* are markers of macrophages/monocytes. *Batf3, Xcr1,* and *Clec9a* are markers of dendritic cells. B-C) UMAP of macrophages, monocytes, and dendritic cells from KP (1761 cells) or KPF1F2 (1530 cells) macro-dissected tumor regions. Cells colored by B) cluster identity, C) cell subtype identity, or D) genotype. E) UMAPs of relative expression of subtype marker genes (Alveolar macrophages: *Marco*, M1-like macrophages: *Il1b*, Interstitial macrophages: *Mafb,* DCs: *Flt3*, Monocytes: *Ace*). F) GSEA showing enrichment of the hallmark signature associated with pro-inflammatory TNFA signaling in KPF1F2-specific alveolar macrophage cluster 1 compared to KP-specific cluster 0. G) Violin plot showing relative expression of M1-like, pro-inflammatory genes enriched in KPF1F2-specific cluster 1 vs KP-specific cluster 0. H) Dot plot showing expression of neutrophil recruitment chemokines *Cxcl1, Cxcl2*, and *Cxcl5* in KPF1F2 vs KP tumor cells. I-K) UMAP of neutrophils from KP (215 cells) or KPF1F2 (549 cells) tumor regions. Cells colored by I) cluster identity, J) cell subtype identity, or K) genotype. L) Heatmap of the top 50 differentially expressed genes in Siglecf^high^ vs Siglecf^low^ neutrophils, with genes of interest labeled. M) UMAPs of gene module scores of Siglecf^high^ and Siglecf^low^ gene signatures from Engblom et al. N) UMAP of relative expression of *Cxcr2* in neutrophils. O) Proportion of Siglecf^high^ and Siglecf^low^ cells per genotype. P) CellChat analysis showing differential interactions between immune cell subtypes. Red lines indicate stronger interactions in KPF1F2 tumors compared to KP and blue lines indicate stronger interactions in KP tumors compared to KPF1F2. Edge weights are proportional to the magnitude of differential interaction strength between genotypes.

**Supplemental Figure 6, related to Figure 6.**
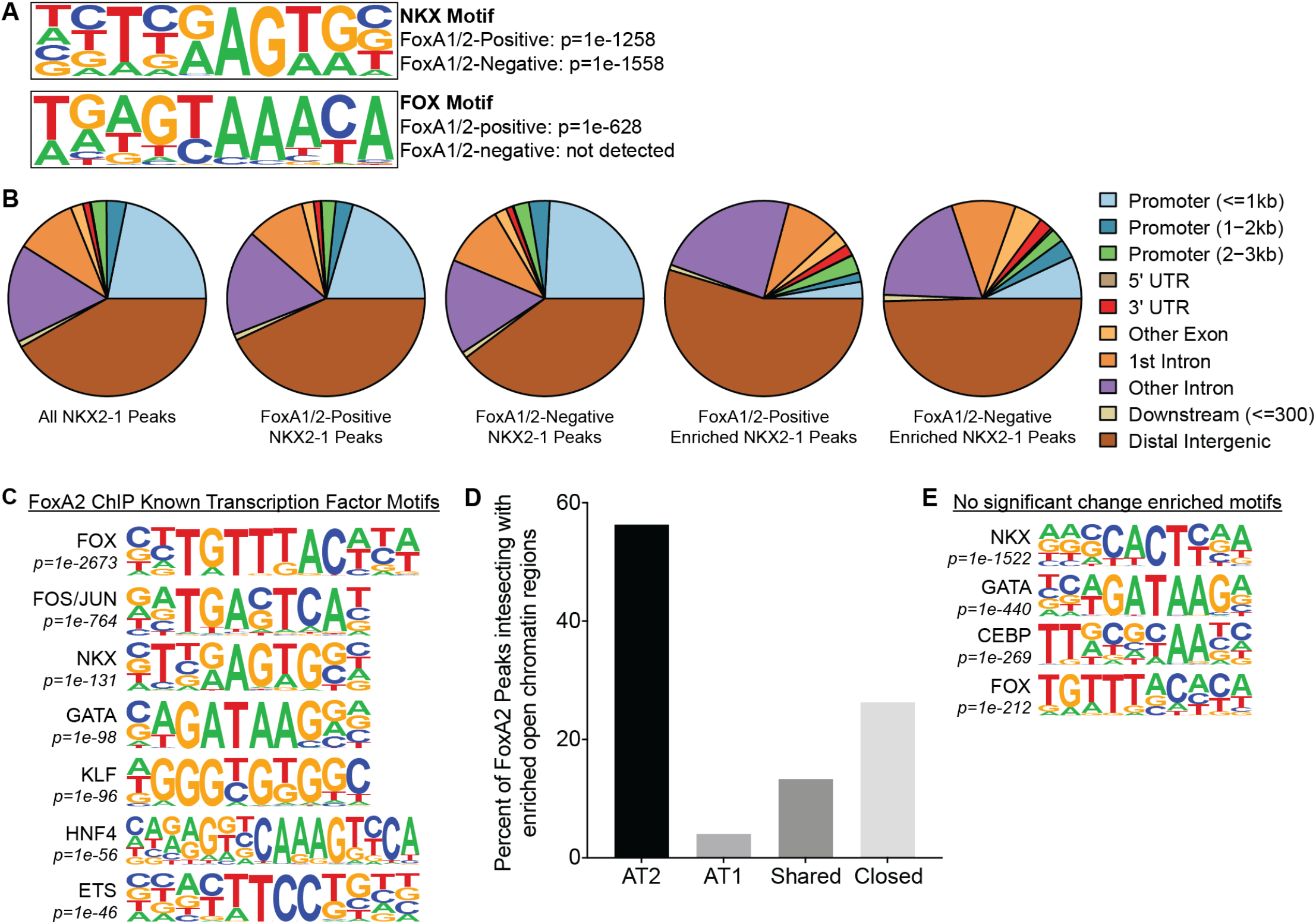
A) NKX and FOX transcription factor motifs and their corresponding p-values in FoxA1/2-positive and FoxA1/2-negative NKX2-1 ChIP peaks. B) Genomic occupancy of all NKX2-1 ChIP peaks, FoxA1/2-positive NKX2-1 ChIP peaks, FoxA1/2-negative NKX2-1 ChIP peaks, NKX2-1 ChIP peaks enriched in FoxA1/2-positive cells, and NKX2-1 ChIP peaks enriched in FoxA1/2-negative cells. C) Select transcription factor motifs detected in FoxA2 ChIPseq peaks. D) The percent of all FoxA2 ChIPseq peaks found within open chromatin regions specific to normal murine AT2 or AT1 cells, open chromatin regions shared in both AT2 and AT1 cells, or regions of chromatin closed in both AT2 and AT1 cells. The majority (56%) of FoxA2 peaks are found within open chromatin regions unique to AT2 cells. E) Select transcription factor motifs for NKX2-1 peaks not called as significantly enriched in either condition.

**Supplemental Figure 7, related to Figure 7.**
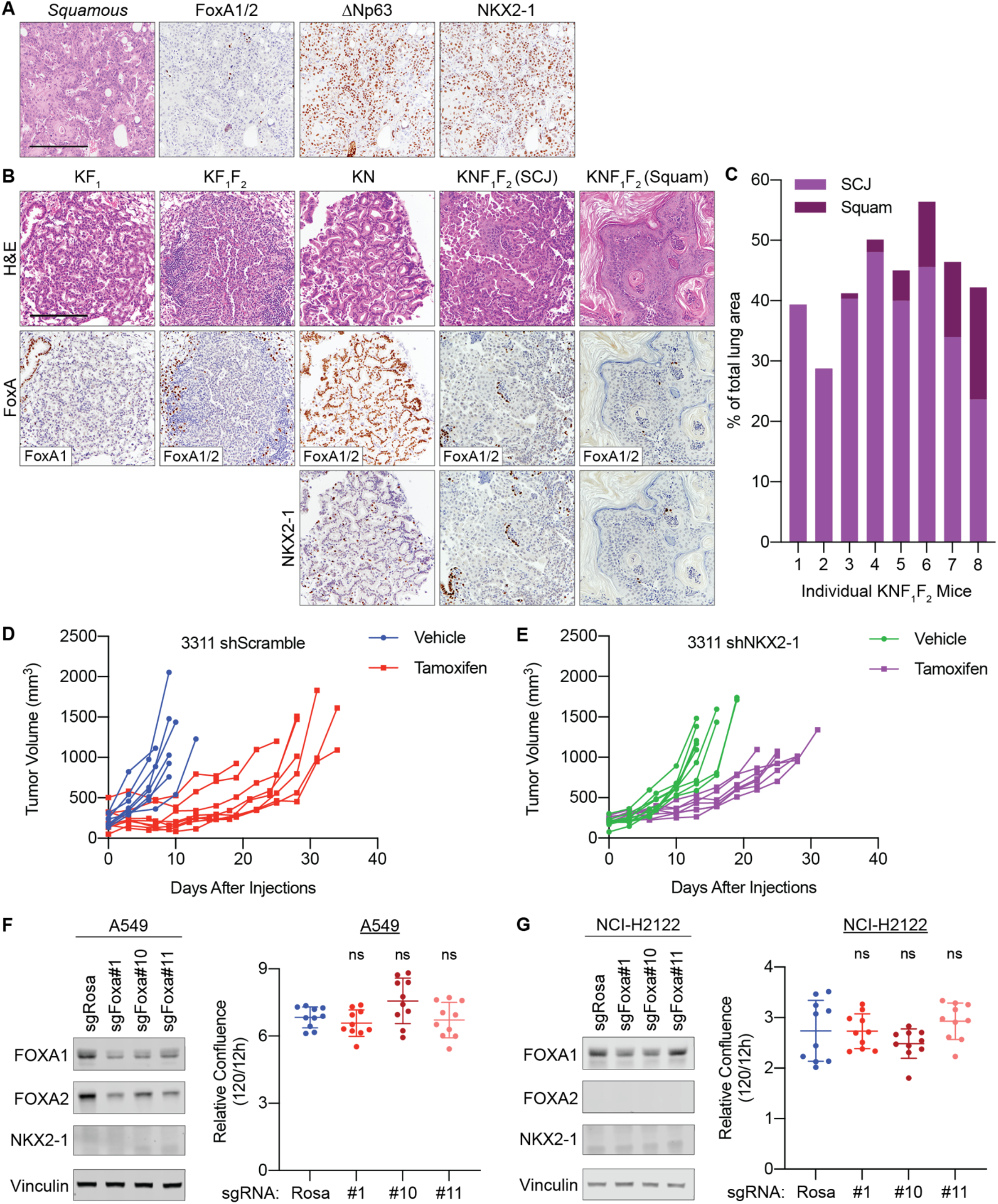
A) Representative images of squamous, βNp63-positive, NKX2-1-positive tumors that emerged in the FoxA1/2-negative cohort of long term KP survival studies (scale bar 500 μm). B) Representative images of KF_1_, KF_1_F_2_, KN, and KNF_1_F_2_ tumors in which alleles were deleted via tamoxifen treatment 6 weeks post-tumor initiation, and tissue was collected 12 weeks post-tumor initiation (SCJ=Squamocolumnar Junction; Squam=Squamous; scale bar 500 μm). C) Percentage of tumor burden in individual KNF_1_F_2_ mice attributable to SCJ and Squamous phenotypes. D) Measurements of subcutaneous tumor volume for individual mice allografted with 3311 shScramble and injected with vehicle or tamoxifen. Each line represents an individual tumor. E) Measurements of subcutaneous tumor volume for individual mice allografted with 3311 shNKX2-1 and injected with vehicle or tamoxifen. Each line represents an individual tumor. F-G) Left: Western blot of protein extract from A549 (F) and NCI-H2122 (G) cell lines transduced with three separate lentivectors that simultaneously express unique sgRNAs targeting both FoxA1 and FoxA2 (sgFoxA#1, sgFoxA#10, and sgFoxA#11), as well as a non-targeting dual sgRNA control (sgNT). Protein collected concurrently with proliferation assay seeding (6 days post-lentiviral sgRNA transduction/2 days post-selection). Right: Relative increase in percent confluence in Incucyte-based growth assay: percent confluence at 120 hours normalized to percent confluence 12 hours post-seeding (n=2 biological replicates with 5 technical replicates each).

## REFERENCES

Adamson, B., Norman, T.M., Jost, M., Cho, M.Y., Nunez, J.K., Chen, Y., Villalta, J.E., Gilbert, L.A., Horlbeck, M.A., Hein, M.Y., et al. (2016). A Multiplexed Single-Cell CRISPR Screening Platform Enables Systematic Dissection of the Unfolded Protein Response. Cell 167, 1867–1882 e1821. 10.1016/j.cell.2016.11.048.

Bachurski, C.J., Yang, G.H., Currier, T.A., Gronostajski, R.M., and Hong, D. (2003). Nuclear factor I/thyroid transcription factor 1 interactions modulate surfactant protein C transcription. Mol Cell Biol 23, 9014–9024. 10.1128/MCB.23.24.9014-9024.2003.

Basseres, D.S., D’Alo, F., Yeap, B.Y., Lowenberg, E.C., Gonzalez, D.A., Yasuda, H., Dayaram, T., Kocher, O.N., Godleski, J.J., Richards, W.G., et al. (2012). Frequent downregulation of the transcription factor Foxa2 in lung cancer through epigenetic silencing. Lung Cancer 77, 31–37. 10.1016/j.lungcan.2012.01.011.

Bejarano, P.A., Baughman, R.P., Biddinger, P.W., Miller, M.A., Fenoglio-Preiser, C., al-Kafaji, B., Di Lauro, R., and Whitsett, J.A. (1996). Surfactant proteins and thyroid transcription factor-1 in pulmonary and breast carcinomas. Mod Pathol 9, 445–452.

Bernardo, G.M., and Keri, R.A. (2012). FOXA1: a transcription factor with parallel functions in development and cancer. Biosci Rep 32, 113–130. 10.1042/BSR20110046.

Besnard, V., Wert, S.E., Hull, W.M., and Whitsett, J.A. (2004). Immunohistochemical localization of Foxa1 and Foxa2 in mouse embryos and adult tissues. Gene Expr Patterns 5, 193–208. 10.1016/j.modgep.2004.08.006.

Bingle, C.D. (1997). Thyroid transcription factor-1. Int J Biochem Cell Biol 29, 1471–1473.

Bochkis, I.M., Schug, J., Ye, D.Z., Kurinna, S., Stratton, S.A., Barton, M.C., and Kaestner, K.H. (2012). Genome-wide location analysis reveals distinct transcriptional circuitry by paralogous regulators Foxa1 and Foxa2. PLoS Genet 8, e1002770. 10.1371/journal.pgen.1002770.

Boggaram, V. (2009). Thyroid transcription factor-1 (TTF-1/Nkx2.1/TITF1) gene regulation in the lung. Clin Sci (Lond) 116, 27–35. 10.1042/CS20080068.

Bohinski, R.J., Di Lauro, R., and Whitsett, J.A. (1994). The lung-specific surfactant protein B gene promoter is a target for thyroid transcription factor 1 and hepatocyte nuclear factor 3, indicating common factors for organ-specific gene expression along the foregut axis. Mol Cell Biol 14, 5671–5681.

Boumahdi, S., and de Sauvage, F.J. (2020). The great escape: tumour cell plasticity in resistance to targeted therapy. Nat Rev Drug Discov 19, 39–56. 10.1038/s41573-019-0044-1.

Bushnell, B. (2021). BBtools.

Camolotto, S.A., Pattabiraman, S., Mosbruger, T.L., Jones, A., Belova, V.K., Orstad, G., Streiff, M., Salmond, L., Stubben, C., Kaestner, K.H., and Snyder, E. (2018). FoxA1 and FoxA2 drive gastric differentiation and suppress squamous identity in NKX2-1-negative lung cancer. Elife 7. 10.7554/eLife.38579.

Campos-Parra, A.D., Aviles, A., Contreras-Reyes, S., Rojas-Marin, C.E., Sanchez-Reyes, R., Borbolla-Escoboza, R.J., and Arrieta, O. (2014). Relevance of the novel IASLC/ATS/ERS classification of lung adenocarcinoma in advanced disease. Eur Respir J 43, 1439–1447. 10.1183/09031936.00138813.

Casanova-Acebes, M., Dalla, E., Leader, A.M., LeBerichel, J., Nikolic, J., Morales, B.M., Brown, M., Chang, C., Troncoso, L., Chen, S.T., et al. (2021). Tissue-resident macrophages provide a pro-tumorigenic niche to early NSCLC cells. Nature 595, 578–584. 10.1038/s41586-021-03651-8.

Caswell, D.R., Chuang, C.-H., Yang, D., Chiou, S.-H., Cheemalavagu, S., Kim-Kiselak, C., Connolly, A., and Winslow, M.M. (2014). Obligate Progression Precedes Lung Adenocarcinoma Dissemination. Cancer Discovery 4, 781–789. 10.1158/2159-8290.cd-13-0862.

Cha, Y.J., and Shim, H.S. (2017). Biology of invasive mucinous adenocarcinoma of the lung. Transl Lung Cancer Res 6, 508–512. 10.21037/tlcr.2017.06.10.

Choi, J., Park, J.E., Tsagkogeorga, G., Yanagita, M., Koo, B.K., Han, N., and Lee, J.H. (2020). Inflammatory Signals Induce AT2 Cell-Derived Damage-Associated Transient Progenitors that Mediate Alveolar Regeneration. Cell Stem Cell 27, 366–382 e367. 10.1016/j.stem.2020.06.020.

Dankort, D., Filenova, E., Collado, M., Serrano, M., Jones, K., and McMahon, M. (2007). A new mouse model to explore the initiation, progression, and therapy of BRAFV600E-induced lung tumors. Genes Dev 21, 379–384. 10.1101/gad.1516407.

Dobin, A., Davis, C.A., Schlesinger, F., Drenkow, J., Zaleski, C., Jha, S., Batut, P., Chaisson, M., and Gingeras, T.R. (2013). STAR: ultrafast universal RNA-seq aligner. Bioinformatics 29, 15–21. 10.1093/bioinformatics/bts635.

Du, Y., Guo, M., Whitsett, J.A., and Xu, Y. (2015). ’LungGENS’: a web-based tool for mapping single-cell gene expression in the developing lung. Thorax 70, 1092–1094. 10.1136/thoraxjnl-2015-207035.

Engblom, C., Pfirschke, C., Zilionis, R., Da Silva Martins, J., Bos, S.A., Courties, G., Rickelt, S., Severe, N., Baryawno, N., Faget, J., et al. (2017). Osteoblasts remotely supply lung tumors with cancer-promoting SiglecF(high) neutrophils. Science 358. 10.1126/science.aal5081.

Feldser, D.M., Kostova, K.K., Winslow, M.M., Taylor, S.E., Cashman, C., Whittaker, C.A., Sanchez-Rivera, F.J., Resnick, R., Bronson, R., Hemann, M.T., and Jacks, T. (2010). Stage-specific sensitivity to p53 restoration during lung cancer progression. Nature 468, 572–575. 10.1038/nature09535.

Gao, B., Xie, W., Wu, X., Wang, L., and Guo, J. (2020). Functionally analyzing the important roles of hepatocyte nuclear factor 3 (FoxA) in tumorigenesis. Biochim Biophys Acta Rev Cancer 1873, 188365. 10.1016/j.bbcan.2020.188365.

Gao, N., LeLay, J., Vatamaniuk, M.Z., Rieck, S., Friedman, J.R., and Kaestner, K.H. (2008). Dynamic regulation of Pdx1 enhancers by Foxa1 and Foxa2 is essential for pancreas development. Genes Dev 22, 3435–3448. 10.1101/gad.1752608.

Han, X., Li, F., Fang, Z., Gao, Y., Li, F., Fang, R., Yao, S., Sun, Y., Li, L., Zhang, W., et al. (2014). Transdifferentiation of lung adenocarcinoma in mice with Lkb1 deficiency to squamous cell carcinoma. Nat Commun 5, 3261. 10.1038/ncomms4261.

Herriges, J.C., Yi, L., Hines, E.A., Harvey, J.F., Xu, G., Gray, P.A., Ma, Q., and Sun, X. (2012). Genome-scale study of transcription factor expression in the branching mouse lung. Dev Dyn 241, 1432–1453. 10.1002/dvdy.23823.

Hight, S.K., Mootz, A., Kollipara, R.K., McMillan, E., Yenerall, P., Otaki, Y., Li, L.S., Avila, K., Peyton, M., Rodriguez-Canales, J., et al. (2020). An in vivo functional genomics screen of nuclear receptors and their co-regulators identifies FOXA1 as an essential gene in lung tumorigenesis. Neoplasia 22, 294–310. 10.1016/j.neo.2020.04.005.

Jin, S., Guerrero-Juarez, C.F., Zhang, L., Chang, I., Ramos, R., Kuan, C.H., Myung, P., Plikus, M.V., and Nie, Q. (2021). Inference and analysis of cell-cell communication using CellChat. Nat Commun 12, 1088. 10.1038/s41467-021-21246-9.

Kaestner, K.H. (2005). The making of the liver: developmental competence in foregut endoderm and induction of the hepatogenic program. Cell Cycle 4, 1146–1148. 10.4161/cc.4.9.2033.

Kaestner, K.H. (2010). The FoxA factors in organogenesis and differentiation. Curr Opin Genet Dev 20, 527–532. 10.1016/j.gde.2010.06.005.

Kobayashi, Y., Tata, A., Konkimalla, A., Katsura, H., Lee, R.F., Ou, J., Banovich, N.E., Kropski, J.A., and Tata, P.R. (2020). Persistence of a regeneration-associated, transitional alveolar epithelial cell state in pulmonary fibrosis. Nat Cell Biol 22, 934–946. 10.1038/s41556-020-0542-8.

LaFave, L.M., Kartha, V.K., Ma, S., Meli, K., Del Priore, I., Lareau, C., Naranjo, S., Westcott, P.M.K., Duarte, F.M., Sankar, V., et al. (2020). Epigenomic State Transitions Characterize Tumor Progression in Mouse Lung Adenocarcinoma. Cancer Cell 38, 212–228 e213. 10.1016/j.ccell.2020.06.006.

Le Magnen, C., Shen, M.M., and Abate-Shen, C. (2018). Lineage Plasticity in Cancer Progression and Treatment. Annu Rev Cancer Biol 2, 271–289. 10.1146/annurev-cancerbio-030617-050224.

Lee, C.L., Moding, E.J., Huang, X., Li, Y., Woodlief, L.Z., Rodrigues, R.C., Ma, Y., and Kirsch, D.G. (2012). Generation of primary tumors with Flp recombinase in FRT-flanked p53 mice. Dis Model Mech 5, 397–402. 10.1242/dmm.009084.

Lee, C.S., Friedman, J.R., Fulmer, J.T., and Kaestner, K.H. (2005). The initiation of liver development is dependent on Foxa transcription factors. Nature 435, 944–947. 10.1038/nature03649.

Li, C.M., Gocheva, V., Oudin, M.J., Bhutkar, A., Wang, S.Y., Date, S.R., Ng, S.R., Whittaker, C.A., Bronson, R.T., Snyder, E.L., et al. (2015). Foxa2 and Cdx2 cooperate with Nkx2-1 to inhibit lung adenocarcinoma metastasis. Genes Dev 29, 1850–1862. 10.1101/gad.267393.115.

Liao, Y., Smyth, G.K., and Shi, W. (2019). The R package Rsubread is easier, faster, cheaper and better for alignment and quantification of RNA sequencing reads. Nucleic Acids Research 47, e47–e47. 10.1093/nar/gkz114.

Little, D.R., Gerner-Mauro, K.N., Flodby, P., Crandall, E.D., Borok, Z., Akiyama, H., Kimura, S., Ostrin, E.J., and Chen, J. (2019). Transcriptional control of lung alveolar type 1 cell development and maintenance by NK homeobox 2-1. Proc Natl Acad Sci U S A 116, 20545–20555. 10.1073/pnas.1906663116.

Little, D.R., Lynch, A.M., Yan, Y., Akiyama, H., Kimura, S., and Chen, J. (2021). Differential chromatin binding of the lung lineage transcription factor NKX2-1 resolves opposing murine alveolar cell fates in vivo. Nat Commun 12, 2509. 10.1038/s41467-021-22817-6.

Liu, C., Glasser, S.W., Wan, H., and Whitsett, J.A. (2002). GATA-6 and thyroid transcription factor-1 directly interact and regulate surfactant protein-C gene expression. J Biol Chem 277, 4519–4525. 10.1074/jbc.M107585200.

Love, M.I., Huber, W., and Anders, S. (2014). Moderated estimation of fold change and dispersion for RNA-seq data with DESeq2. Genome Biol 15, 550. 10.1186/s13059-014-0550-8.

Maier, B., Leader, A.M., Chen, S.T., Tung, N., Chang, C., LeBerichel, J., Chudnovskiy, A., Maskey, S., Walker, L., Finnigan, J.P., et al. (2020). A conserved dendritic-cell regulatory program limits antitumour immunity. Nature 580, 257–262. 10.1038/s41586-020-2134-y.

Marjanovic, N.D., Hofree, M., Chan, J.E., Canner, D., Wu, K., Trakala, M., Hartmann, G.G., Smith, O.C., Kim, J.Y., Evans, K.V., et al. (2020). Emergence of a High-Plasticity Cell State during Lung Cancer Evolution. Cancer Cell 38, 229–246 e213. 10.1016/j.ccell.2020.06.012.

Meza, R., Meernik, C., Jeon, J., and Cote, M.L. (2015). Lung cancer incidence trends by gender, race and histology in the United States, 1973-2010. PLoS One 10, e0121323. 10.1371/journal.pone.0121323.

Minoo, P., Hu, L., Xing, Y., Zhu, N.L., Chen, H., Li, M., Borok, Z., and Li, C. (2007). Physical and functional interactions between homeodomain NKX2.1 and winged helix/forkhead FOXA1 in lung epithelial cells. Mol Cell Biol 27, 2155–2165. 10.1128/MCB.01133-06.

Miyoshi, H., and Stappenbeck, T.S. (2013). In vitro expansion and genetic modification of gastrointestinal stem cells in spheroid culture. Nature Protocols 8, 2471–2482. 10.1038/nprot.2013.153.

Mollaoglu, G., Guthrie, M.R., Bohm, S., Bragelmann, J., Can, I., Ballieu, P.M., Marx, A., George, J., Heinen, C., Chalishazar, M.D., et al. (2017). MYC Drives Progression of Small Cell Lung Cancer to a Variant Neuroendocrine Subtype with Vulnerability to Aurora Kinase Inhibition. Cancer Cell 31, 270–285. 10.1016/j.ccell.2016.12.005.

Mollaoglu, G., Jones, A., Wait, S.J., Mukhopadhyay, A., Jeong, S., Arya, R., Camolotto, S.A., Mosbruger, T.L., Stubben, C.J., Conley, C.J., et al. (2018). The Lineage-Defining Transcription Factors SOX2 and NKX2-1 Determine Lung Cancer Cell Fate and Shape the Tumor Immune Microenvironment. Immunity 49, 764–779 e769. 10.1016/j.immuni.2018.09.020.

Moses, M.A., George, A.L., Sakakibara, N., Mahmood, K., Ponnamperuma, R.M., King, K.E., and Weinberg, W.C. (2019). Molecular Mechanisms of p63-Mediated Squamous Cancer Pathogenesis. Int J Mol Sci 20. 10.3390/ijms20143590.

Niederst, M.J., Sequist, L.V., Poirier, J.T., Mermel, C.H., Lockerman, E.L., Garcia, A.R., Katayama, R., Costa, C., Ross, K.N., Moran, T., et al. (2015). RB loss in resistant EGFR mutant lung adenocarcinomas that transform to small-cell lung cancer. Nat Commun 6, 6377. 10.1038/ncomms7377.

Paranjapye, A., Mutolo, M.J., Ebron, J.S., Leir, S.H., and Harris, A. (2020). The FOXA1 transcriptional network coordinates key functions of primary human airway epithelial cells. Am J Physiol Lung Cell Mol Physiol 319, L126–l136. 10.1152/ajplung.00023.2020.

Quintanal-Villalonga, A., Chan, J.M., Yu, H.A., Pe’er, D., Sawyers, C.L., Sen, T., and Rudin, C.M. (2020). Lineage plasticity in cancer: a shared pathway of therapeutic resistance. Nat Rev Clin Oncol 17, 360–371. 10.1038/s41571-020-0340-z.

Robinson, J.L., Holmes, K.A., and Carroll, J.S. (2013). FOXA1 mutations in hormone-dependent cancers. Front Oncol 3, 20. 10.3389/fonc.2013.00020.

Russell, P.A.W., Z.; Wright, G.M.; Daniels, M.; Conron, M.; Williams, R.A. (2011). Does lung adenocarcinoma subtype predict patient survival?: A clinicopathologic study based on the new International Association for the Study of Lung Cnacer/American Thoracic Society/European Respiratory Society international multidisciplinary lung adenocarcinoma classification. Journal of thoracic oncology: official publication of the International Association for the Study of Lung Cnacer, 1496–1504.

Samarasinghe, K.T.G., and Crews, C.M. (2021). Targeted protein degradation: a promise for undruggable proteins. Cell Chem Biol. 10.1016/j.chembiol.2021.04.011.

Schonhuber, N., Seidler, B., Schuck, K., Veltkamp, C., Schachtler, C., Zukowska, M., Eser, S., Feyerabend, T.B., Paul, M.C., Eser, P., et al. (2014). A next-generation dual-recombinase system for time- and host-specific targeting of pancreatic cancer. Nat Med 20, 1340–1347. 10.1038/nm.3646.

Shai, A., Dankort, D., Juan, J., Green, S., and McMahon, M. (2015). TP53 Silencing Bypasses Growth Arrest of BRAFV600E-Induced Lung Tumor Cells in a Two-Switch Model of Lung Tumorigenesis. Cancer Res 75, 3167–3180. 10.1158/0008-5472.CAN-14-3701.

Shi, J., Wang, E., Milazzo, J.P., Wang, Z., Kinney, J.B., and Vakoc, C.R. (2015). Discovery of cancer drug targets by CRISPR-Cas9 screening of protein domains. Nat Biotechnol 33, 661–667. 10.1038/nbt.3235.

Snyder, E.L., Watanabe, H., Magendantz, M., Hoersch, S., Chen, T.A., Wang, D.G., Crowley, D., Whittaker, C.A., Meyerson, M., Kimura, S., and Jacks, T. (2013). Nkx2-1 represses a latent gastric differentiation program in lung adenocarcinoma. Molecular cell 50, 185–199. 10.1016/j.molcel.2013.02.018.

Strunz, M., Simon, L.M., Ansari, M., Kathiriya, J.J., Angelidis, I., Mayr, C.H., Tsidiridis, G., Lange, M., Mattner, L.F., Yee, M., et al. (2020). Alveolar regeneration through a Krt8+ transitional stem cell state that persists in human lung fibrosis. Nat Commun 11, 3559. 10.1038/s41467-020-17358-3.

Sun, Z., and Yang, P. (2006). Gene expression profiling on lung cancer outcome prediction: present clinical value and future premise. Cancer Epidemiol Biomarkers Prev 15, 2063–2068. 10.1158/1055-9965.EPI-06-0505.

Sund, N.J., Ang, S.L., Sackett, S.D., Shen, W., Daigle, N., Magnuson, M.A., and Kaestner, K.H. (2000). Hepatocyte nuclear factor 3beta (Foxa2) is dispensable for maintaining the differentiated state of the adult hepatocyte. Mol Cell Biol 20, 5175–5183.

Tavernari, D., Battistello, E., Dheilly, E., Petruzzella, A.S., Mina, M., Sordet-Dessimoz, J., Peters, S., Krueger, T., Gfeller, D., Riggi, N., et al. (2021). Nongenetic Evolution Drives Lung Adenocarcinoma Spatial Heterogeneity and Progression. Cancer Discov 11, 1490–1507. 10.1158/2159-8290.CD-20-1274.

Teng, M., Zhou, S., Cai, C., Lupien, M., and He, H.H. (2021). Pioneer of prostate cancer: past, present and the future of FOXA1. Protein & Cell 12, 29–38. 10.1007/s13238-020-00786-8.

Travaglini, K.J., Nabhan, A.N., Penland, L., Sinha, R., Gillich, A., Sit, R.V., Chang, S., Conley, S.D., Mori, Y., Seita, J., et al. (2020). A molecular cell atlas of the human lung from single-cell RNA sequencing. Nature 587, 619–625. 10.1038/s41586-020-2922-4.

Travis, W.D., Brambilla, E., Noguchi, M., Nicholson, A.G., Geisinger, K.R., Yatabe, Y., Beer, D.G., Powell, C.A., Riely, G.J., Van Schil, P.E., et al. (2011). International association for the study of lung cancer/american thoracic society/european respiratory society international multidisciplinary classification of lung adenocarcinoma. J Thorac Oncol 6, 244–285. 10.1097/JTO.0b013e318206a221.

Tu, S., Li, M., Chen, H., Tan, F., Xu, J., Waxman, D.J., Zhang, Y., and Shao, Z. (2021). MAnorm2 for quantitatively comparing groups of ChIP-seq samples. Genome Res 31, 131–145. 10.1101/gr.262675.120.

Unni, A.M., Lockwood, W.W., Zejnullahu, K., Lee-Lin, S.Q., and Varmus, H. (2015). Evidence that synthetic lethality underlies the mutual exclusivity of oncogenic KRAS and EGFR mutations in lung adenocarcinoma. Elife 4, e06907. 10.7554/eLife.06907.

Varma, S., Cao, Y., Tagne, J.B., Lakshminarayanan, M., Li, J., Friedman, T.B., Morell, R.J., Warburton, D., Kotton, D.N., and Ramirez, M.I. (2012). The transcription factors Grainyhead-like 2 and NK2-homeobox 1 form a regulatory loop that coordinates lung epithelial cell morphogenesis and differentiation. J Biol Chem 287, 37282–37295. 10.1074/jbc.M112.408401.

Wan, H., Dingle, S., Xu, Y., Besnard, V., Kaestner, K.H., Ang, S.L., Wert, S., Stahlman, M.T., and Whitsett, J.A. (2005). Compensatory roles of Foxa1 and Foxa2 during lung morphogenesis. J Biol Chem 280, 13809–13816. 10.1074/jbc.M414122200.

Wang, L., Lu, Q., Gao, W., and Yu, S. (2021). Recent advancement on development of drug-induced macrophage polarization in control of human diseases. Life Sci 284, 119914. 10.1016/j.lfs.2021.119914.

Watanabe, H., Francis, J.M., Woo, M.S., Etemad, B., Lin, W., Fries, D.F., Peng, S., Snyder, E.L., Tata, P.R., Izzo, F., et al. (2013). Integrated cistromic and expression analysis of amplified NKX2-1 in lung adenocarcinoma identifies LMO3 as a functional transcriptional target. Genes Dev 27, 197–210. 10.1101/gad.203208.112.

Weir, B.A., Woo, M.S., Getz, G., Perner, S., Ding, L., Beroukhim, R., Lin, W.M., Province, M.A., Kraja, A., Johnson, L.A., et al. (2007). Characterizing the cancer genome in lung adenocarcinoma. Nature 450, 893–898. 10.1038/nature06358.

Winslow, M.M., Dayton, T.L., Verhaak, R.G., Kim-Kiselak, C., Snyder, E.L., Feldser, D.M., Hubbard, D.D., DuPage, M.J., Whittaker, C.A., Hoersch, S., et al. (2011). Suppression of lung adenocarcinoma progression by Nkx2-1. Nature 473, 101–104. 10.1038/nature09881.

Yamaguchi, T., Yanagisawa, K., Sugiyama, R., Hosono, Y., Shimada, Y., Arima, C., Kato, S., Tomida, S., Suzuki, M., Osada, H., and Takahashi, T. (2012). NKX2-1/TITF1/TTF-1-Induced ROR1 is required to sustain EGFR survival signaling in lung adenocarcinoma. Cancer Cell 21, 348–361. 10.1016/j.ccr.2012.02.008.

Yang, D., Jones, M.G., Naranjo, S., Rideout, W.M., Min, K.H., Ho, R., Wu, W., Replogle, J.M., Page, J.L., Quinn, J.J., et al. (2021). Lineage Recording Reveals the Phylodynamics, Plasticity and Paths of Tumor Evolution. bioRxiv, 2021.2010.2012.464111. 10.1101/2021.10.12.464111.

Yi, M., Tong, G.X., Murry, B., and Mendelson, C.R. (2002). Role of CBP/p300 and SRC-1 in transcriptional regulation of the pulmonary surfactant protein-A (SP-A) gene by thyroid transcription factor-1 (TTF-1). J Biol Chem 277, 2997–3005. 10.1074/jbc.M109793200.

Young, N.P., Crowley, D., and Jacks, T. (2011). Uncoupling cancer mutations reveals critical timing of p53 loss in sarcomagenesis. Cancer Res 71, 4040–4047. 10.1158/0008-5472.CAN-10-4563.

Yu, G., Wang, L.G., and He, Q.Y. (2015). ChIPseeker: an R/Bioconductor package for ChIP peak annotation, comparison and visualization. Bioinformatics 31, 2382–2383. 10.1093/bioinformatics/btv145.

Yuan, A., Hsiao, Y.J., Chen, H.Y., Chen, H.W., Ho, C.C., Chen, Y.Y., Liu, Y.C., Hong, T.H., Yu, S.L., Chen, J.J., and Yang, P.C. (2015). Opposite Effects of M1 and M2 Macrophage Subtypes on Lung Cancer Progression. Sci Rep 5, 14273. 10.1038/srep14273.

Zewdu, R., Mehrabad, E.M., Ingram, K., Fang, P., Gillis, K.L., Camolotto, S.A., Orstad, G., Jones, A., Mendoza, M.C., Spike, B.T., and Snyder, E.L. (2021). An NKX2-1/ERK/WNT feedback loop modulates gastric identity and response to targeted therapy in lung adenocarcinoma. Elife 10. 10.7554/eLife.66788.

Zilionis, R., Engblom, C., Pfirschke, C., Savova, V., Zemmour, D., Saatcioglu, H.D., Krishnan, I., Maroni, G., Meyerovitz, C.V., Kerwin, C.M., et al. (2019). Single-Cell Transcriptomics of Human and Mouse Lung Cancers Reveals Conserved Myeloid Populations across Individuals and Species. Immunity 50, 1317–1334 e1310. 10.1016/j.immuni.2019.03.009.

